# Ultrasmall chemogenetic tags with group-transfer ligands

**DOI:** 10.1101/2025.05.10.653252

**Authors:** Sreekanth Vedagopuram, Shaimaa Sindi, Santosh K. Chaudhary, Rajaiah Pergu, Prashant Singh, Endri Karaj, Jeffrey E. Fung, Arghya Deb, Stephan J. DeCarlo, Jaron A. M. Mercer, Kei Yamada, Diego Rodriguez, David R. Liu, Amit Choudhary

## Abstract

Chemogenetic tags are valuable tools for studying functions of a given protein-of-interest (POI) lacking small-molecule ligands, but most tags are too large for several POIs. Here, we report two ultrasmall chemogenetic tags (mgTag and cTag) of 36 and 50 amino acids (aa) that, to the best of our knowledge, are the smallest reported. These tags exhibit *transferase*-type reactivity with their ligands to append any moiety of interest to the tag. cTag utilizes an engineered C1 domain-bearing cysteine that undergoes group-transfer reaction with its ligand. Likewise, mgTag utilizes an engineered zinc-finger domain-bearing cysteine that undergoes group-transfer reaction with its molecular-glue ligand in the presence of cereblon (CRBN). We applied these tags in the context of cell signaling and proximity induction. While the fusion of HaloTag (297 aa) to the KRAS^G12D^ (188 aa) disrupted its ability to activate the growth-signaling pathway, fusion of mgTag or cTag did not. Group-transfer of BRD4 binder to tags appended to Abelson kinase (ABL) induced proximity between ABL and BRD4, resulting in the latter’s phosphorylation. Deletion of the *transferase*-type reactivity reduced phosphorylation levels, suggesting that proximity-inducing chimeras with group-transfer design may be more efficacious in certain scenarios. We envision these ultrasmall tags to have wide-ranging applications, including in basic science, biotechnology, and medicine.

**TOC:** 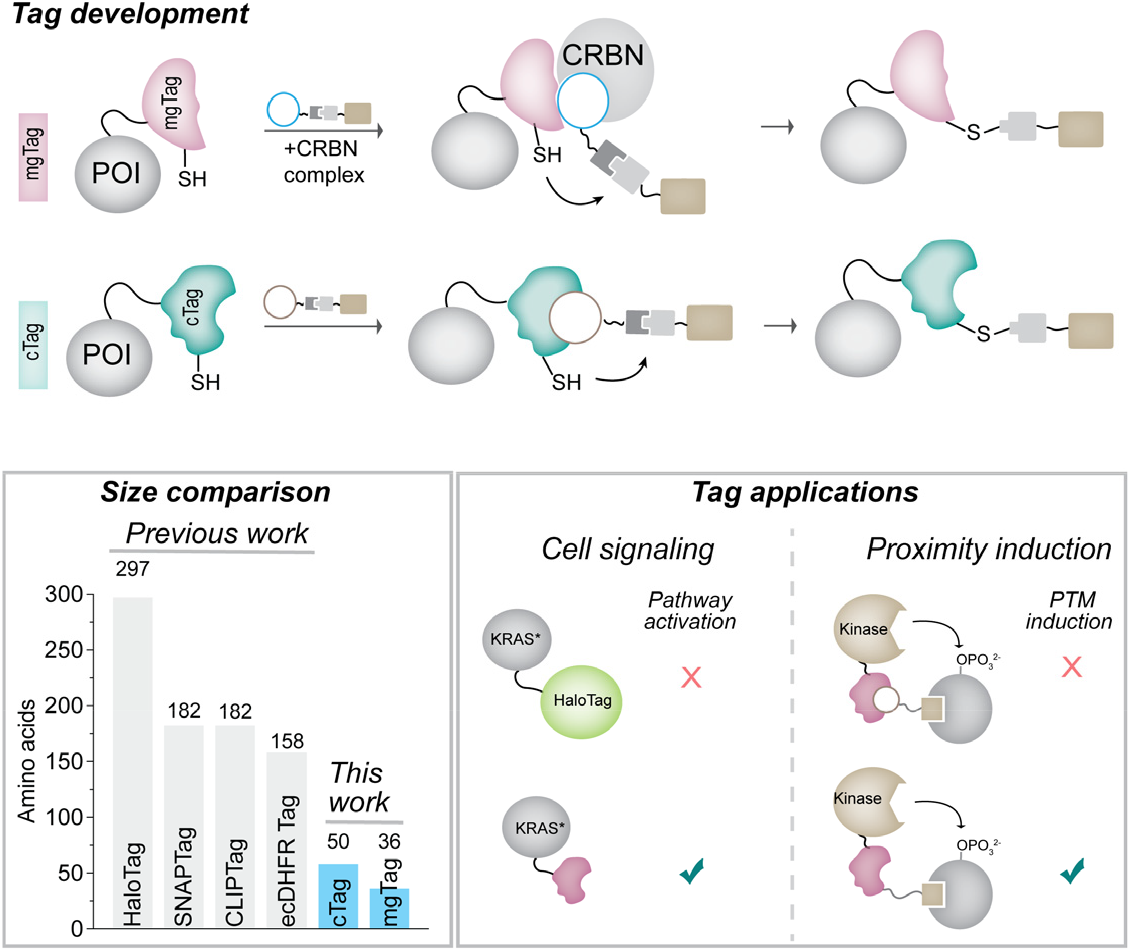

---

Chemogenetic protein tags and their small-molecule ligands are essential research tools ^1^ as only ≈10 % of the proteome has small-molecule ligands, most of which are enzyme inhibitors.^2^ While extant covalent chemogenetic tags (e.g., HaloTag, SNAP-Tag, Figure 1A)^3-8^ have been immensely impactful, emergent applications require significantly smaller tags with drug-like ligands. For example, studies on several high-therapeutic value signaling proteins that lack ligands (e.g., transcription factors) can benefit from tags, but these proteins are nearly the same size as contemporary tags (∼33 kDa, Figure 1A) and function *via* packed protein complexes that contemporary tags will disrupt.^9-16^ CRISPR-based technologies are advancing towards efficient genome-wide, pooled knock-in screens, but contemporary tags are too large for both the technology and several protein-of-interest (POI).^17-24^ While contemporary tags have enabled rapid evaluation of the efficacy of proximity induction between POI and large ubiquitin-ligase complexes for POI degradation, emergent chimeras for other post-translational modifications (e.g., phosphorylation, acetylation) involve smaller-sized complexes that require smaller tags.^25-29^ Finally, while contemporary tags are of non-human origin, several applications of chemogenetic tags (e.g., in CAR T cell engineering) require sequences of human origin to avert immunogenicity.^30-36^ Herein, we report two chemogenetic tags of 36 or 50 amino acids (and their drug-like ligands) that are smaller than any reported covalent chemogenetic tags to the best of our knowledge (Figure 1B, C). Both tags are of human origin and compatible with small-sized signaling proteins or protein-editing technologies.

**Figure 1.**
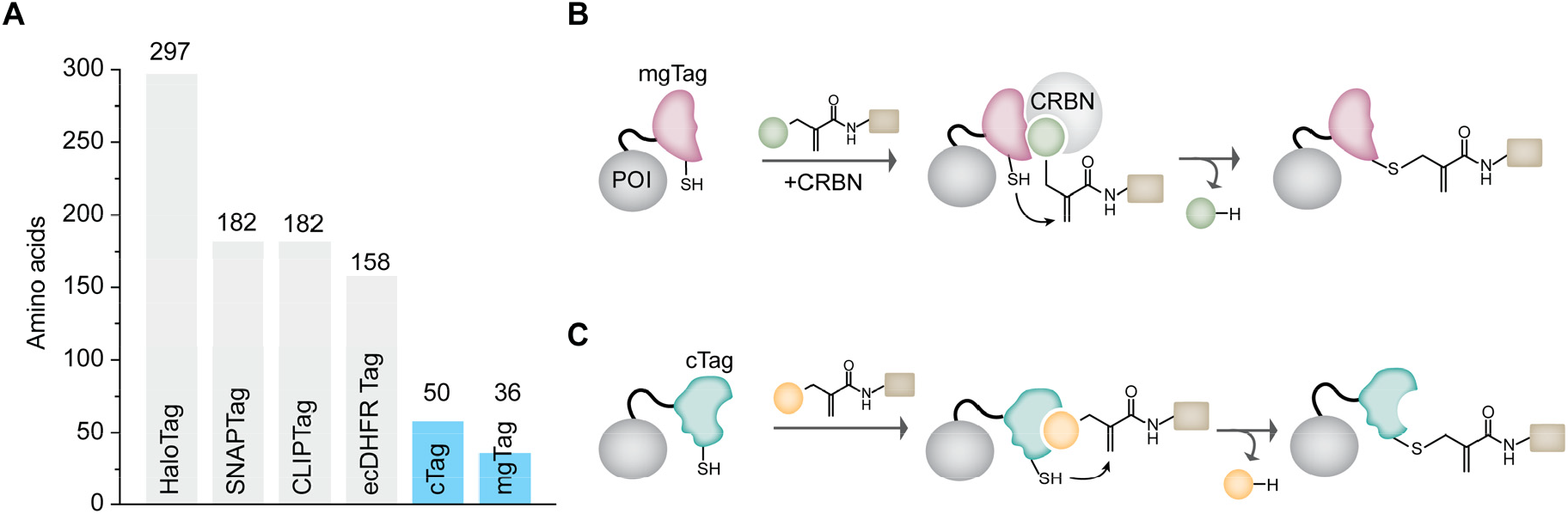
(**A**) Size of the existing covalent chemogenetic tags. (**B**) A molecular-glue Tag (mgTag) based on ternary-complex guided group-transfer labeling. (**C**) A C1 domain tag (cTag) based on binary complex guided group-transfer labeling.

We recently reported a molecular glue PT-179 that induces proximity between cereblon (CRBN), the substrate receptor of the CRL4^CRBN^ E3-based ubiquitin ligase, and SD40, a 36 amino acid tag that can be fused to POI.^37,38^ We and others have also reported group-transfer linkers that use the protein’s ligand to append a group to the cysteine or lysine side chain with concomitant ligand release.^39-50^ Group-transfer reactions, which are reminiscent of transferases (e.g., kinases), uncouple the pharmacology of the group being transferred (e.g., phosphoryl for kinases) from the ligand (e.g., ATP for kinases). We envisioned engineering cysteine on the SD40 tag, which, in the presence of a CRBN complex and a group-transfer probe (PT-179 with a group-transfer linker), will undergo a *transferase*-type reaction with the release of PT-179, thereby affording a miniature molecular glue tag (mgTag, Figure 1B).

Guided by the reported ternary complex structure of SD40:PT-179:CRBN complex (Figure 2A),^38^ we engineered cysteine at various positions on SD40 proximal to PT-179 to generate six constructs (v1-6) with nanoluciferase (NLuc) as POI (Figure 2B). In parallel, we appended various groups (e.g., alkyne handle **1** or a fluorophore **2**) to PT-179 via the group-transfer methacrylamide linker (Figure 2C).^43,47^ We confirmed that **1** could undergo group transfer with *N*-Ac-cysteine-OMe with a rate constant of 5.7 × 10^−5^ s^-1^ (Figure 2D, E), in agreement with previous reports on similar systems.^43^ Using compound **2**, which has fluorophore as the transferred group (Figure 2C), we screened six constructs in cells for labeling using in-gel fluorescence (Figure 2F). We observed the labeling of only the v2 construct (hereafter named as mgTag) and confirmed that this labeling was dose-dependent and detectable at even at 10 nM (Figure 2G). To confirm that this labeling was POI-independent, we fused mgTag to oncogenic *KRAS*^*G12D*^ and observed similar dose-dependent labeling (Figure 2H).

**Figure 2.**
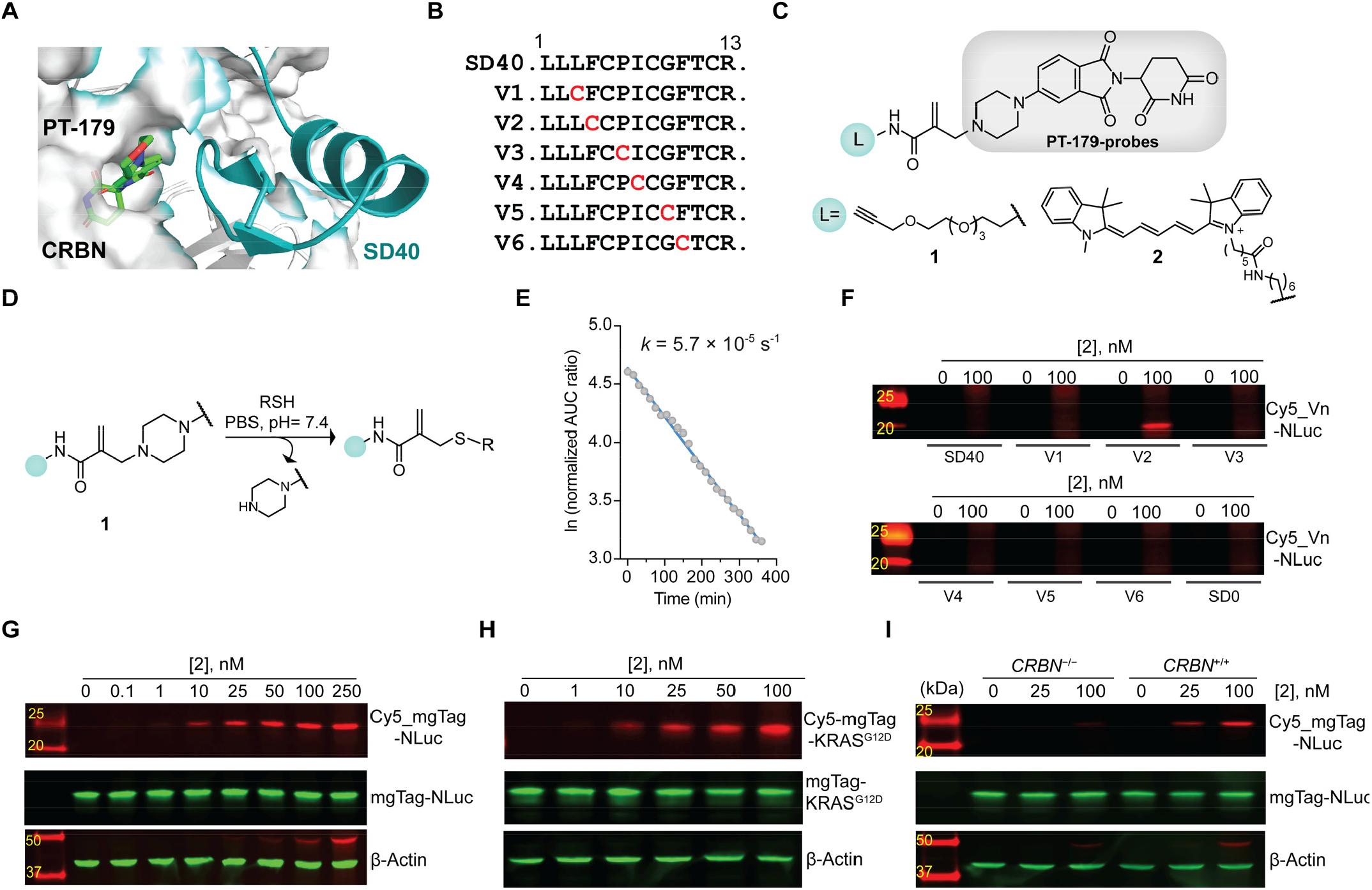
(**A**) Cryo-EM structure of SD40, PT179, and CRBN ternary complex PDB: 8TNQ. (**B**) Cysteine tiling on SD40 at sites proximal to PT-179. (**C**) Structures of PT-179 based probes with alkyne (**1**) or fluorophore (**2**) as the transferred group. (**D-E**) Biochemical validation of the group-transfer reactivity of **1** with cysteine. (**F**) In-gel fluorescence-based screening of cysteine-engineered SD40 constructs (v1-6) using **2** (0 nM left lane, 100 nM right lane). SD0 refers to the initial sequence used for the evolution of SD40.^38^ (**G**) Dose-dependent labeling of mgTag.NLuc with **2**. (**H**) Dose-dependent labeling of mgTag.KRAS^G12D^ with **2 (I**) Labeling studies of **2** in *CRBN*^-/-^ or *CRBN*^+/+^ cell lines.

Furthermore, to confirm that the group-transfer labeling was operating by molecular-glue mechanism, we utilized a CRBN double knock-out cell line (*CRBN*^-/-^)^51^ and confirmed that labeling occurred only in *CRBN*^+/+^ line but not *CRBN*^-/-^ (Figure 2I). These studies validate that mgTag can be used without degradation of POI.

Next, we ventured to employ group-transfer chemistry to generate a miniature tag from the C1 domain. Protein Kinase C (PKC) is activated by ligands such as the phorbol ester TPA that possess a polar headgroup and a lipid tail; the polar headgroup binds to the C1 domain of PKC while the lipid tail facilitates PKC’s membrane translocation (Figure 3A).^52-54^ Since C1 domains are small (∼50 amino acids), we hypothesized that a miniature tag could be generated by grafting cysteine on the C1 domain and placing an appropriate group-transfer linker on the headgroup of the PKC ligand. We initiated our studies by determining if the C1 domain can fold and bind to the PKC ligands without requiring other PKC domains. Taking advantage of its miniature size, we chemically synthesized the C1 domain peptide and used circular dichroism to confirm the presence of the folded state─the circular dichroism spectra showed peaks at 208 and 222 nm (Figure 3B), indicative of a folded peptide. Since the C1 domain contains two zinc finger-like motifs, removing zinc ions from the buffer resulted in spectra resembling random coil. (Figure 3B). Next, we confirmed that the ligand binding pocket was preserved by quantifying the binding of PKC ligand, phorbol ester. We expressed and purified the C1 domain fused to maltose-binding protein (MBP) and tested the binding using microscale thermophoresis (MST). Gratifyingly, we observed a dose-dependent shift in fluorescence, indicating a *K*_d_ of 190 nM (Figure 3C). These studies confirmed that the C1 domain, when isolated from PKC and fused to another protein, maintains its ability to fold and bind to PKC ligands.

**Figure 3.**
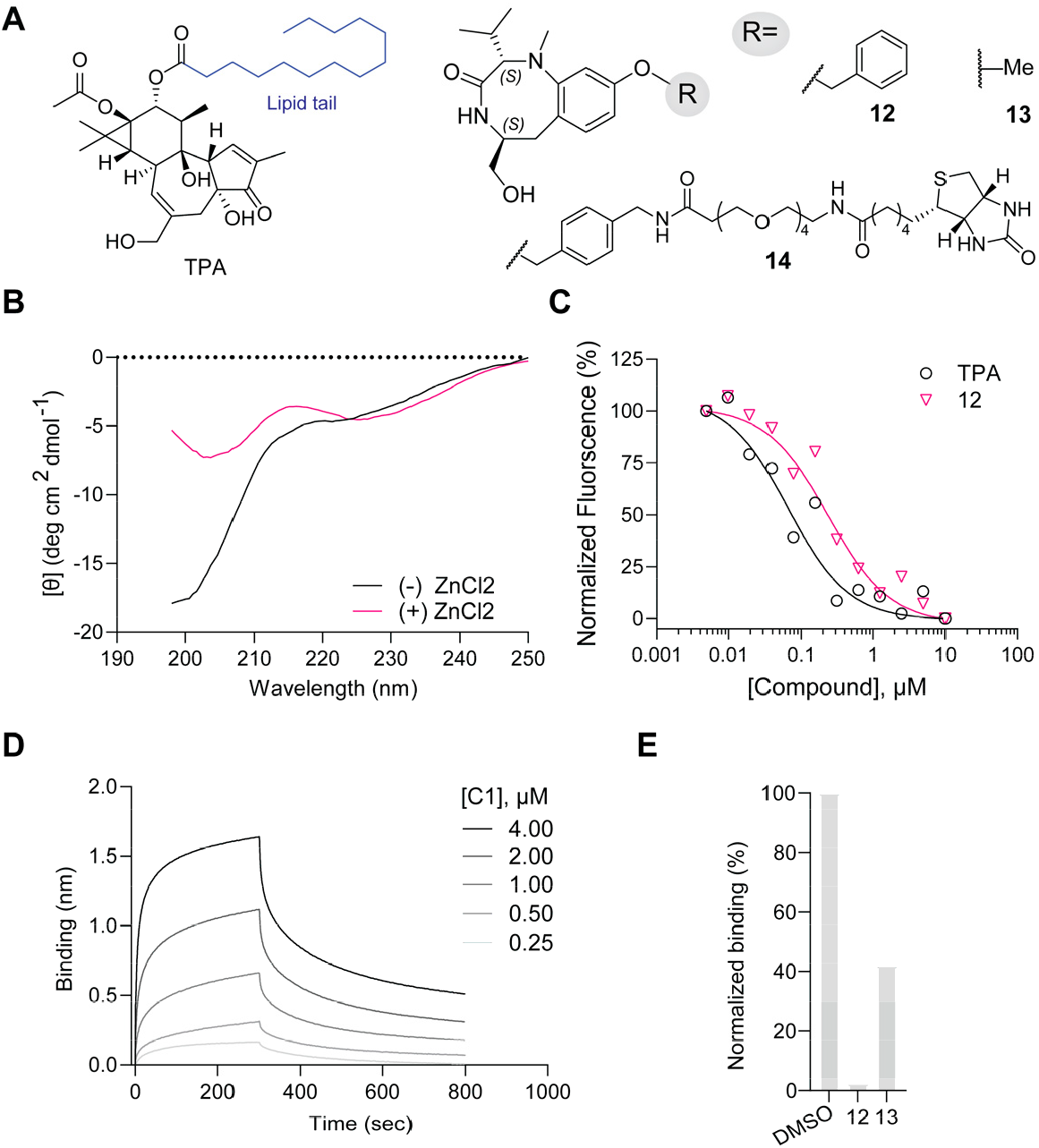
(**A**) Structures of TPA and benzolactam probes (**12, 13** and **14**). (**B**) Circular dichroism spectra of C1 domain in presence and absence of zinc ions. **(C)** Microscale thermophoresis for TPA and **12. (D)** BLI sensograms for dose-dependent binding of **14. (E)** Competition-based BLI experiment for **12** and **13**.

**Scheme 1.**
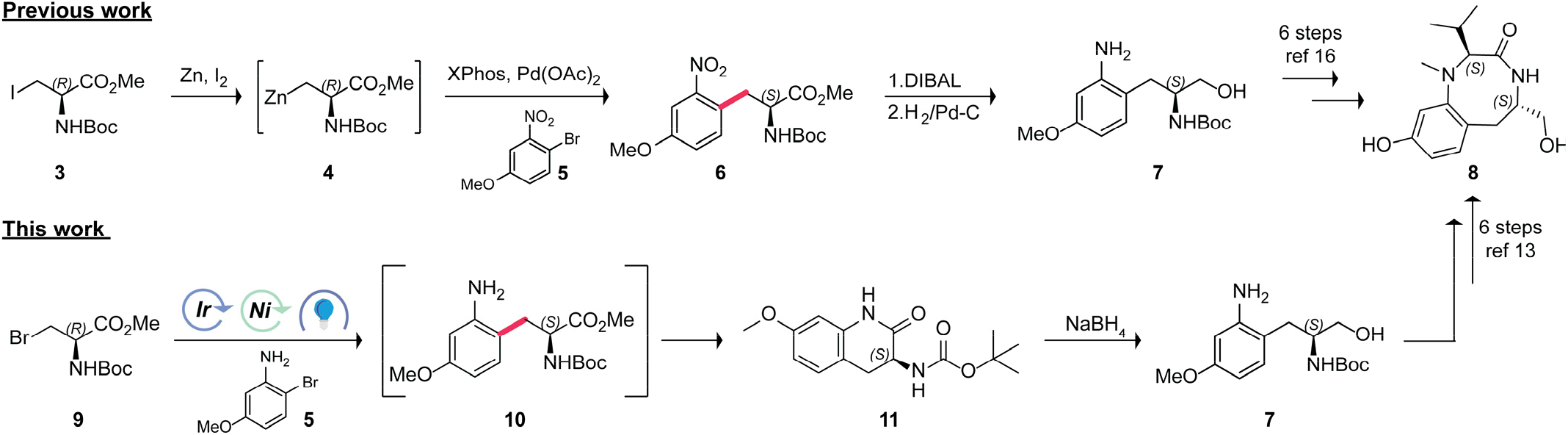
Improved synthetic route to access benzolactam scaffold.

The phorbol ester head group, while a potent PKC binder, is challenging to synthesize,^55^ preventing deep structure-activity relationship studies necessary to identify the group-transfer linker attachment site. We focused our efforts on the benzolactam scaffold, a potent PKC binder that is synthetically more tractable, although current approaches still require 10 steps.^56^ We further improved the reported synthesis by replacing the laborious and non-scalable Negishi coupling of compounds **3** and **5** with a photochemical reaction catalyzed by iridium and nickel, providing key intermediate benzolactam in 8 steps (Scheme 1). In this optimized approach, intermediate **9** is directly coupled with **5** using visible-light photoredox catalysis,^57,58^ providing **10** that cyclizes to **11** during the removal of volatiles. **11** was reduced to **7**, which was converted to benzolactam intermediate **8** following our previous report.^56^ Next, we generated benzolactam analog **12** (Figure 3A) and tested binding to the C1.MBP fusion using MST and obtained a *K*_d_ of 230 nM that is comparable to that of phorbol ester TPA. We confirmed binding in an orthogonal assay using biolayer interferometry (BLI). Here, we immobilized a biotinylated analog of **12** (compound **14**) on the tip of the BLI sensor and assessed its binding with the C1 domain in the solution. We observed a dose-dependent increase in the signal, confirming binding (Figure 3D). We implemented this assay in a competition format where **12** was pre-incubated with the C1 domain and observed a decrease in BLI signal by 41-fold (Figure 3E and Figure S1A). We explored analogs of **12** lacking benzyl group (compound **13**, Figure 3A) to reduce the size, but this analog decreased signal by only ∼3-fold, suggesting that removing the benzyl group causes substantial loss of binding (Figure 3E and Figure S1B). We further confirmed this observation using MST (Figure S1C). These studies established a highly yielding route to access the benzolactam scaffold and allowed us to identify a minimalist binder of the C1 domain.

Following successful binding studies, we grafted cysteines around the benzolactam binding pocket (Figure 4A, S1D) in an approach mimicking mgTag—the cysteines were placed within the pocket (marked in grey) and on the surface (marked in green). We also generated a library of benzolactam analogs bearing thiol-reactive warheads (e.g., acrylamides, epoxides, haloacetamides, Figure S1E) at sites that will be proximal to the engineered cysteines and studied labeling using intact mass (Figures S2-5). Intriguingly, for cysteines grafted at the base of the pocket, we did not observe labeling with any of the thiol-reactive warheads on the benzolactam scaffold (Figures S2-5). Our efforts to enhance the cysteine reactivity by placing a proximal histidine also were unsuccessful (Figure S1D and S2-5). We hypothesize that these mutations potentially perturb the binding pocket and/or zinc finger.

**Figure 4.**
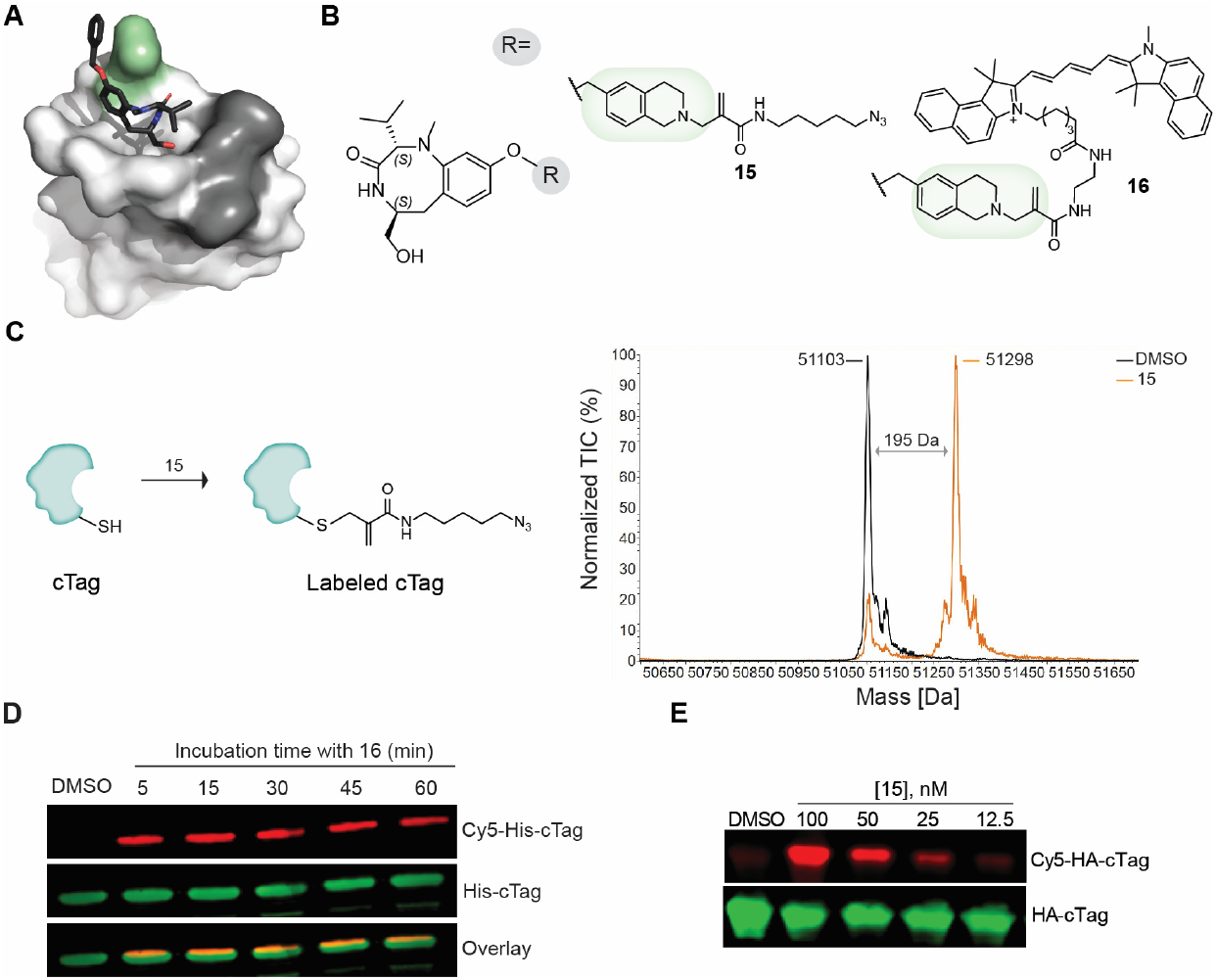
(**A**) Crystal structure of C1 domain with **4**. Colored regions point to sites of cysteine engineering. **(B)** Structures for cTag probes **15** and **16 (C)** Schematic of biochemical labeling of cTag by probe **15** (l*eft*) and Intact mass analysis of cTag treated with DMSO and compound **15** (*right*). The theoretical mass adduct of labeled cTag is +194 Da. **(D)** Kinetic analysis of cTag labeling by **16**. (**E**) Dose-dependent cellular labeling of cTag by **15**, monitored *via* click reaction with DBCO-Cy5.

We focused our efforts on engineered cysteine on the surface (green site, Figure 4A) and were gratified to observe group-transfer reactivity with methacrylamide **15** (Figure 4B) akin to mgTag above and yielding a C1 domain tag (cTag, Figure 4C). To confirm that cTag reactivity is proximity-driven, we replaced the critical benzyl group with a piperidyl group (Figure S6A) and observed no labeling. Furthermore, we observed fast-labeling kinetics of cTag with **16**, a hallmark of increased effective molarity owing to proximity induction between cysteine and the reactive group-**16** attained saturating labeling within 5 minutes at low micromolar protein and ligand concentrations (Figure 4D). Using **15** (labels cTag with an azide) and click chemistry with DBCO-fluorophore,^59^ we confirmed labeling in cells at as low as 25 nM (Figure 4E). By clicking DBCO-biotin, we successfully pulldown MBP fused to cTag from the lysate (Figure S6B). To confirm that our cTag ligand was not activating PKC, we used PKC motif antibody and immunoblotting to assess proteome-wide changes in the phosphorylation. Gratifyingly, we did not observe significant changes in the phosphorylation status compared to DMSO at the concentrations used for the labeling studies (Figure S6C). Collectively, these studies identify a miniature tag based on an engineered C1 domain that can undergo group-transfer chemistry with a benzolactam scaffold.

We next examined the impact of tag size on the signaling of oncogenic KRAS^G12D^, which, upon N-terminal myristoylation, translocates to the membrane to form nanoclusters and activate the mitogen activation protein kinase (MAPK) pathway.^13-16^ We hypothesize that contemporary tags (e.g., HaloTag, 33kDa) that are larger than KRAS (21 kDa) and myristoyl group will perturb the ability of KRAS to form clusters and downstream signaling. To test this hypothesis, we fused HaloTag, mgTag, *or cTag* to the *N*-terminus of *KRAS*^*G12D*^ and assessed MAPK pathway activation by measuring levels of phosphorylated ERK. While cells with transient expression of KRAS^G12D^ or mgTag /cTag fused KRAS^G12D^ activated the MAPK pathway, cells expression HaloTag.KRAS^G12D^ fusion did not exhibit MAPK pathway activation (Figure 5A). These studies suggest that mgTag and cTag do not perturb the native function of a small signaling protein.

**Figure 5.**
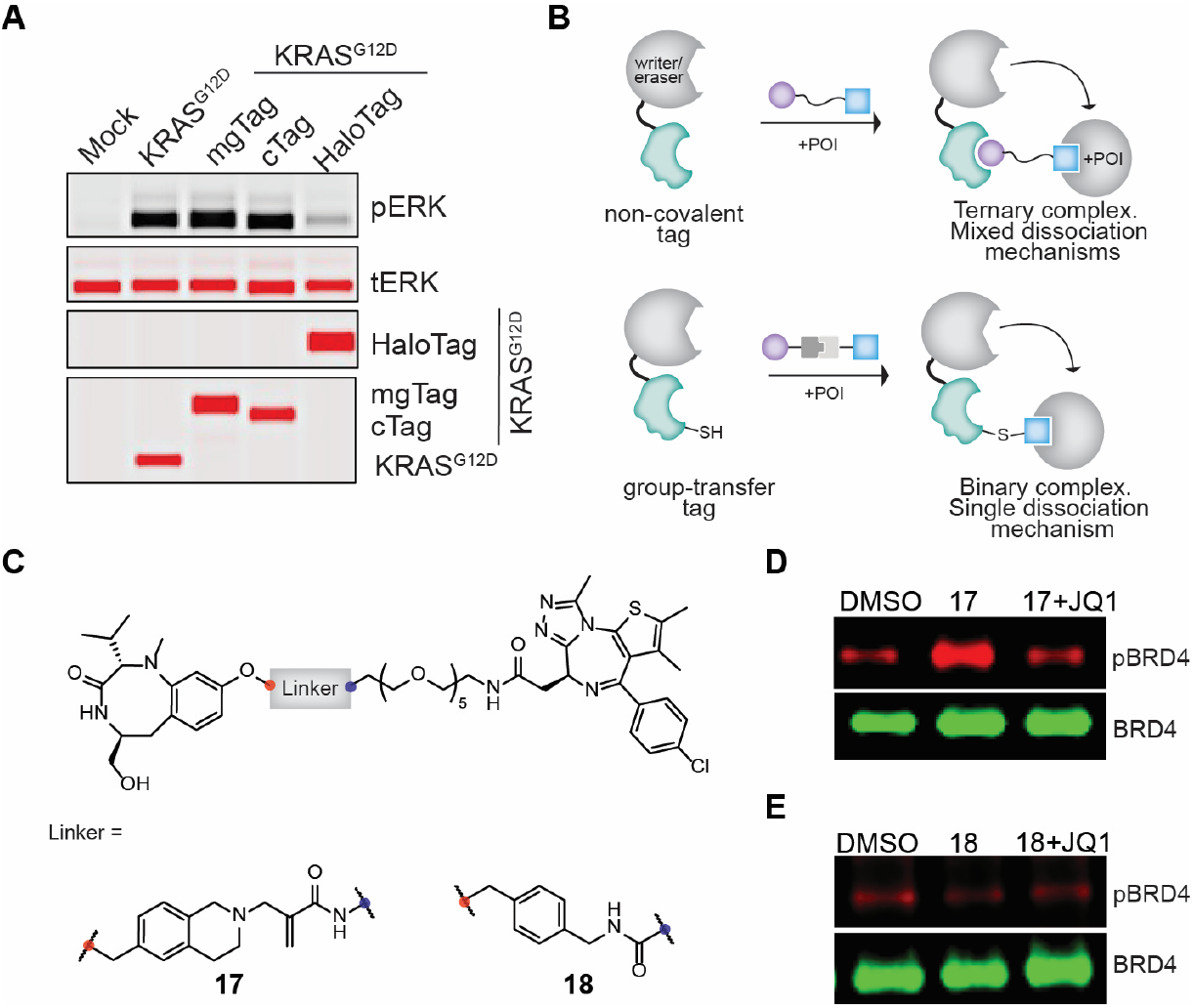
**(A)** Downstream MAPK signaling (measured via pERK) of mgTag, cTag and HaloTag-fused KRAS^G12D^. **(B)** Schematic of protein editing by recruiting PTM writers or erasers with non-covalent (*top*) or group-transfer tags (*bottom*). (**C**) Structures of group-transfer (**17**) and non-covalent (**18**) cTag-JQ1 probes for targeted BRD4 phosphorylation. **(D-E)** In-cellular phosphorylation of BRD4 with cTag or mgTag fused Abl utilizing compounds **17 (D)** and **18** (**E**) respectively.

Next, we explored the compatibility of mgTag and cTag with induced-proximity modalities for appending smaller post-translational modifications. We have previously reported that chimeric small molecules induce POI phosphorylation by bringing POI in proximity to a kinase. Since many kinases lack appropriate recruiting ligands, chemogenetic tags can be impactful. In fact, chemogenetic tags have allowed systematic evaluation of various ubiquitin ligases lacking ligands.^60^ We fused ABL kinase with either mgTag or cTag and synthesized probes which contain JQ1 that binds to BRD4,^61^ to induce proximity between ABL and BRD4 (Figures 5B, C, and S6D). Gratifyingly, we observed increased BRD4 phosphorylation with cTag probe **17**, which was competed out by an excess of JQ1 (Figure 5D, S6D for mgTag). Intriguingly, compound **18**, which lacks a group-transfer reactive group, was unable to induce BRD4 phosphorylation at 250 nM (the concentration used for **17**) but was able to do so at 10 µM,^56^ suggesting a 40-fold increase in potency (Figure 5E). These studies suggest that covalent recruitment of PTM writers or eraser enzymes is more efficient than non-covalent recruitment. Importantly, these studies confirm that mgTag and cTag can be used for the development of induced-proximity modalities.

We describe two miniature chemogenetic tags and their drug-like ligands that, to the best of our knowledge, are the smallest known covalent tags. The ligands of these tags exhibit a transferase-like reactivity and, unlike many covalent ligands, are neutral from a translational entropy standpoint. Covalent ligands operate by an associative mechanism, but these group-transfer ligands operate by an associative-dissociative mechanism, which results in the same number of species after the reaction. These tags do not perturb the activity of even small signaling proteins (21 kDa) and can be used to evaluate emergent-induced proximity modalities further. Furthermore, tags of similar length have been successfully used with CRISPR-based knock-in technologies,^62^ and these studies will advance efforts to develop genome-wide knock-in screens. Employment of bump-hole and directed evolution strategies can yield orthogonal mgTag and cTags, which may find applications in synthetic biology and cell engineering. Thus, we envision the widespread application of these tags in basic research, biomedicine, and biotechnology.

## ACKNOWLEDGMENTS

This work was supported by DARPA (HR00112120010) and NIH (1U01DK137242, R01GM137606, R01DK129464, R01GM132825).

## AUTHOR CONTRIBUTIONS

S.V, S.S, S.K.C, R.P, P.S, and E.K contributed equally to this work.

## REFERENCES

(1) Pick a Tag and Explore the Functions of Your Pet Protein. Vandemoortele, G.; Eyckerman, S.; Gevaert, K. Trends in Biotechnology 2019, 37, 1078–1090

(2) Unexplored therapeutic opportunities in the human genome. Oprea, T. I.; Bologa, C. G.; Brunak, S.; Campbell, A.; Gan, G. N.; Gaulton, A.; Gomez, S. M.; Guha, R.; Hersey, A.; Holmes, J.; Jadhav, A.; Jensen, L. J.; Johnson, G. L.; Karlson, A.; Leach, A. R.; Ma’ayan, A.; Malovannaya, A.; Mani, S.; Mathias, S. L.; McManus, M. T.; Meehan, T. F.; von Mering, C.; Muthas, D.; Nguyen, D.-T.; Overington, J. P.; Papadatos, G.; Qin, J.; Reich, C.; Roth, B. L.; Schürer, S. C.; Simeonov, A.; Sklar, L. A.; Southall, N.; Tomita, S.; Tudose, I.; Ursu, O.; Vidović, D.; Waller, A.; Westergaard, D.; Yang, J. J.; Zahoránszky-Köhalmi, G. Nature Reviews Drug Discovery 2018, 17, 317-332.PMC6339563

(3) A general method for the covalent labeling of fusion proteins with small molecules in vivo. Keppler, A.; Gendreizig, S.; Gronemeyer, T.; Pick, H.; Vogel, H.; Johnsson, K. Nat Biotechnol 2003, 21, 86–9

(4) Specific labeling of cell surface proteins with chemically diverse compounds. George, N.; Pick, H.; Vogel, H.; Johnsson, N.; Johnsson, K. J Am Chem Soc 2004, 126, 8896–7

(5) Multicolor imaging of cell surface proteins. Vivero-Pol, L.; George, N.; Krumm, H.; Johnsson, K.; Johnsson, N. J Am Chem Soc 2005, 127, 12770–1

(6) An engineered protein tag for multiprotein labeling in living cells. Gautier, A.; Juillerat, A.; Heinis, C.; Corrêa, I. R., Jr.; Kindermann, M.; Beaufils, F.; Johnsson, K. Chem Biol 2008, 15, 128–36

(7) HaloTag: A Novel Protein Labeling Technology for Cell Imaging and Protein Analysis. Los, G. V.; Encell, L. P.; McDougall, M. G.; Hartzell, D. D.; Karassina, N.; Zimprich, C.; Wood, M. G.; Learish, R.; Ohana, R. F.; Urh, M.; Simpson, D.; Mendez, J.; Zimmerman, K.; Otto, P.; Vidugiris, G.; Zhu, J.; Darzins, A.; Klaubert, D. H.; Bulleit, R. F.; Wood, K. V. ACS Chem. Biol. 2008, 3, 373–382

(8) Second-generation covalent TMP-tag for live cell imaging. Chen, Z.; Jing, C.; Gallagher, S. S.; Sheetz, M. P.; Cornish, V. W. J Am Chem Soc 2012, 134, 13692-9.PMC3433398

(9) Spatial organization and signal transduction at intercellular junctions. Manz, B. N.; Groves, J. T. Nature Reviews Molecular Cell Biology 2010, 11, 342–352

(10) Nanoclustering as a dominant feature of plasma membrane organization. Garcia-Parajo, M. F.; Cambi, A.; Torreno-Pina, J. A.; Thompson, N.; Jacobson, K. J Cell Sci 2014, 127, 4995-5005.PMC4260763

(11) Biomolecular condensates: organizers of cellular biochemistry. Banani, S. F.; Lee, H. O.; Hyman, A. A.; Rosen, M. K. Nature Reviews Molecular Cell Biology 2017, 18, 285–298

(12) Advances in targeting ‘undruggable’ transcription factors with small molecules. Henley, M. J.; Koehler, A. N. Nat Rev Drug Discov 2021, 20, 669–688

(13) A structural model of a Ras-Raf signalosome. Mysore, V. P.; Zhou, Z. W.; Ambrogio, C.; Li, L.; Kapp, J. N.; Lu, C.; Wang, Q.; Tucker, M. R.; Okoro, J. J.; Nagy-Davidescu, G.; Bai, X.; Plückthun, A.; Jänne, P. A.; Westover, K. D.; Shan, Y.; Shaw, D. E. Nat Struct Mol Biol 2021, 28, 847-857.PMC8643099

(14) KRAS interaction with RAF1 RAS-binding domain and cysteine-rich domain provides insights into RAS-mediated RAF activation. Tran, T. H.; Chan, A. H.; Young, L. C.; Bindu, L.; Neale, C.; Messing, S.; Dharmaiah, S.; Taylor, T.; Denson, J.-P.; Esposito, D.; Nissley, D. V.; Stephen, A. G.; McCormick, F.; Simanshu, D. K. Nature Communications 2021, 12, 1176

(15) Targeting KRAS in cancer. Singhal, A.; Li, B. T.; O’Reilly, E. M. Nature Medicine 2024, 30, 969–983

(16) Plasma membrane-associated ARAF condensates fuel RAS-related cancer drug resistance. Li, W.; Shi, X.; Tan, C.; Jiang, Z.; Li, M.; Ji, Z.; Zhou, J.; Luo, M.; Fan, Z.; Ding, Z.; Fang, Y.; Sun, J.; Ding, J.; Lu, H.; Ma, W.; Xie, W.; Su, W. Nat Chem Biol 2025

(17) Efficient and flexible tagging of endogenous genes by homology-independent intron targeting. Serebrenik, Y. V.; Sansbury, S. E.; Kumar, S. S.; Henao-Mejia, J.; Shalem, O. Genome Res 2019, 29, 1322-1328.PMC6673721

(18) Pooled tagging and hydrophobic targeting of endogenous proteins for unbiased mapping of unfolded protein responses. Sansbury, S. E.; Serebrenik, Y. V.; Lapidot, T.; Burslem, G. M.; Shalem, O. bioRxiv 2023, 2023.07.13.548611

(19) High-throughput PRIME-editing screens identify functional DNA variants in the human genome. Ren, X.; Yang, H.; Nierenberg, J. L.; Sun, Y.; Chen, J.; Beaman, C.; Pham, T.; Nobuhara, M.; Takagi, M. A.; Narayan, V.; Li, Y.; Ziv, E.; Shen, Y. Mol Cell 2023, 83, 4633-4645.e9.PMC10766087

(20) High-throughput evaluation of genetic variants with prime editing sensor libraries. Gould, S. I.; Wuest, A. N.; Dong, K.; Johnson, G. A.; Hsu, A.; Narendra, V. K.; Atwa, O.; Levine, S. S.; Liu, D. R.; Sánchez Rivera, F. J. Nat Biotechnol 2024

(21) High-throughput screening of human genetic variants by pooled prime editing. Herger, M.; Kajba, C. M.; Buckley, M.; Cunha, A.; Strom, M.; Findlay, G. M. bioRxiv 2024, 2024.04.01.587366

(22) High-throughput optimized prime editing mediated endogenous protein tagging for pooled imaging of protein localization. Sanchez, H. M.; Lapidot, T.; Shalem, O. bioRxiv 2024, 2024.09.16.613361

(23) Pooled endogenous protein tagging and recruitment for systematic profiling of protein function. Serebrenik, Y. V.; Mani, D.; Maujean, T.; Burslem, G. M.; Shalem, O. Cell Genom 2024, 4, 100651.PMC11602618

(24) A benchmarked, high-efficiency prime editing platform for multiplexed dropout screening. Cirincione, A.; Simpson, D.; Yan, W.; McNulty, R.; Ravisankar, P.; Solley, S. C.; Yan, J.; Lim, F.; Farley, E. K.; Singh, M.; Adamson, B. Nat Methods 2025, 22, 92-101.PMC11725502

(25) Chemical tags for labeling proteins inside living cells. Jing, C.; Cornish, V. W. Acc Chem Res 2011, 44, 784-92.PMC3232020

(26) HaloPROTACS: Use of Small Molecule PROTACs to Induce Degradation of HaloTag Fusion Proteins. Buckley, D. L.; Raina, K.; Darricarrere, N.; Hines, J.; Gustafson, J. L.; Smith, I. E.; Miah, A. H.; Harling, J. D.; Crews, C. M. ACS Chem Biol 2015, 10, 1831-7.PMC4629848

(27) Proximity-Based Modalities for Biology and Medicine. Liu, X.; Ciulli, A. ACS Cent Sci 2023, 9, 1269-1284.PMC10375889

(28) Proximity-inducing modalities: the past, present, and future. Singh, S.; Tian, W.; Severance, Z. C.; Chaudhary, S. K.; Anokhina, V.; Mondal, B.; Pergu, R.; Singh, P.; Dhawa, U.; Singha, S.; Choudhary, A. Chemical Society Reviews 2023, 52, 5485-5515.37477631

(29) Chemogenetic Tools in Focus: Proximity, Conformation, and Sterics. Shen, J.; Zhou, G.; Wang, W. Chemistry–Methods 2024, 4, e202300051

(30) Remote control of therapeutic T cells through a small molecule-gated chimeric receptor. Wu, C. Y.; Roybal, K. T.; Puchner, E. M.; Onuffer, J.; Lim, W. A. Science 2015, 350, aab4077.PMC4721629

(31) Human antibody-based chemically induced dimerizers for cell therapeutic applications. Hill, Z. B.; Martinko, A. J.; Nguyen, D. P.; Wells, J. A. Nat Chem Biol 2018, 14, 112-117.PMC6352901

(32) Sensitive and adaptable pharmacological control of CAR T cells through extracellular receptor dimerization. Leung, W. H.; Gay, J.; Martin, U.; Garrett, T. E.; Horton, H. M.; Certo, M. T.; Blazar, B. R.; Morgan, R. A.; Gregory, P. D.; Jarjour, J.; Astrakhan, A. JCI Insight 2019, 5.PMC6629089

(33) A computationally designed chimeric antigen receptor provides a small-molecule safety switch for T-cell therapy. Giordano-Attianese, G.; Gainza, P.; Gray-Gaillard, E.; Cribioli, E.; Shui, S.; Kim, S.; Kwak, M. J.; Vollers, S.; Corria Osorio, A. J.; Reichenbach, P.; Bonet, J.; Oh, B. H.; Irving, M.; Coukos, G.; Correia, B. E. Nat Biotechnol 2020, 38, 426–432

(34) A conformation-specific ON-switch for controlling CAR T cells with an orally available drug. Zajc, C. U.; Dobersberger, M.; Schaffner, I.; Mlynek, G.; Pühringer, D.; Salzer, B.; Djinović-Carugo, K.; Steinberger, P.; De Sousa Linhares, A.; Yang, N. J.; Obinger, C.; Holter, W.; Traxlmayr, M. W.; Lehner, M. Proc Natl Acad Sci U S A 2020, 117, 14926-14935.PMC7334647

(35) Reversible ON- and OFF-switch chimeric antigen receptors controlled by lenalidomide. Jan, M.; Scarfò, I.; Larson, R. C.; Walker, A.; Schmidts, A.; Guirguis, A. A.; Gasser, J. A.; Słabicki, M.; Bouffard, A. A.; Castano, A. P.; Kann, M. C.; Cabral, M. L.; Tepper, A.; Grinshpun, D. E.; Sperling, A. S.; Kyung, T.; Sievers, Q. L.; Birnbaum, M. E.; Maus, M. V.; Ebert, B. L. Sci Transl Med 2021, 13.PMC8045771

(36) An IMiD-inducible degron provides reversible regulation for chimeric antigen receptor expression and activity. Carbonneau, S.; Sharma, S.; Peng, L.; Rajan, V.; Hainzl, D.; Henault, M.; Yang, C.; Hale, J.; Shulok, J.; Tallarico, J.; Porter, J.; Brogdon, J. L.; Dranoff, G.; Bradner, J. E.; Hild, M.; Guimaraes, C. P. Cell Chem Biol 2021, 28, 583

(37) Proteolysis-targeting chimeras with reduced off-targets. Nguyen, T. M.; Sreekanth, V.; Deb, A.; Kokkonda, P.; Tiwari, P. K.; Donovan, K. A.; Shoba, V.; Chaudhary, S. K.; Mercer, J. A. M.; Lai, S.; Sadagopan, A.; Jan, M.; Fischer, E. S.; Liu, D. R.; Ebert, B. L.; Choudhary, A. Nat Chem 2024, 16, 218-228.PMC10913580

(38) Continuous evolution of compact protein degradation tags regulated by selective molecular glues. Mercer, J. A. M.; DeCarlo, S. J.; Roy Burman, S. S.; Sreekanth, V.; Nelson, A. T.; Hunkeler, M.; Chen, P. J.; Donovan, K. A.; Kokkonda, P.; Tiwari, P. K.; Shoba, V. M.; Deb, A.; Choudhary, A.; Fischer, E. S.; Liu, D. R. Science 2024, 383, eadk4422

(39) 6-Substituted-4-(3-bromophenylamino)quinazolines as putative irreversible inhibitors of the epidermal growth factor receptor (EGFR) and human epidermal growth factor receptor (HER-2) tyrosine kinases with enhanced antitumor activity. Tsou, H. R.; Mamuya, N.; Johnson, B. D.; Reich, M. F.; Gruber, B. C.; Ye, F.; Nilakantan, R.; Shen, R.; Discafani, C.; DeBlanc, R.; Davis, R.; Koehn, F. E.; Greenberger, L. M.; Wang, Y. F.; Wissner, A. J Med Chem 2001, 44, 2719–34

(40) Structure- and reactivity-based development of covalent inhibitors of the activating and gatekeeper mutant forms of the epidermal growth factor receptor (EGFR). Ward, R. A.; Anderton, M. J.; Ashton, S.; Bethel, P. A.; Box, M.; Butterworth, S.; Colclough, N.; Chorley, C. G.; Chuaqui, C.; Cross, D. A.; Dakin, L. A.; Debreczeni, J.; Eberlein, C.; Finlay, M. R.; Hill, G. B.; Grist, M.; Klinowska, T. C.; Lane, C.; Martin, S.; Orme, J. P.; Smith, P.; Wang, F.; Waring, M. J. J Med Chem 2013, 56, 7025–48

(41) Rapid labelling and covalent inhibition of intracellular native proteins using ligand-directed N-acyl-N-alkyl sulfonamide. Tamura, T.; Ueda, T.; Goto, T.; Tsukidate, T.; Shapira, Y.; Nishikawa, Y.; Fujisawa, A.; Hamachi, I. Nature Communications 2018, 9, 1870.PMC5951806

(42) A programmable chemical switch based on triggerable Michael acceptors. Zhuang, J.; Zhao, B.; Meng, X.; Schiffman, J. D.; Perry, S. L.; Vachet, R. W.; Thayumanavan, S. Chemical Science 2020, 11, 2103–2111

(43) Systematic Study of the Glutathione Reactivity of N-Phenylacrylamides: 2. Effects of Acrylamide Substitution. Birkholz, A.; Kopecky, D. J.; Volak, L. P.; Bartberger, M. D.; Chen, Y.; Tegley, C. M.; Arvedson, T.; McCarter, J. D.; Fotsch, C.; Cee, V. J. J Med Chem 2020, 63, 11602–11614

(44) Enhanced Suppression of a Protein–Protein Interaction in Cells Using Small-Molecule Covalent Inhibitors Based on an N-Acyl-N-alkyl Sulfonamide Warhead. Ueda, T.; Tamura, T.; Kawano, M.; Shiono, K.; Hobor, F.; Wilson, A. J.; Hamachi, I. Journal of the American Chemical Society 2021, 143, 4766–4774

(45) Tunable Methacrylamides for Covalent Ligand Directed Release Chemistry. Reddi, R. N.; Resnick, E.; Rogel, A.; Rao, B. V.; Gabizon, R.; Goldenberg, K.; Gurwicz, N.; Zaidman, D.; Plotnikov, A.; Barr, H.; Shulman, Z.; London, N. J Am Chem Soc 2021, 143, 4979-4992.PMC8041284

(46) Site-Specific Labeling of Endogenous Proteins Using CoLDR Chemistry. Reddi, R. N.; Rogel, A.; Resnick, E.; Gabizon, R.; Prasad, P. K.; Gurwicz, N.; Barr, H.; Shulman, Z.; London, N. Journal of the American Chemical Society 2021, 143, 20095–20108

(47) Development and Applications of Chimera Platforms for Tyrosine Phosphorylation. Pergu, R.; Shoba, V. M.; Chaudhary, S. K.; Munkanatta Godage, D. N. P.; Deb, A.; Singha, S.; Dhawa, U.; Singh, P.; Anokhina, V.; Singh, S.; Siriwardena, S. U.; Choudhary, A. ACS Central Science 2023, 9, 1558-1566.PMC10450875

(48) Lysine-Reactive N-Acyl-N-aryl Sulfonamide Warheads: Improved Reaction Properties and Application in the Covalent Inhibition of an Ibrutinib-Resistant BTK Mutant. Kawano, M.; Murakawa, S.; Higashiguchi, K.; Matsuda, K.; Tamura, T.; Hamachi, I. J Am Chem Soc 2023, 145, 26202–26212

(49) Targeted Covalent Modification Strategies for Drugging the Undruggable Targets. Tamura, T.; Kawano, M.; Hamachi, I. Chemical Reviews 2025

(50) Covalent Proximity Inducers. London, N. Chemical Reviews 2025, 125, 326–368

(51) Thalidomide promotes degradation of SALL4, a transcription factor implicated in Duane Radial Ray syndrome. Donovan, K. A.; An, J.; Nowak, R. P.; Yuan, J. C.; Fink, E. C.; Berry, B. C.; Ebert, B. L.; Fischer, E. S. Elife 2018, 7.PMC6156078

(52) Mechanism of Interaction of Protein Kinase C with Phorbol Esters: REVERSIBILITY AND NATURE OF MEMBRANE ASSOCIATION (*). Mosior, M.; Newton, A. C. Journal of Biological Chemistry 1995, 270, 25526–25533

(53) C1 domains: structure and ligand-binding properties. Das, J.; Rahman, G. M. Chem Rev 2014, 114, 12108–31

(54) Structural anatomy of Protein Kinase C C1 domain interactions with diacylglycerol and other agonists. Katti, S. S.; Krieger, I. V.; Ann, J.; Lee, J.; Sacchettini, J. C.; Igumenova, T. I. Nature Communications 2022, 13, 2695

(55) Nineteen-step total synthesis of (+)-phorbol. Kawamura, S.; Chu, H.; Felding, J.; Baran, P. S. Nature 2016, 532, 90–93

(56) Synthetic Reprogramming of Kinases Expands Cellular Activities of Proteins. Shoba, V. M.; Munkanatta Godage, D. N. P.; Chaudhary, S. K.; Deb, A.; Siriwardena, S. U.; Choudhary, A. Angew Chem Int Ed Engl 2022, 61, e202202770.PMC9527066

(57) Photosensitized, energy transfer-mediated organometallic catalysis through electronically excited nickel(II). Welin, E. R.; Le, C.; Arias-Rotondo, D. M.; McCusker, J. K.; MacMillan, D. W. C. Science 2017, 355, 380–385

(58) Synthesis of Enantiopure Unnatural Amino Acids by Metallaphotoredox Catalysis. Faraggi, T. M.; Rouget-Virbel, C.; Rincón, J. A.; Barberis, M.; Mateos, C.; García-Cerrada, S.; Agejas, J.; de Frutos, O.; MacMillan, D. W. C. Organic Process Research & Development 2021, 25, 1966–1973

(59) A Strain-Promoted [3 + 2] Azide−Alkyne Cycloaddition for Covalent Modification of Biomolecules in Living Systems. Agard, N. J.; Prescher, J. A.; Bertozzi, C. R. Journal of the American Chemical Society 2004, 126, 15046–15047

(60) Electrophilic PROTACs that degrade nuclear proteins by engaging DCAF16. Zhang, X.; Crowley, V. M.; Wucherpfennig, T. G.; Dix, M. M.; Cravatt, B. F. Nat Chem Biol 2019, 15, 737-746.PMC6592777

(61) Selective inhibition of BET bromodomains. Filippakopoulos, P.; Qi, J.; Picaud, S.; Shen, Y.; Smith, W. B.; Fedorov, O.; Morse, E. M.; Keates, T.; Hickman, T. T.; Felletar, I.; Philpott, M.; Munro, S.; McKeown, M. R.; Wang, Y.; Christie, A. L.; West, N.; Cameron, M. J.; Schwartz, B.; Heightman, T. D.; La Thangue, N.; French, C. A.; Wiest, O.; Kung, A. L.; Knapp, S.; Bradner, J. E. Nature 2010, 468, 1067-73.PMC3010259

(62) Programmable deletion, replacement, integration and inversion of large DNA sequences with twin prime editing. Anzalone, A. V.; Gao, X. D.; Podracky, C. J.; Nelson, A. T.; Koblan, L. W.; Raguram, A.; Levy, J. M.; Mercer, J. A. M.; Liu, D. R. Nat Biotechnol 2022, 40, 731-740.PMC9117393

