## Supporting Information for "Ultrasmall chemogenetic tags with group-transfer ligands"

**1. Biological Materials and methods.**

**1. 1. Materials and general procedures.**

1.1.1. Plasmids

All constructs were cloned in-house, and sequences are provided at 1.17.

1.1.2. Antibodies

Phospho-Tyrosine (P-Tyr-1000) MultiMab® (cat# 8954), Ha-Tag (6E2) (cat# 2367), Phospho-p44/42 MAPK (Erk1/2) (Thr202/Tyr204) (D13.14.4E) (cat# 4370), p44/42 MAPK (Erk1/2) (L34F12) (cat# 4696), DYKDDDDK Tag (D6W5B) Rabbit mAb (cat# 14793), Phospho-PKC substrate motif (Cat# 6967S) and β-Actin (Cat# 4970S, 3700S) were purchased from cell signaling. Anti-Ras monoclonal antibody (Cat# ab108602) was purchased from AbCam. Anti-nanoluciferase monoclonal antibody (Cat# N7000) was purchased from Promega.

1.1.3. Reagents

OptiPlate-384 (White Opaque 384-well Microplate, Cat#6007290), Corning® 96-well plates with Solid White Flat Bottom (Corning cat#3917), 96-well microplate, PS, F-Bottom (Greiner, Cat# 655900), His-Tag labeling kit RED-tris-NTA 2nd Generation (Nanotemper, Cat# MO-L018), Biocytin (Millipore, Cat# B4261), TransIT®-293 transfection reagent was purchased from Mirus (MIR 2700 and MIR 2300). cOmplete™, Mini, EDTA-free Protease Inhibitor Cocktail (Cat#04693159001) and Inhibitor, Phosphatase, PhosSTOP (Cat#4906837001) were purchased from Millipore Sigma. M-PER™ Mammalian Protein Extraction Reagent (Cat#78501), Pierce™ BCA Protein Assay Kit (Cat#23225), Pierce™ Anti-HA Magnetic Beads (88837), Pierce™ Anti-DYKDDDDK Magnetic beads (A36797), and NuPAGE 4-12% Bis-Tris Protein Gels (Cat#NP0336 or Cat#NP0335), DMEM (Gibco, Cat# 11965092), Opti-MEM I (cat# 11058-021) was purchased from Life Technologies.

**1. 2. Cloning of cTag and mgTag.**

C1 domain and mgTag gene blocks were designed and purchased from genewiz. Gibson assembly ligation was performed to generate the pET23a-His10X-MBP-C1 domain construct and pSV-mgTag-NLuc-IRES2-FLuc, pSV-mgTag-KRASG12D construct. The successful clones were confirmed by sequencing and stored for future use.

**1. 3. Site-directed mutagenesis for rationally designed cTag and mgTag variants.**

All mutations and combinations were created using the Q5 Site-Directed Mutagenesis Kit (New England Biolabs, Cat# E0554S). Primers were designed for each specific cTag and mgTag variant according to the desired mutations, using the wild-type plasmid as a template for generating the cTag and mgTag variants. In brief, after performing the site-directed mutagenesis (SDM) PCR, the amplified template plasmid was added directly to a specialized Kinase-Ligase-DpnI (KLD) enzyme mix, resulting in rapid circularization at room temperature within 5 minutes while also removing the template. This mixture was then transformed into high-efficiency NEB 5-alpha

Competent *E. coli* (New England Biolabs, Cat# C2987H). The mutations were confirmed through sequencing, and the plasmids were stored for future use.

#### 1. 4. Protein expression and purification (cTag and variants)

Plasmids containing various C1 domains fused with His-10X-MBP variants in the pET23a-His6X-MBP-C1 domain were transformed into BL21(DE3) competent cells (Thermo Fischer Scientific, Cat# EC0114) using a chemical transformation method. The primary culture was inoculated using a single colony grown overnight in 10 ml of TB media containing 50 µg/ml of ampicillin (GoldBio, Cat# A-301-5) at 37°C with shaking at 200 rpm. The secondary culture was prepared by inoculating 1% of the primary culture into 1 L of TB media with ampicillin, and the culture was allowed to grow until it reached an $OD_{600nm}$ of 0.6 then shifted to 18°C for 30 minutes and induced by adding 0.3 mM isopropyl -β-D-1-thiogalactosidease (Thermo Fischer Scientific, Cat# 15529019) and supplemented with filtered sterile 100 µM $ZnCl_2$ (Sigma, Cat# 208086), followed by further growth for an additional 16 hours at 200 rpm. Cells were harvested by centrifuging at 6000 rpm (Thermo Fischer Scientific, Sorvall LYNX 6000) for 10 minutes at 4°C. The supernatant was discarded, and the cell pellet was resuspended in lysis buffer (20 mM Tris-Cl, pH 7.4, 150 mM NaCl, 100 µM $ZnCl_2$, 1 mM TCEP, and 10% glycerol). All the subsequent steps were performed at 4°C in the cold room.

Cells lysis was initiated by adding 1 mg/ml of lysozyme along with 0.5 mM PMSF (Thermo Fischer Scientific, Cat# 36987), and 1X protease inhibitor cocktail (Pierce, Cat# A37989) (one tablet in 1 ml of lysis buffer). The cells were incubated for 1 hour with gentle rocking. 0.5% of Triton X-100 (Sigma, Cat# X100) was then added to the lysate, and the mixture was further incubated for 15 minutes. Cells were sonicated using a 250-watt ultrasonic processor (Fischer Scientific, Cat# SFX250) with a 6 mm diameter probe, applying a cycle of 8 seconds ON and 15 seconds OFF for 30 minutes. The lysate was then centrifuged at 16000 rpm for 1 hour. The cell debris was discarded, and the clarified lysate was subjected to affinity chromatography by incubating it with 1.4 ml of pre-equilibrated His-Pur Nickel nitriloacetic acid (Ni-NTA) resin (Thermo Scientific, Cat# 88221) overnight. The resin was washed thrice with 15 column volumes of washing buffer (20 mM Tris-Cl, pH 7.4, 150 mM NaCl, 100 µM $ZnCl_2$, 1 mM TCEP, 40 mM imidazole, and 10% glycerol). Histidine-tagged proteins were eluted using an elution buffer (20 mM Tris-Cl, pH 7.4, 150 mM NaCl, 100 µM $ZnCl_2$, 1 mM TCEP, 300 mM imidazole, and 10% glycerol). The collected protein was concentrated using 10 KDa cut-off Amicon spin concentrators (Millipore, Cat# UFC901008), centrifuged at 3500 rpm for 30 min, and subjected to gel filtration chromatography using an AKTA pure system (Cytiva). The proteins were injected into a HiLoad Superdex-200 column, 120 ml PG (Cytiva, Cat# 28989335) at a flow rate of 1 ml/min using running buffer (20 mM Tris-Cl, pH 7.4, 150 mM NaCl, 100 µM $ZnCl_2$,1 mM TCEP, and 10% glycerol). The protein-containing fractions were analyzed using NuPAGE Bis-Tris gels with a 4-12% gradient (Thermo Scientific, Cat# NP0321BOX) to determine the final purity. The purified protein was quantified using BCA (Pierce Cat# A55864), snap-frozen in liquid nitrogen, and stored in aliquots at -80°C until future use.

#### 1. 5. Circular Dichroism spectroscopy for C1 domain peptide.

Chemically synthesized C1 domain peptide was reconstituted in peptide buffer (25 mM Tris-Cl, pH 7.4, 100 mM NaCl, 1 mM TCEP) with a master stock of 2 mM. The peptide was dialyzed overnight in assay running buffer (2 mM Tris, 15 mM NaCl 0.1 mM TCEP). The secondary structure was examined by monitoring proteins' Far-UV circular dichroism (CD) from 198nm to 250nm on a spectropolarimeter (Jasco, J-815). The spectra were recorded with 100 µM C1 domain peptide in an assay running buffer. The wavelength scan was performed at a scan rate of 10 nm/min using a 1 mm path length cuvette (Starna, Cat# 21-G-1) at 25°C. Each spectrum was averaged over five scans. Raw CD signal ($\Phi_{obs}$) was converted to Mean Residue Ellipticity ($\Phi_{MRE}$) as follows:

$$\Phi_{MRE} = (100*\Phi_{obs})/[d*C*(n-1)]$$

Where $\Phi_{obs}$ is the observed ellipticity (in degrees), C is protein concentration (Molar), d is path length (in centimeters) and n is the total number of amino acids in the protein.

#### 1. 6. Microscale Thermophoresis for C1 domain binding experiments.

MST measurements were conducted using a Monolith NT.115 equipment (NanoTemper Technologies) at room temperature. 1 µM purified MBP fused C1 domain was labeled with NT647 (RED) fluorophore (NanoTemper Technologies, Cat# MO-L-018) and then activated by an LED filter (650 nm). Compounds were diluted from 10 µM to 0.004 µM across 12 concentrations using a two-fold dilution method in a 1X assay buffer. Each dilution was then combined with the NT647 labeled C1 domain. The mixed samples were incubated at room temperature for 10 minutes. Following incubation, the dilutions were transferred into the monolith NT automated standard capillary chips (NanoTemper Technologies, Cat# MO-AK002) via capillary action. The samples were then evaluated using Microscale Thermophoresis (MST) with the auto mode set to LED power and the medium IR laser power. The results were plotted as normalized fluorescence as a function of compound concentration using GraphPad Prism (version 10.2.3).

#### 1. 7. Biolayer-Interferometry experiments for C1 domain binding studies.

All the Binding kinetics were performed in 1X assay buffer (1X PBS pH 7.4, 0.1 mM $ZnCl_2$ 1 mM TCEP, 0.01% Tween-20) using 8-channel Octet R8 (Sartorius) at 25°C. 10 µM of biotinylated **compound 14** was immobilized onto SAX biosensors (Sartorius, Cat# 18-5117) for 300 sec up to a response of 0.8-1 nm. The reference biosensors were immobilized similarly except for dipping in 1X assay buffer instead of substrate DNA. All biosensors were blocked with 100 µM biocytin (Millipore, Cat# B4261) supplemented in 1X assay buffer for 300 seconds and a stable baseline was achieved for another 120 seconds. Sensors were dipped in the C1 domain varying from 4-0.25 µM and preincubated with either DMSO or indicated compounds at 10 µM maintaining the DMSO at 2%. The kinetics were recorded with an association of 300 seconds followed by dissociation for 500 seconds in 1X assay buffer.

The raw data was processed by using inbuilt ‘Octet BLI analysis 12.2 software from Sartorius. The reference biosensors were subtracted using double referencing and subtracted data were aligned with baseline for each kinetics. The aligned were further aligned by using inter-step correction and processed for Savitzky-Golay filtering. The processed data is potted using GraphPad Prism (version 10.2.3) as final binding sensograms. All binding curves for each kinetic experiment were normalized with DMSO and plotted as a relative percent binding to calculate percentage binding.

#### 1. 8. Labeling experiments for cTag using Intact mass spectroscopy

For labeling experiments, 2 µM of purified C1 domain variant L254C is incubated with 20 µM **compound 15**, in a 20 µl reaction containing 1X PBS supplemented with 1mM TCEP and 100 µM $ZnCl_2$ for 30 minutes at room temperature. The reaction was diluted 3X with 0.1% formic acid and filtered to clear any precipitation. 20 µl of filtered reaction was loaded in vials for Intact mass analysis. Intact mass analysis was performed on Waters BioAccord LC-MS System (Waters, Model #) using Acquity UPLC protein BEH C4 column, 2.1mm X 50 mm, 300 Å (Waters, Cat# 186004495) with a resolution capacity of 10 KDa-150 KDa. The column temperature was set at 80 °C, and the autosampler at 10 °C. Mobile phase A was 0.1% formic acid in the water, and mobile phase B was 0.1% formic acid in acetonitrile. 5 µl of samples were injected into the column with a flow rate of 0.4 ml/min. The separation was performed using a gradient of 100% A for 2 min, increasing linearly to 90% B for 5 min. For data analysis, the Inbuilt software UNIFI (version 2.1.2.14) was used. The raw spectra were extracted and deconvoluted using a 10,000:100,000 Da window with 1 Da resolution. The data were processed and represented as intensity counts vs mass.

#### 1. 9. Biochemical labeling of cTag using In-gel fluorescence.

To demonstrate the labeling of cTag biochemically, 2 µM of purified MBP-C1 domain treated with 2 µM of **compound 16** and DMSO in 25 µl reaction for 30 min incubation at room temperature. 10 µl of 4x Laemmli SDS-sample reducing buffer (Bio-RAD, cat# 1610747) was added. Samples were heated at 99ºC for 10 min. After cooling to room temperature, 15 µl of the reaction mixtures were resolved on NuPAGE 4-12% Bis-Tris gels (ThermoFisher, cat # NP0321BOX) with MOPS SDS running buffer (ThermoFisher, cat # NP0001) at 150V over 1.5 hours. Western blotting was performed by transferring the gel to the nitrocellulose membrane using the dry-transfer method by iBlot2 (Invitrogen, Cat# Ib21001). The membranes were blocked with 5% BSA in TBST and incubated with primary antibodies (anti-His tag, Cell Signaling, cat# 9991) for 4 hours at room temperature on a shaker. The unbound primary antibodies were washed out 3 times with TBST buffer and membranes were incubated with a secondary antibody (IRDye800CW, Donkey anti-Mouse, cat#926-32212) for 45 minutes at room temperature. Unbound secondary antibody was washed out 3 times with TBST buffer and the blot was developed on LiCor Odyssey CLx Imaging System. All primary and secondary antibodies were used at 1:1000 and 1:10,000 dilutions, respectively.

**1. 10. In cellular labeling of cTag using In-gel fluorescence in HEK293T cells.**

**Compound treatment:** HEK293T cells were seeded into a 6-well plate (Falcon, Cat# 353046) at the density of 1.2 million cells per well in 2 mL of media one day before cTag transfection (1 µg of DNA in 200 µl OPTI-MEM with 6 µl of trans-IT per well). 18 hours post-transfection, the cells were treated with **15**, at concentrations ranging from 100 nM to 12.5 nM in DMEM. After 4 hours of incubation with the compounds, the media was removed. The cells were washed with 2 mL of ice-cold PBS and lysed using 300 µl of MPER buffer per well (Thermo Scientific, Cat#78505) containing protease (Roche, Cat# 4693159001) and phosphatase inhibitor (Roche, Ca# 4906837001) cocktails (1 tablet each per 10 mL of MPER buffer). The total lysate was sonicated for 15 seconds and spun at 16,000 g for 10 minutes at 4 ºC to separate the supernatant. Total protein concentration was estimated by BCA, and concentrations were adjusted to be equal by adding lysis buffer to samples with higher concentrations.

**In-gel fluorescence:** To demonstrate the labeling of cTag with alkyne-containing **compound 15**, a copper-catalyzed click reaction with sulfoCy5.5 azide and in-gel fluorescence was performed. A click reaction was performed using 350 µM of tris(3-hydroxypropyltriazolylmethyl) amine (THPTA, Lumiprobe, cat# H4050), 100 µM solution of sulfoCy5.5 azide (BroadPharm, cat# BP-22483), 2 mM CuSO4 and 1.5 mM sodium ascorbate along with of 15ug of lysates obtained after treatment with DMSO and **15.** After 30 min incubation at room temperature, 10 µl of 4x Laemmli SDS-sample reducing buffer (Bio-RAD, cat# 1610747) was added. Samples were heated at 100 ºC for 5 min. After cooling to room temperature, 20 µl of the reaction mixtures were resolved on NuPAGE 4-12% Bis-Tris gels (ThermoFisher, cat # NP0321BOX) with MOPS SDS running buffer (ThermoFisher, cat # NP0001) at 200V over 1 hour. Western blotting was performed by transferring the gel to the nitrocellulose membrane using the dry-transfer method by iBlot2 (Invitrogen, Cat# Ib21001). The membranes were blocked with 5% BSA in TBST and incubated with primary antibody (anti-HA, Cat# 2367, Cell Signaling) overnight at 4 ˚C on a shaker in a cold room. The unbound primary antibodies were washed out 3 times with TBST buffer and membranes were further incubated with a secondary antibody (IRDye800CW, Donkey anti-Mouse, Cat# 926-32212) for 1 hour at room temperature. Unbound secondary antibody was washed out 3 times with TBST buffer and blots were developed on LiCor Odyssey CLx Imaging System. All primary and secondary antibodies were used at 1:1000 and 1:10,000 dilutions, respectively.

**1. 11. In cellular labeling of mgTag using In-gel fluorescence in HEK293T cells.**

HEK293T cells were seeded into a 6-well plate at the density of 1 million cells per well in 2 mL of media one day before transfection with nanoluciferase or KRAS fused mgTag variants (1 µg of DNA per million). Cells were treated with Compound 2 at 100 nM dose or DMSO (for screening the mgTag variants) or concentrations ranging from 0.1 nM to 250 nM. After 2 hours of incubation with the compounds, the media was removed, and the cells were washed with 2 mL of ice-cold PBS and kept at -80 ˚C. Cells were thawed on ice and lysed using 200 µL per well of MPER buffer containing protease (Roche, Cat# 4693159001) and phosphatase inhibitor (Roche, Cat# 4906837001) cocktails (1 tablet each per 10 mL of lysis buffer). The total lysate was spun at 20,000 g for 15 minutes at 4 ºC to separate the supernatant. Total protein concentration was estimated by BCA, and lysate protein concentrations were adjusted to 2 µg/µL with lysis buffer. To 20 µg of total protein, 2 µL of 6x Laemmli SDS-sample reducing buffer (ThermoFisher Scientific; J61337.AD) was added and heated at 100 ˚C for 5 min. After cooling to room temperature, samples were resolved on NuPAGE 4-12% Bis-Tris gels (ThermoFisher, cat # NP0335BOX) with MES SDS running buffer (ThermoFisher, cat # NP0002) at 120 V for 45 minutes, followed by 150 V for 35 minutes.

In-gel fluorescence was performed by washing the gels overnight with Coomassie destain solution (100 ml glacial acetic acid, 200 ml methanol, and 700 ml ddH2O) followed by imaging them on developed on LiCor Odyssey CLx Imaging System. Western blotting was performed by transferring the gel to the nitrocellulose membrane using the dry-transfer method by iBlot2 (Invitrogen, Cat# Ib21001). The membranes were blocked with 3 % BSA in TBST and incubated with 1:1000 dilution of primary antibodies (mouse anti-nanoluciferase antibody and Rabbit anti-β-Actin antibody; Rabbit anti-Ras antibody and Mouse anti-β-Actin antibody) overnight at 4 °C on a shaker. The unbound primary antibodies were washed out 3 times with TBST buffer, and membranes were further incubated with 1:10000 dilution of secondary antibodies (IRDye800CW, Donkey anti-Mouse or anti-Rabbit, Cat# 926-32212, 926-32213) for 1 hour at room temperature. Unbound secondary antibody was washed out 3 times with TBST buffer, and blots were developed on LiCor Odyssey CLx Imaging System.

**1. 12. PKC activation studies for monitoring proteome-wide phosphorylation.**

HEK293T cells were seeded into 12 well plates (Fisherscientific, Cat# 07-200-82) at a density of 0.75 million cells per well and grown overnight for adherence. Cells were incubated with compounds **15** for 4 hours and washed with cold PBS, harvested, and lysed using MPER buffer (Thermo Scientific, cat#78505) supplemented with protease (Roche, Cat# 4693159001) and phosphatase inhibitor (Roche, ca# 4906837001) cocktails. The total lysate was spun at 16 000 g for 10 min at 4 °C to separate the supernatant. The total protein concentration was estimated by BCA**.** An equal amount of total lysate (1µg/µl) was run on the automated simple western (Jess) using all required reagents from Protein Simple. The results were analyzed using the Compass SW software. Primary antibodies: anti-Phospho-PKC substrate motif, (Cell Signaling, Cat# 6967S), anti- β-Actin, Cell Signaling, Cat# 4967). All primary antibodies were used at 1:50 dilutions. Secondary antibodies: anti-rabbit-HRP (cat# 042-206, Protein Simple), used without further dilutions, and anti-mouse-NIR (cat# 043-821, Protein Simple) used at 1:20 dilution.

**1. 13. Effect of miniature tags fused with KRAS on pERK/EKR level for MAPK pathways.**

250ng of the KRAS fusion constructs were transfected into HEK293T cells transfection reagent TransIT LT1 (Mirus Bio, Cat# 2034) and incubated for 24 hours. Cells were then washed with 1xPBS and stored at -80$^0$C until proceed further. Cells were lysed using MPER buffer (cat#78505, Thermo Scientific) supplemented with protease (cat# 4693159001, Roche) cocktail. The cell lysates were spun at 16,000 g for 10 min at 4 °C to separate the supernatant. The total protein concentration was estimated by BCA assay (cat# 23225, Thermo Scientific). An equal amount of total lysate (1µg/µl) was run on the automated simple western (Jess) using all required reagents from Protein Simple. The results were analyzed using the Compass SW software. Primary antibodies: pERK (cat# 4370S, Cell Signaling Technology), total ERK (cat# 4696S, Cell Signaling Technology), and HA (cat# 2367S, Cell Signaling Technology). All primary antibodies were used at 1:50 dilutions. Secondary antibodies: anti-rabbit-HRP (cat# 042-206, Protein Simple), used without further dilutions, and anti-mouse-NIR (cat# 043-821, Protein Simple) used at 1:20 dilution.

#### 1. 14. Immunoblotting studies for GRIPs-mediated tyrosine phosphorylation of BRD4 by cTag fused with Abl in HEK293T cells.

HEK293T cells were seeded into a 6-well plate (Falcon, Cat# 353046) at the density of 1.2 million cells per well and grown 12 hr overnight for adherence. 200ng of cTag-Abl-HA and 1 µg of BDR4-Flag were co-transfected using transfection reagent TransIT LT1 (Mirus Bio, Cat# 2034). After 24 hours, cells were treated with Abl-Chimeras at 250nM concentration in DMEM for 4 hours. Cells were washed with cold PBS, harvested, and lysed using MPER buffer (Thermo Scientific, cat#78505) supplemented with protease (Roche, Cat# 4693159001,) and phosphatase inhibitor (Roche, ca# 4906837001) cocktails. The total lysate was spun at 16 000 g for 10 min at 4 °C to separate the supernatant. The total protein concentration was estimated by BCA**.**

For Immunoprecipitation, Anti-Flag-magnetic beads (Thermo Scientific, cat#A36798) 20 µl slurry per sample were taken and equilibrated twice with MPER buffer. An equal amount of the above-prepared supernatant (total protein) was added to the equilibrated beads and incubated with gentle rotation overnight at 4 ˚C in a cold room. Unbound protein was washed out using TBST buffer three times, and the bound protein was eluted with 4x SDS sample buffer (20 µl per tube) by heating at 99°C for 10 min. The eluted protein was pipetted out by magnetic separation to a fresh tube. Equal volumes of pulldown samples were resolved by 4-12% Bis-Tris NuPAGE gels (Thermo Scientific, Cat# NP0321BOX) and transferred to nitrocellulose membrane using the dry-transfer method by iBlot2 (Invitrogen, Cat# Ib21001). The membranes were blocked with 5% BSA in TBST followed by incubation with primary antibodies (anti-pTyr-pan, cat#8954, Cell Signaling; anti-HA, cat#2367, Cell Signaling) overnight at 4 ˚C on a shaker in a cold room. The unbound primary antibodies were washed out 3 times with TBST buffer and membranes were further incubated with secondary antibodies (IRDye800CW, Donkey anti-Mouse, cat#926-32212; IRDye680RD, Donkey anti-Rabbit, cat#926-68073) for 1 hour at room temperature. Unbound secondary antibodies were washed out 3 times with TBST buffer and blots were developed on LiCor Odyssey CLx Imaging System. All primary and secondary antibodies were used at 1:1000 and 1:10,000 dilutions, respectively.

#### 1. 15. Immunoblotting studies for GRIPs-mediated tyrosine phosphorylation of BRD4 by mgTag fused with Abl in HEK293T cells.

200ng of mgTag-Abl-HA and 2000 ng of 3xHA-BDR4 were co-transfected into HEK293T cells using transfection reagent TransIT LT1 (Mirus Bio, Cat# 2034). After 24 hours, cells were treated with Abl-chimeras at 1µM concentration in DMEM for 6 hours. Cells were then washed with 1xPBS and stored at -80$^0$C until proceeding further. Cells were lysed using MPER buffer (cat# 78505, Thermo Scientific) supplemented with protease (Roche, Cat# 4693159001,) and phosphatase inhibitor (Roche, ca# 4906837001) cocktails. The total lysate was spun at 16,000 g for 10 min at 4 °C to separate the supernatant. The total protein concentration was estimated by BCA (cat# 23225, Thermo Scientific). For Immunoprecipitation, Anti-HA-magnetic beads (cat# 88837, Thermo Scientific,) 10 µl slurry per sample were taken and equilibrated twice with MPER buffer. An equal amount of the above-prepared supernatant (total protein) was added to the equilibrated beads and incubated with gentle rotation overnight at 4 ˚C in a cold room. Unbound protein was washed out using 1xTBST buffer three times, and the bound protein was eluted with 1x SDS sample buffer (20 µl per tube) by heating at 95°C for 5 min. The eluted protein was pipetted out by magnetic separation to a fresh tube. Equal volumes of pulldown samples were run on the automated simple western (Jess) using all required reagents from Protein Simple. The results were analyzed using the Compass SW software. Primary antibodies: pY-1000 (cat# 8954S, Cell Signaling Technology) and HA (cat# 2367S, Cell Signaling Technology). All primary antibodies were used at 1:50 dilutions. Secondary antibodies: anti-rabbit-HRP (cat# 042-206, Protein Simple), used without further dilutions, and anti-mouse-NIR (cat# 043-821, Protein Simple) used at 1:20 dilution.

**1. 16. Kinetic experiments with miniature Tag binders.**

General procedure for the measurements of the kinetics of the reaction of alkynes with N-Ac-Cys-OMe: To a 4:1 mixture of PBS (pH=7.4) and acetonitrile (970 µl), 10 µl solution of N-(4-methoxyphenyl)acetamide (10 mM stock in DMSO), 10 µl solution of alkyne (10 mM stock in DMSO) and 10 µl solution of *N*-Ac-Cys-OMe (100 mM stock in DMSO) were added and the reaction was monitored for decay in the concentration of alkyne with respect to the internal standard at 25 °C. Assuming pseudo-first-order reaction kinetics, the natural logarithm of the alkyne concentration was plotted against different time points, and the rate constant was determined from the slope of the fitted line. Each data represents a time point of n=1 of the same reaction mixture.

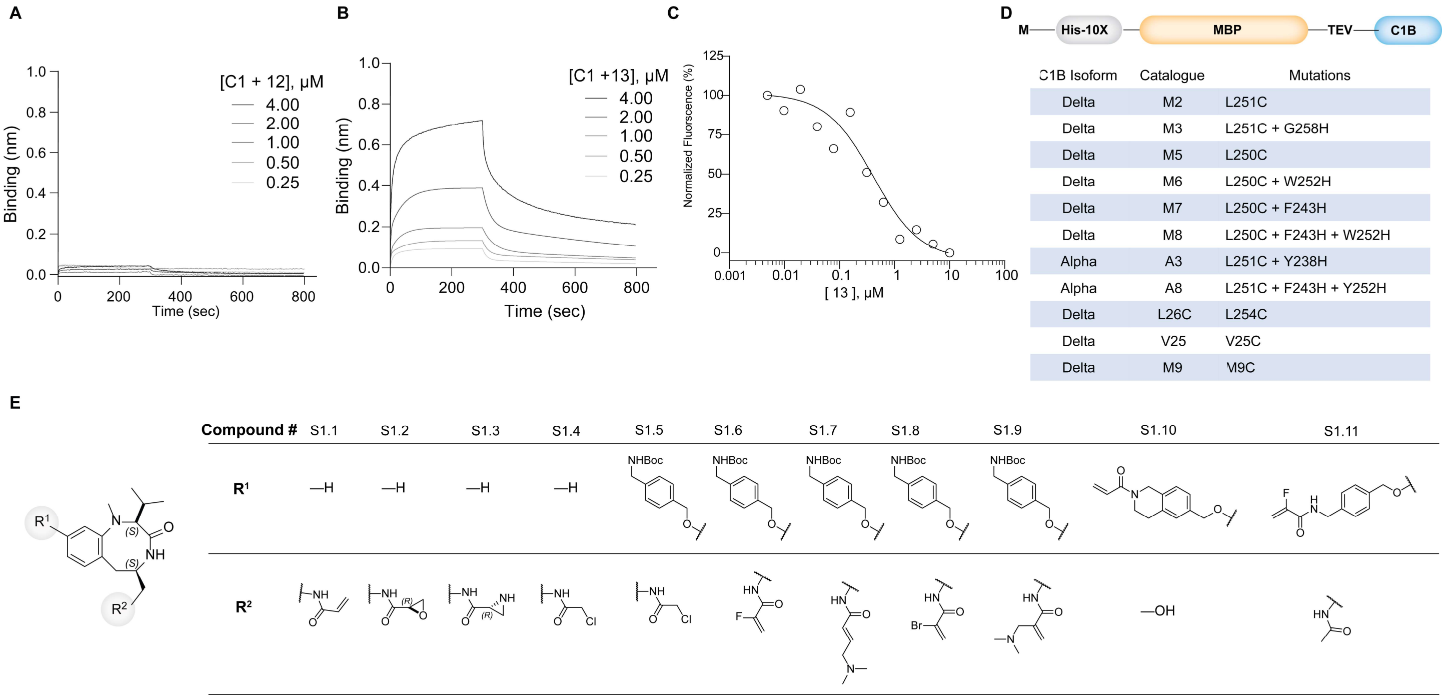

**Figure S1**. (**A,B**) BLI competition assays for compounds 12 (**A**) and 13 (**B**). (**C**) MST data for compound **13.** (**D**) Library of C1B mutants fused with MBP. All proteins were expressed in BL21(DE3) expression host and purified using Ni-NTA affinity chromatography. (**E**) Library of benzolactam analogs with binding pocket targeting electrophiles (compounds **S1.1-1.9**) or surface targeting electrophiles (**S1.10-1.11**)

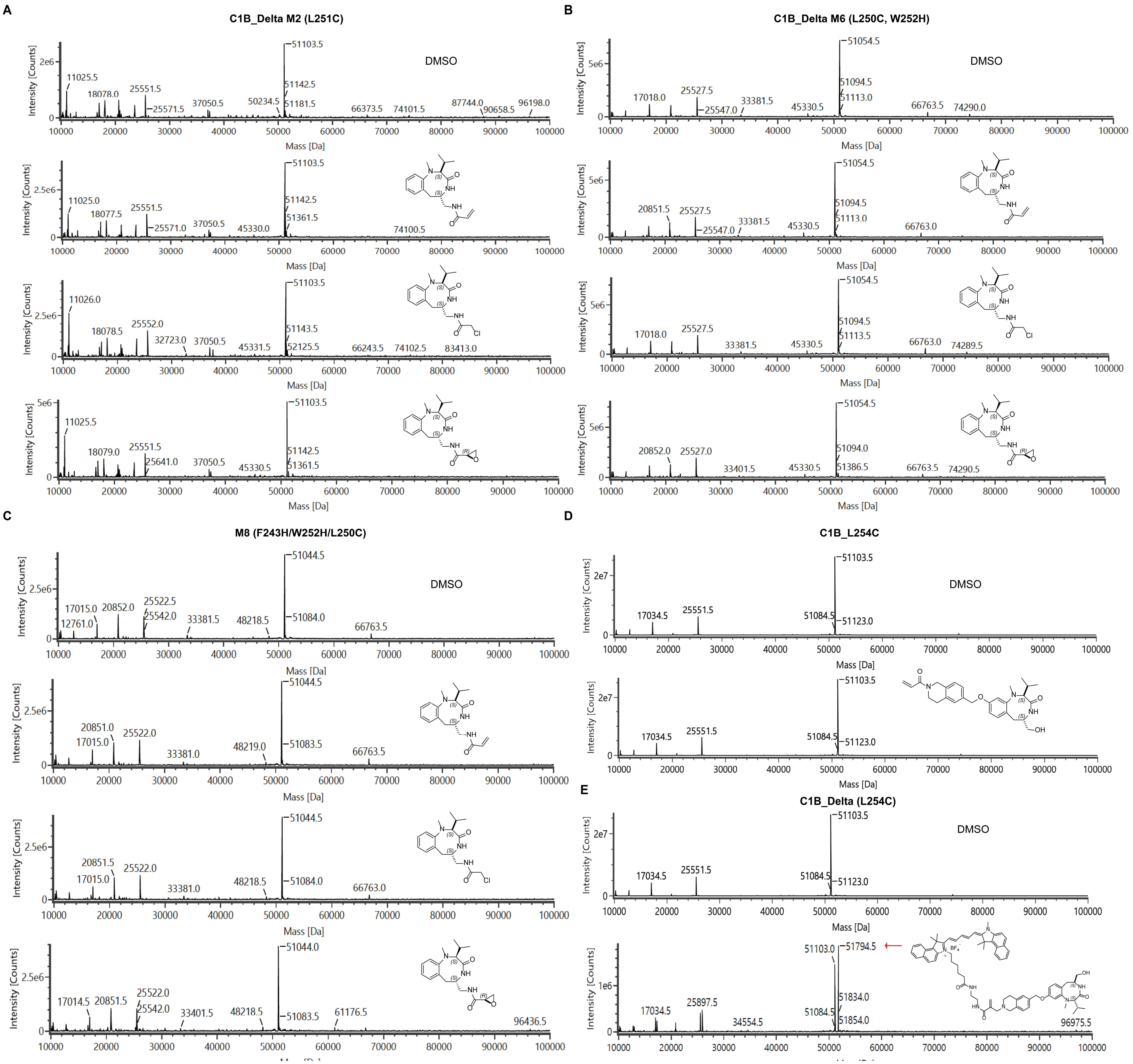

**Figure S2**. (**A**) MS spectra for labeling of C1B_delta_M2_(L251C) with probes **S1.1**, **S1.2** and **S1.4**. (**B**) MS spectra for labeling of C1B_delta_M6_(L250C and W252H) with probes **S1.1**, **S1.2** and **S1.4**. (**C**) MS spectra for labeling of C1B_delta_M8_(F243H,L250C and W252H) with probes **S1.1**, **S1.2** and **S1.4.** **(D)** MS spectra for labeling of C1B_delta_L254C (cTag) with compound **16**.

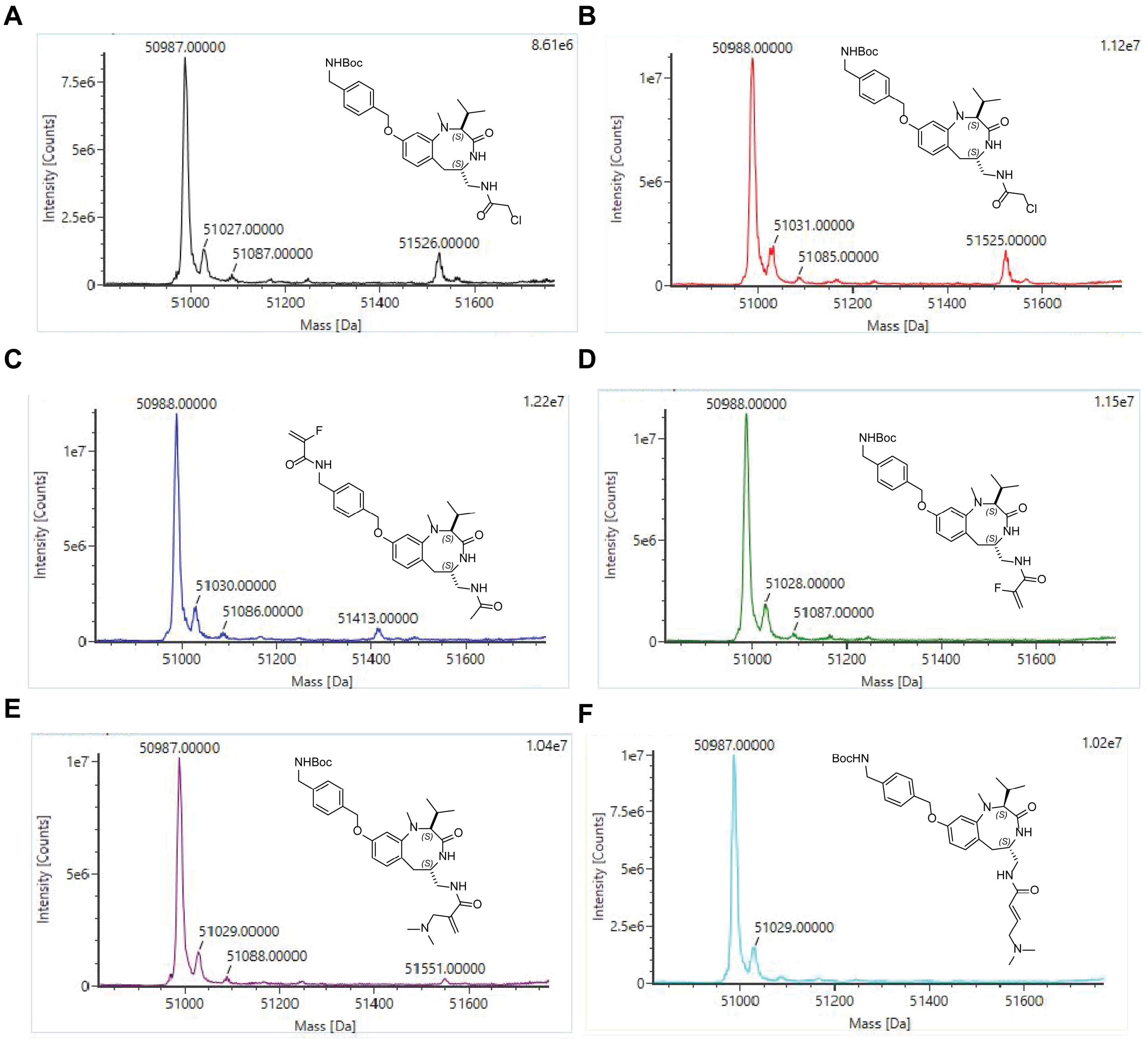

**Figure S3**. MS spectra of labeling of probes **S1.5** (**A**), **S1.5** (**B**), **S1.11** (**C**), **S1.6** (**D**), **S1.9** (**E**), and **S1.7** (**F**) with C1B L250A, W252C mutant.

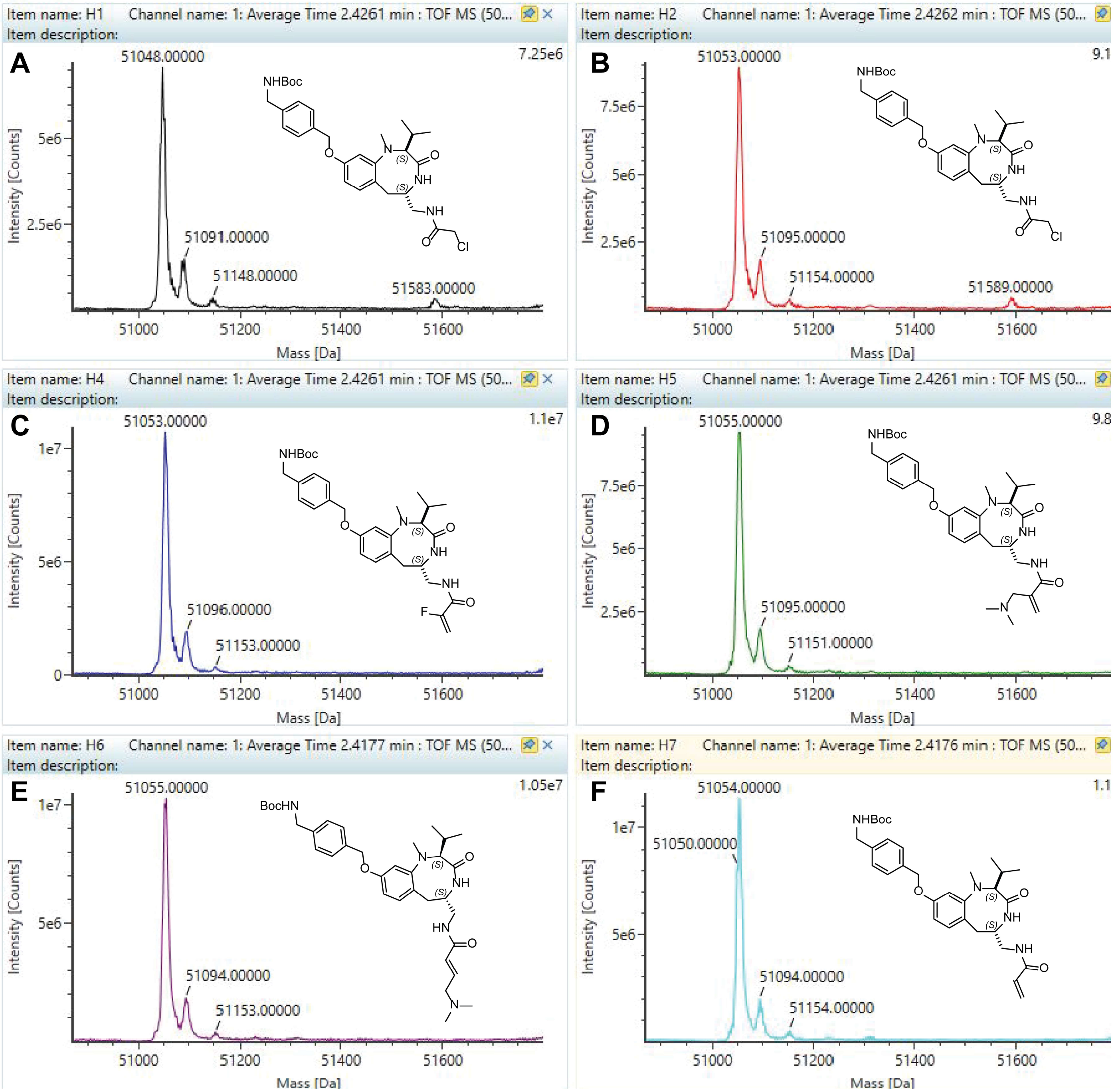

**Figure S4**. MS spectra of labeling of probes **S1.5** (**A**), **S1.5** (**B**), **S1.11** (**C**), **S1.6** (**D**), **S1.9** (**E**), and **S4.1** (**F**) with C1B L250H, W252C mutant.

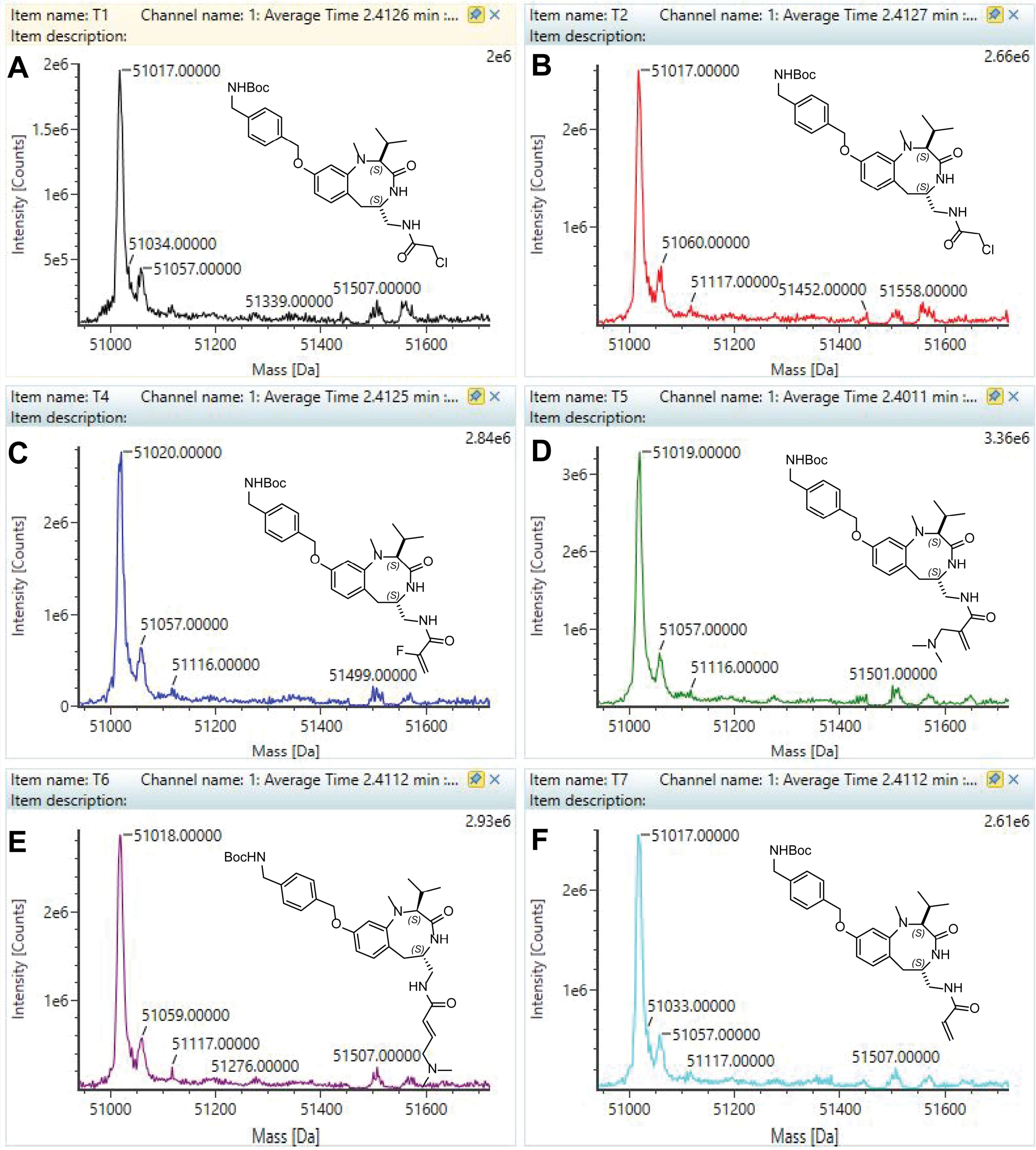

**Figure S5**. MS spectra of labeling of probes **S1.5** (**A**), **S1.5** (**B**), **S1.11** (**C**), **S1.6** (**D**), **S1.9** (**E**), and **S4.1** (**F**) with C1B L250T, W252C mutant.

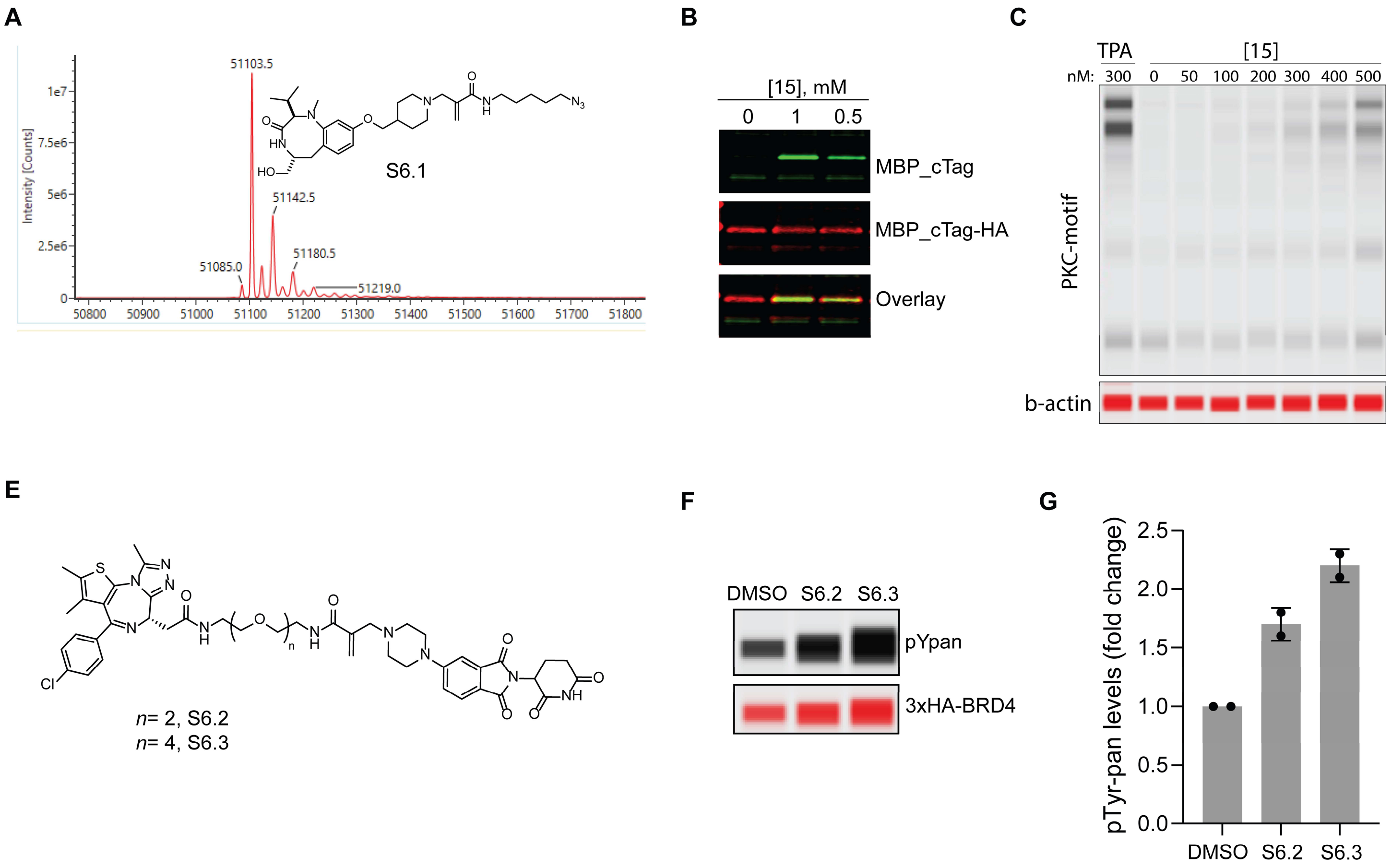

**Figure S6**. (**A**) Intact mass LCMS analysis of cTag treated with compound **S6.1**. (**B**) Pulldown of MBP-cTag using probe **15**. (**C**) Structures of probes **S6.2-3** to recruit mgTag-tagged Abl to BRD4. (**D**) Western blot analysis of proteome-wide PKC substrate phosphorylation levels using anti-Phospho-PKC substrate motif after treatment with probe **15**. (**E**) BRD4 phosphorylation by probes **S6.2-3**. (**F**) Quantitation of BRD4 phosphorylation by probes **S6.2-3** at 1 μM concentration and 6 hours tratement (n=2).

### 1. 17. Plasmid sequences

SD40 gene block:

tgccacctgacgtctaagaacgtctcaccatgtccggaCTCCTGCTCTTTTGCCCCATCTGCGGATTCACCTGTAGGCAGAA AGGGAATCTCCTTCGGCACATCAACTTGCATACAGGTGAGAAATTGTTCAAGTATCATCTGTACggcgggaga cggctcactcaaaggcggtaat

SD40 v1 gene block:

tgccacctgacgtctaagaacgtctcaccatgtccggaCTCCTGTGTTTTTGCCCCATCTGCGGATTCACCTGTAGGCAGAA AGGGAATCTCCTTCGGCACATCAACTTGCATACAGGTGAGAAATTGTTCAAGTATCATCTGTACggcgggaga cggctcactcaaaggcggtaat

SD40 v2 gene block:

tgccacctgacgtctaagaacgtctcaccatgtccggaCTCCTGCTCTGTTGCCCCATCTGCGGATTCACCTGTAGGCAGAA
AGGGAATCTCCTTCGGCACATCAACTTGCATACAGGTGAGAAATTGTTCAAGTATCATCTGTACggcgggaga
cggctcactcaaaggcggtaat

SD40 v3 gene block:

tgccacctgacgtctaagaacgtctcaccatgtccggaCTCCTGCTCTTTTGCTGTATCTGCGGATTCACCTGTAGGCAGAA
AGGGAATCTCCTTCGGCACATCAACTTGCATACAGGTGAGAAATTGTTCAAGTATCATCTGTACggcgggaga
cggctcactcaaaggcggtaat

SD40 v4 gene block:

tgccacctgacgtctaagaacgtctcaccatgtccggaCTCCTGCTCTTTTGCCCCTGTTGCGGATTCACCTGTAGGCAGAA
AGGGAATCTCCTTCGGCACATCAACTTGCATACAGGTGAGAAATTGTTCAAGTATCATCTGTACggcgggaga
cggctcactcaaaggcggtaat

SD40 v5 gene block:

tgccacctgacgtctaagaacgtctcaccatgtccggaCTCCTGCTCTTTTGCCCCATCTGCTGTTTCACCTGTAGGCAGAAA
GGGAATCTCCTTCGGCACATCAACTTGCATACAGGTGAGAAATTGTTCAAGTATCATCTGTACggcgggagac
ggctcactcaaaggcggtaat

SD40 v6 gene block:

tgccacctgacgtctaagaacgtctcaccatgtccggaCTCCTGCTCTTTTGCCCCATCTGCGGATGTACCTGTAGGCAGAA
AGGGAATCTCCTTCGGCACATCAACTTGCATACAGGTGAGAAATTGTTCAAGTATCATCTGTACggcgggaga
cggctcactcaaaggcggtaat

KRAS gene block-1: (SD40v2-HA-Linker-KRASG12D)

CTATAGGGTTAACTATTCTAGAGCTAGCgccaccATGTCCGGACTCCTGCTCTGTTGCCCCATCTGCGGATTC
ACCTGTAGGCAGAAAGGGAATCTCCTTCGGCACATCAACTTGCATACAGGTGAGAAATTGTTCAAGTATCA
TCTGTACggcggctccGGTGGTGGCGGGAGCGGAGGTGGAGGctcgAGCGGTGCGATCGCCtatccatatgatgtgc
ccgactatgcctcacttggaggaccctctggatccaccatgactgaatataaacttgtggtagttggagctgacggcgtaggcaagagtgccttgac
gatacagctaattcagaatc

KRAS gene block-2: (KRASG12D-homology arm)

Gacgatacagctaattcagaatcattttgtggacgaatatgatccaacaatagaggattcctacaggaagcaagtagtaattgatggagaaacctgtc
tcttggatattctcgacacagcaggtcaagaggagtacagtgcaatgagggaccagtacatgaggactggggagggctttctttgtgtatttgccata
aataatactaaatcatttgaagatattcaccattatagagaacaaattaaaagagttaaggactctgaagatgtacctatggtcctagtaggaaataaat
gtgatttgccttccagaacagtagacacaaaacaggctcaggacttagcaagaagttatggaattccttttattgaaacatcagcaaagacaagacag
ggtgttgatgatgccttctatacattagttcgagaaattcgaaaacataaagaaaagatgagcaaagatggtaaaaagaagaaaaagaagtcaaag
acaaagtgtgtaattatgtaatagCCTGCAGGAGAGTTTAAACCATCATCACCATCACC

**Cloning of N-SD40v2-ABL plasmid.** C-tag-ABL plasmid (sequence given below) was digested using BamHI and BsmBI enzymes and gel extracted the larger ABL fragment and ligated with the PCR amplified (using primers listed in the below table) fragments of SDv2 and ABL using Gibson ligation.

| | |
|---|---|
| SD4v2-ABL_FP | cttaagcttggtaccgagctcggatccgccaccATGTCCGGACTCCTGC |
| SDv2-ABL_RP | ccgctgaagggcttcACCACCggagccgccGTACAG |
| ABL_FP | ctccGGTGGTgaagcccttcagc |
| ABL_RP | ggtgttgaagcggctctcgga |

**C-tag-ABL plasmid sequence:**

Gaagatcctttgatcttttctacggggtctgacgctcagtggaacgaaaactcacgttaagggattttggtcatgagattatcaaaaaggatcttcacctagatc
cttttaaattaaaaatgaagttttaaatcaatctaaagtatatatgagtaaacttggtctgacagttaccaatgcttaatcagtgaggcacctatctcagcgatctgt
ctatttcgttcatccatagttgcctgactccccgtcgtgtagataactacgatacgggagggcttaccatctggccccagtgctgcaatgataccgcgagaccc
acgctcaccggctccagatttatcagcaataaaccagccagccggaagggccgagcg
cagaagtggtcctgcaactttatccgcctccatccagtctattaattgttgccgggaagctagagtaagtagttcgccagttaatagtttgcgcaacgttgttgcc
attgctacaggcatcgtggtgtcacgctcgtcgtttggtatggcttcattcagctccggttcccaacgatcaaggcgagttacatgatcccccatgttgtgcaaaa
aagcggttagctccttcggtcctccgatcgttgtcagaagtaagttggccgcagtgttatcactcatggttatggcagcactgcataattctcttactgtcatgccat
ccgtaagatgcttttctgtgactggtgagtactcaaccaagtcattctgagaatagtgtatgcggcgaccgagttgctcttgcccggcgtcaatacgggataata
ccgcgccacatagcagaactttaaaagtgctcatcattggaaaacgttcttcggggcgaaaactctcaaggatcttaccgctgttgagatccagttcgatgta
acccactcgtgcacccaactgatcttcagcatcttttactttcaccagcgtttctgggtgagcaaaaacaggaaggcaaaatgccgcaaaaaagggaataa
gggcgacacggaaatgttgaatactcatactcttcctttttcaatattattgaagcatttatcagggttattgtctcatgagcggatacatatttgaatgtatttagaa
aaataaacaaataggggttccgcgcacatttccccgaaaagtgccacctgacgtcgacggatcgggagatctcccgatcccctatggtgcactctcagtac
aatctgctctgatgccgcatagttaagccagtatctgctccctgcttgtgtgttggaggtcgctgagtagtgcgcgagcaaaatttaagctacaacaaggcaag
gcttgaccgacaattgcatgaagaatctgcttagggttaggcgttttgcgctgcttcgcgatgtacgggccagatatacgcgttgacattgattattgactagttat
taatagtaatcaattacggggtcattagttcatagcccatatatggagttccgcgttacataacttacggtaaatggcccgcctggctgaccgcccaacgaccc
ccgcccattgacgtcaataatgacgtatgttcccatagtaacgccaatagggactttccattgacgtcaatgggtggagtatttacggtaaactgcccacttgg
cagtacatcaagtgtatcatatgccaagtacgccccctattgacgtcaatgacggtaaatggcccgcctggcattatgcccagtacatgaccttatgggacttt
cctacttggcagtacatctacgtattagtcatcgctattaccatggtgatgcggttttggcagtacatcaatgggcgtggatagcggtttgactcacggggatttcc
aagtctccaccccattgacgtcaatgggagtttgttttggcaccaaaatcaacgggactttccaaaatgtcgtaacaactccgccccattgacgcaaatgggc
ggtaggcgtgtacggtgggaggtctatataagcagagctctctggctaactagagaacccactgcttactggcttatcgaaattaatacgactcactataggg
agacccaagctggctagcgtttaaacttaagcttggtaccgagctcggatccgccaccatgggagcggataagggacccgacactgatgaccccaggag
caagcacaagttcaaaatccacacttacggaagccccaccttctgcgatcactgtgggtcactgctctatggacttatccatcaagggatgaaatgtgacac
ctgcgatatgaacgttcacaagcaatgcgtcatcaatgtccccagcctctgcggaatggatcacactgagaagaagttgcttaaccaagaagcccttcagc
ggccagtagcatctgactttgagcctcagggtctgagtgaagccgctcgttggaactccaaggaaaaccttctcgctggacccagtgaaaatgaccccaac
cttttcgttgcactgtatgattttgtggccagtggagataacactctaagcataactaaaggtgaaaagctccgggtcttaggctataatcacaatggggaatgg
tgtgaagcccaaaccaaaaatggccaaggctgggtcccaagcaactacatcacgccagtcaacagtctggagaaacactcctggtaccatggcctgtg
tcccgcaatgccgctgagtatctgctgagcagcgggatcaatggcagcttcttggtgcgtgagagtgagagcagtcctggccagaggtccatctcgctgag

atacgaagggagggtgtaccattacaggatcaacactgcttctgatggcaagctctacgtctcctccgagagccgcttcaacaccctggccgagttggttcat
catcattcaacggtggccgacgggctcatcaccacgctccattatccagccccaaagcgcaacaagcccactgtctatggtgtgtcccccaactacgacaa
gtgggagatggaacgcacggacatcaccatgaagcacaagctgggcgggggccagtacggggaggtgtacgagggcgtgtggaagaaatacagcct
gacggtggccgtgaagaccttgaaggaggacaccatggaggtggaagagttcttgaaagaagctgcagtcatgaaagagatcaaacaccctaacctgg
tgcagctccttggggtctgcacccgggagcccccgttctatatcatcactgagttcatgacctacgggaacctcctggactacctgagggagtgcaaccggc
aggaggtgaacgccgtggtgctgctgtacatggccactcagatctcgtcagccatggagtacctggagaagaaaaacttcatccacagagatcttgctgcc
cgaaactgcctggtaggggagaaccacttggtgaaggtagctgattttggcctgagcaggttgatgacaggggacacctacacagcccatgctggagcca
agttccccatcaaatggactgcacccgagagcctggcctacaacaagttctccatcaagtccgacgtctgggcatttggagtattgctttgggaaattgctacc
tatggcatgtccccttacccgggaattgacctgtcccaggtgtatgagctgctagagaaggactaccgcatggagcgcccagaaggctgcccagagaagg
tctatgaactcatgcgagcatgttggcagtggaatccctctgaccggccctcctttgctgaaatccaccaagcctttgaaacaatgttccaggaatccagtatct
cagacgattacaaggatgacgacgataagtgataaacccgctgatcagcctcgactgtgccttctagttgccagccatctgttgtttgcccctcccccgtgcctt
ccttgaccctggaaggtgccactcccactgtcctttcctaataaaatgaggaaattgcatcgcattgtctgagtaggtgtcattctattctggggggtggggtggg
gcaggacagcaagggggaggattgggaagacaatagcaggcatgctggggatgcggtgggctctatggcttctgaggcggaaagaaccagctgggg
ctctagggggtatccccacgcgccctgtagcggcgcattaagcgcggcgggtgtggtggttacgcgcagcgtgaccgctacacttgccagcgccctagcg
cccgctcctttcgctttcttcccttcctttctcgccacgttcgccggctttccccgtcaagctctaaatcgggggctccctttagggttccgatttagtgctttacggca
cctcgaccccaaaaaacttgattagggtgatggttcacgtagtgggccatcgccctgatagacggtttttcgccctttgacgttggagtccacgttctttaatagt
ggactcttgttccaaactggaacaacactcaaccctatctcggtctattcttttgatttataagggattttgccgatttcggcctattggttaaaaaatgagctgattta
acaaaaatttaacgcgaattaattctgtggaatgtgtgtcagttagggtgtggaaagtccccaggctccccagcaggcagaagtatgcaaagcatgcatctc
aattagtcagcaaccaggtgtggaaagtccccaggctccccagcaggcagaagtatgcaaagcatgcatctcaattagtcagcaaccatagtcccgccc
ctaactccgcccatcccgcccctaactccgcccagttccgcccattctccgccccatggctgactaattttttttatttatgcagaggccgaggccgcctctgcct
ctgagctattccagaagtagtgaggaggcttttttggaggcctaggcttttgcaaaaagctcccgggagcttgtatatccattttcggatctgatcaagagacag
gatgaggatcgtttcgcatgattgaacaagatggattgcacgcaggttctccggccgcttgggtggagaggctattcggctatgactgggcacaacagaca
atcggctgctctgatgccgccgtgttccggctgtcagcgcaggggcgcccggttctttttgtcaagaccgacctgtccggtgccctgaatgaactgcaggacg
aggcagcgcggctatcgtggctggccacgacgggcgttccttgcgcagctgtgctcgacgttgtcactgaagcgggaagggactggctgctattgggcga
agtgccggggcaggatctcctgtcatctcaccttgctcctgccgagaaagtatccatcatggctgatgcaatgcggcggctgcatacgcttgatccggctacct
gcccattcgaccaccaagcgaaacatcgcatcgagcgagcacgtactcggatggaagccggtcttgtcgatcaggatgatctggacgaagagcatcagg
ggctcgcgccagccgaactgttcgccaggctcaaggcgcgcatgcccgacggcgaggatctcgtcgtgacccatggcgatgcctgcttgccgaatatcat
ggtggaaaatggccgcttttctggattcatcgactgtggccggctgggtgtggcggaccgctatcaggacatagcgttggctacccgtgatattgctgaagag
cttggcggcgaatgggctgaccgcttcctcgtgctttacggtatcgccgctcccgattcgcagcgcatcgccttctatcgccttcttgacgagttcttctgagcgg
gactctggggttcgaaatgaccgaccaagcgacgcccaacctgccatcacgagatttcgattccaccgccgccttctatgaaaggttgggcttcggaatcgt
tttccgggacgccggctggatgatcctccagcgcggggatctcatgctggagttcttcgcccaccccaacttgtttattgcagcttataatggttacaaataaag
caatagcatcacaaatttcacaaataaagcatttttttcactgcattctagttgtggtttgtccaaactcatcaatgtatcttatcatgtctgtataccgtcgacctcta
gctagagcttggcgtaatcatggtcatagctgtttcctgtgtgaaattgttatccgctcacaattccacacaacatacgagccggaagcataaagtgtaaagcc
tggggtgcctaatgagtgagctaactcacattaattgcgttgcgctcactgcccgctttccagtcgggaaacctgtcgtgccagctgcattaatgaatcggcca
acgcgcggggagaggcggtttgcgtattgggcgctcttccgcttcctcgctcactgactcgctgcgctcggtcgttcggctgcggcgagcggtatcagctcact
caaaggcggtaatacggttatccacagaatcaggggataacgcaggaaagaacatgtgagcaaaaggccagcaaaaggccaggaaccgtaaaaag

gccgcgttgctggcgtttttccataggctccgcccccctgacgagcatcacaaaaatcgacgctcaagtcagaggtggcgaaacccgacaggactataaa
gataccaggcgtttccccctggaagctccctcgtgcgctctcctgttccgaccctgccgcttaccggatacctgtccgcctttctcccttcgggaagcgtggcgct
ttctcatagctcacgctgtaggtatctcagttcggtgtaggtcgttcgctccaagctgggctgtgtgcacgaaccccccgttcagcccgaccgctgcgccttatc
cggtaactatcgtcttgagtccaacccggtaagacacgacttatcgccactggcagcagccactggtaacaggattagcagagcgaggtatgtaggcggt
gctacagagttcttgaagtggtggcctaactacggctacactagaagaacagtatttggtatctgcgctctgctgaagccagttaccttcggaaaaagagttg
gtagctcttgatccggcaaacaaaccaccgctggtagcggtggtttttttgtttgcaagcagcagattacgcgcagaaaaaaaggatctcaa

### 2. Chemistry

#### 1. 18. Materials

All reagents were acquired from commercial suppliers and utilized without additional purification for chemical synthesis. Reactions were conducted in round-bottom flasks, utilizing Teflon®-coated magnetic stir bars for agitation. Environmental control was maintained for moisture and air-sensitive reactions by conducting these processes under a nitrogen or argon atmosphere. Any moisture or air-sensitive liquids or solutions were transferred using syringes flushed with nitrogen. To ensure the purity of organic solvents, they were degassed by bubbling nitrogen or argon through the liquid as necessary. Flash column chromatography was executed using silica gel (60 Å mesh, 20–40 μm) on a Teledyne Isco CombiFlash Rf system for purification. Additionally, HPLC purification was carried out on a Teledyne ISCO ACCQPrep™ equipped with an XBridge BEH Prep Column (19 mm x 250 mm). UPLC-MS analysis was performed using a Waters ACQUITY UPLC I-Class 15 PLUS system and an ACQUITY SQ Detector 2 for detailed mass spectrometric evaluation. Nuclear magnetic resonance (NMR) spectroscopy was conducted at room temperature on a Bruker AVANCE III HD 400 MHz spectrometer. The spectra were collected at 400 MHz for 1H NMR and at 101 MHz for 13C NMR. The chemical shifts for 1H and 13C are reported in parts per million (ppm), referencing residual solvent signals. The NMR solvents were sourced from Cambridge Isotope Laboratories, Inc., and data were obtained in solvents such as $CDCl_3$, $CD_3OD$, and DMSO-$d_6$. In the presentation of 1H NMR data, the following format is used: chemical shift value in ppm, multiplicity (denoted as s for singlet, br s for broad singlet, d for doublet, t for triplet, dd for doublet of doublets, and m for multiplet), integration value, and the coupling constant value in hertz (Hz). High-resolution mass spectrometry (HRMS) was conducted using an Agilent 6210/6220 ESI-TOF system.

### 1. 19. Synthetic procedures and compound characterization.

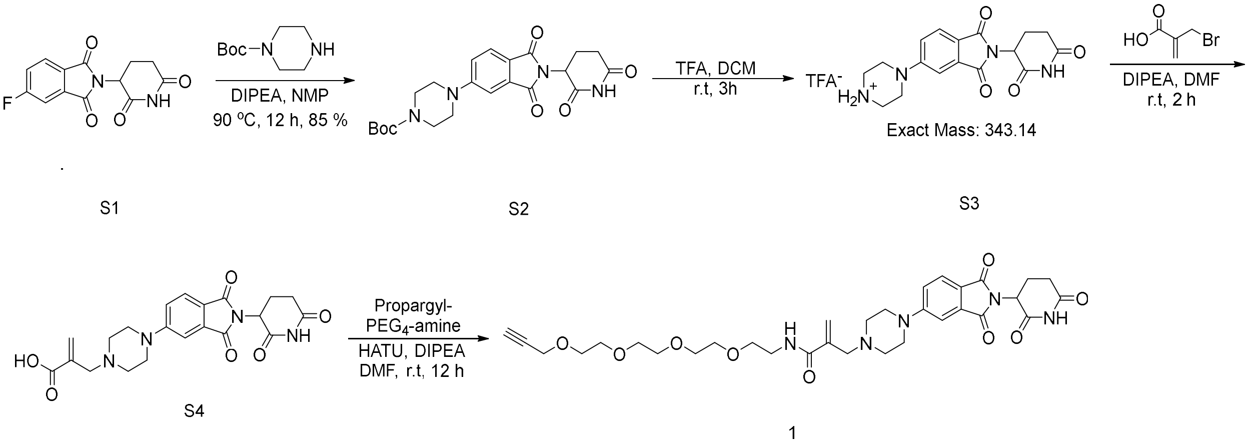

**Scheme S1**. Synthetic route to access compound **1**.

tert-butyl 4-(2-(2,6-dioxopiperidin-3-yl)-1,3-dioxoisoindolin-5-yl)piperazine-1-carboxylate (**S2**)

To a stirred solution of 5-fluoro-pomalidomide (**S1**) (69 mg, 0.25 mmol) in NMP (3 mL) were added DIPEA (2 equiv.) and N-Boc-piperazine (51 mg, 0.28 mmol, 1.1 equiv.). The resulting mixture was heated at 90 °C for 12 hr, after which it was cooled to r.t and diluted with EtOAc and Brine. The organic layer was collected, dried over $Na_2SO_4$, filtered, and concentrated under reduced pressure. The crude mixture was purified on silica gel chromatography in DCM/MeOH (0->10% MeOH) and used for the next step without further characterization.

2-((4-(2-(2,6-dioxopiperidin-3-yl)-1,3-dioxoisoindolin-5-yl)piperazin-1-yl)methyl)-N-(3,6,9,12-tetraoxapentadec-14-yn-1-yl)acrylamide (**1**).

To a stirred solution of compound (**S2**) (93 mg, 0.21 mmol) in DCM (1 mL), TFA (1 mL) was added. The resulting mixture was stirred at r.t. for 2 h, after which the volatiles were removed under reduced pressure. The crude mixture was used for the next step without any further purification. To a stirred solution of compound (**S3**) (from the previous step) in anhydrous DMF (1 mL), were added DIPEA (0.17 mL, 1 mmol, 5 equiv.) and bromomethyl-acrylic-acid (38 mg, 0.23 mmol, 1.1 equiv.). The resulting mixture was stirred at r.t for 2 h and product formation was minted by LCMS. Then HATU (117 mg, 0.31 mmol, 1.5 equiv.) and amino-PEG4-alkyne (53 mg, 0.23 mmol, 1.1 equiv.) were added and stirred at r.t for an additional 12 h. The reaction was then diluted with brine and EtOAC. The organic layer was collected, dried over $Na_2SO_4$, filtered, and concentrated under reduced pressure. The crude mixture was purified on reverse phase HPLC, providing pure compound (**1**) (28 mg, 21% yield over 3 steps). $^1$H NMR (400 MHz, MeOD) δ 7.60 (d, *J* = 8.5 Hz, 1H), 7.30 (d, *J* = 2.3 Hz, 1H), 7.18 (dd, *J* = 8.6, 2.3 Hz, 1H), 6.10 (s, 1H), 5.66 (s, 1H), 4.98 (dd, *J* = 12.5, 5.4 Hz, 1H), 4.12 – 4.02 (m, 2H), 3.51 (td, *J* = 13.8, 8.1 Hz, 18H), 3.39 (t, *J* = 5.2 Hz, 2H), 3.21 (p, *J* = 1.6 Hz, 3H), 2.98 – 2.72 (m, 5H), 2.70 – 2.54 (m, 2H), 2.07 – 1.96 (m, 1H). $^{13}$C NMR (101 MHz, MeOD) δ 173.2, 170.2, 167.8, 167.7, 167.5, 155.2, 134.2, 124.7, 118.5, 108.5, 79.3, 74.6, 70.2, 70.1, 69.9, 69.9, 69.1, 68.7, 57.6, 51.4, 49.1, 39.1, 30.8, 22.4.

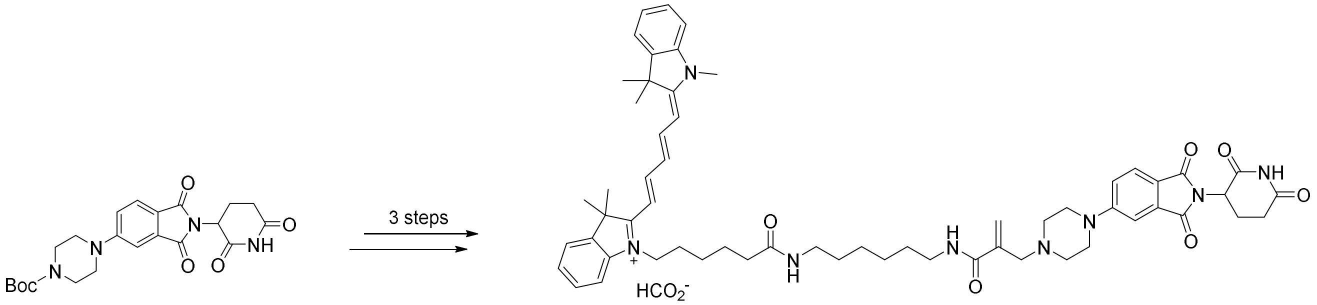

**Scheme 2**. Synthetic route to access compound **2**.

**Compound 2** was synthesized following the same synthetic route with compound **1** (scheme S1), at a 0.1 mmol scale and 17% overall yield. $^1$H NMR (400 MHz, MeOD) δ 8.47 (s, 1H), 8.25 (t, *J* = 13.1 Hz, 2H), 7.68 (dd, *J* = 8.5, 2.1 Hz, 2H), 7.50 (d, *J* = 7.4 Hz, 2H), 7.42 (tdd, *J* = 7.5, 6.1, 1.2 Hz, 2H), 7.36 (t, *J* = 2.5 Hz, 1H), 7.33 – 7.23 (m, 5H), 6.62 (t, *J* = 12.3 Hz, 1H), 6.28 (dd, *J* = 13.7, 5.2 Hz, 2H), 6.08 (t, *J* = 2.2 Hz, 1H), 5.55 (d, *J* = 1.6 Hz, 1H), 5.51 (s, 1H), 5.07 (dd, *J* = 12.5, 5.4 Hz, 1H), 4.09 (t, *J* = 7.5 Hz, 2H), 3.63 (s, 3H), 3.51 – 3.44 (m, 4H), 3.35 – 3.25 (m, 7H), 3.15 – 3.05 (m, 3H), 2.78 – 2.67 (m, 2H), 2.66 (s, 4H), 2.16 (t, *J* = 7.3 Hz, 2H), 1.89 – 1.77 (m, 1H), 1.73 (s, 8H), 1.69 – 1.63 (m, 1H), 1.59 – 1.51 (m, 2H), 1.50 – 1.25 (m, 9H). $^{13}$C NMR (101 MHz, MeOD) δ 174.2, 174.0, 173.2, 173.2, 170.2, 168.0, 167.9, 167.6, 167.5, 155.6, 154.1, 142.8, 142.2, 141.3, 141.1, 138.7, 134.2, 128.4, 128.3, 125.2,

124.9, 124.8, 124.7, 124.3, 122.0, 121.9, 119.6, 118.0, 110.7, 110.5, 108.1, 103.0, 102.9, 60.3, 53.4, 51.8, 49.1, 49.1, 49.1, 43.4, 38.8, 38.7, 35.3, 30.8, 30.1, 29.0, 28.9, 27.3, 26.8, 26.6, 26.4, 26.4, 26.2, 26.0, 25.1, 22.4.

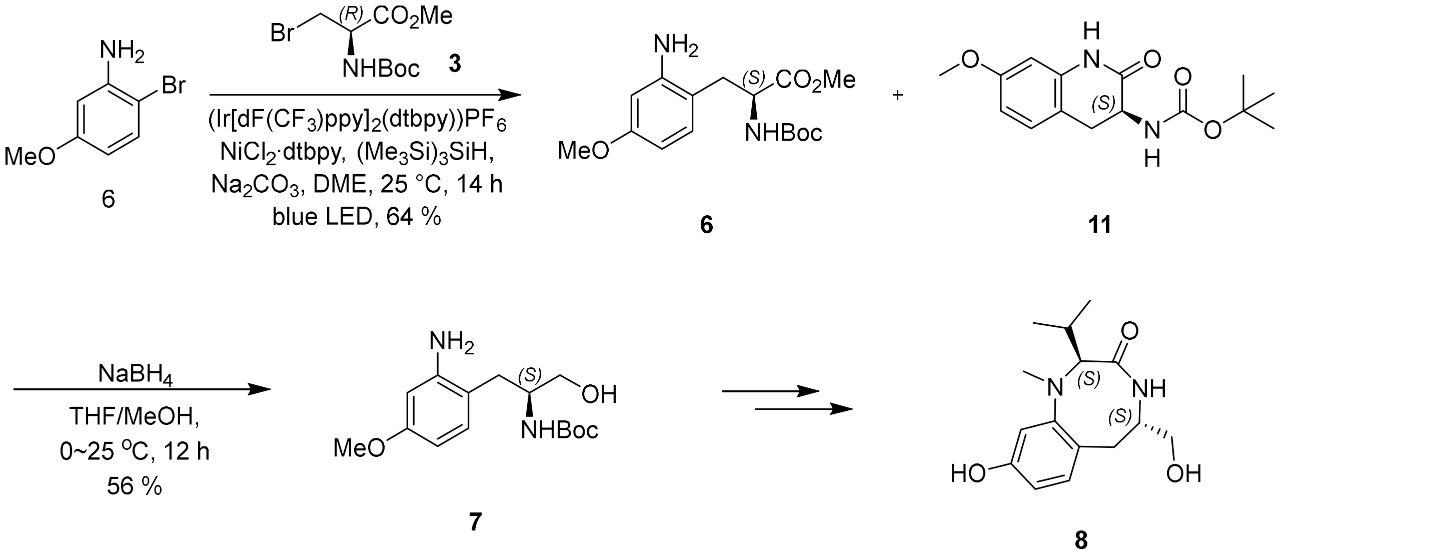

**Scheme S3**. Improved synthetic route to access benzolactam scaffold **8**. Compound 8 is synthesized from 7 following our previously reported route.[1]

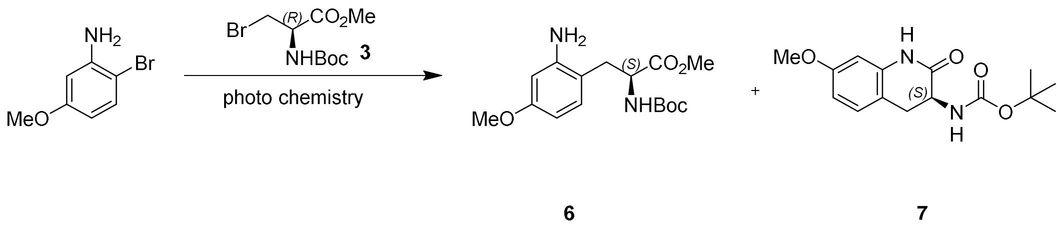

**Scheme S4**. Schematic of the photochemistry-mediated cross-coupling reaction.

**Compound 11:** To a 40 mL vial equipped with a stir bar was added 2-bromo-5-methoxy-aniline (0.5 g, 2.47 mmol, 1 eq), methyl (2R)-3-bromo-2-(tert-butoxycarbonylamino)propanoate (0.9 g, 3.22 mmol, 1.3 eq), Ir[dF($CF_3$)ppy]$_2$(dtbpy)($PF_6$) (27.8 mg, 24.8 μmol, 0.01 eq), $NiCl_2$·dtbpy (14.8 mg, 37.1 μmol, 0.015 eq), ($Me_3Si$)$_3$SiH (615 mg, 2.47 mmol, 763 μL, 1 eq) and $Na_2CO_3$ (524.58 mg, 4.95 mmol, 2 eq) in DME (25 mL). The vial was sealed and placed under nitrogen was added. The reaction was stirred and irradiated with a blue 4 × 50 W LED lamp (3 cm away), with cooling water to keep the reaction temperature at 25 °C for 14 hr. The reaction was run in 10 batches in parallel. LCMS showed methyl (S)-3-(2-amino-4-methoxyphenyl)-2-((tert-butoxycarbonyl)amino)propanoate (**6**) was the main product. On completion, each reaction mixture was quenched by $H_2O$ (30 mL), and the combined mixture was extracted with ethyl acetate (200 mL × 3). The combined organic phase was washed with saturated brine (200 mL × 3) dried over anhydrous sodium sulfate, filtered and concentrated under reduced pressure to give a residue. The residue was purified by flash silica gel chromatography with eluent of 0~15% Ethyl acetate/Petroleum ether gradient @ 100 mL/min)) to give tert-butyl (S)-(7-methoxy-2-oxo-1,2,3,4-tetrahydroquinolin-3-yl)carbamate (**11**) (6 g, 15.8 mmol, 63.86% yield, 77% purity) as a brown oil.

*General procedure for preparation of compound 1-4 - tert-butyl (S)-(1-(2-amino-4-*methoxyphenyl)-3-hydroxypropan-2-yl)carbamate*.*

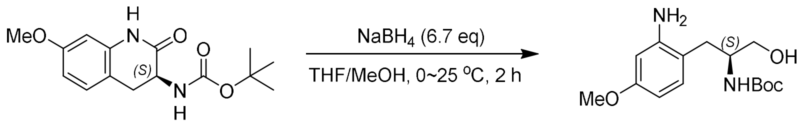

**11** **7**

**Scheme S5**. Reduction of cyclized intermediate.

**Compound 7:** To a solution of tert-butyl (S)-(7-methoxy-2-oxo-1,2,3,4-tetrahydroquinolin-3-yl)carbamate (**11**) (6 g, 15.8 mmol, 1.00 eq) in THF (25 mL) and MeOH (25 mL) was added $NaBH_4$ (4 g, 106 mmol, 6.7 eq) at 0 °C under $N_2$ atmosphere. Then, the mixture was stirred at 25 °C for 2 hr under an $N_2$ atmosphere. Upon completion, the reaction mixture was quenched by $NH_4Cl$ (100 ml) at 0 °C, and then ethyl acetate (50 mL × 3) was extracted. The combined organic phase was washed with saturated brine (30 mL × 2), dried over anhydrous sodium sulfate, filtered, and concentrated under reduced pressure to give a residue. The residue was purified by flash silica gel chromatography with eluent of 0~40% Ethyl acetate/Petroleum ether gradient to give tert-butyl (S)-(1-(2-amino-4-

methoxyphenyl)-3-hydroxypropan-2-yl)carbamate (**7**) (3 g, 8.81 mmol, 55.72% yield, 87% purity) as a brown solid. Spectroscopic data was consistent with our previous reports.[1]

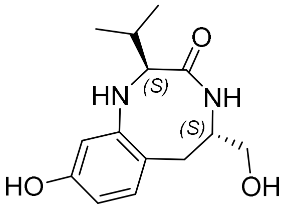

**Compound 8** was synthesized as shown in scheme **S3**, following our established protocols[1]. $^{1}$H NMR (400 MHz, MeOD) δ 6.88 (d, *J* = 8.3 Hz, 1H), 6.58 (d, *J* = 2.5 Hz, 1H), 6.40 (dd, *J* = 8.2, 2.5 Hz, 1H), 4.31 (dt, *J* = 8.7, 4.1 Hz, 1H), 3.61 (dd, *J* = 11.0, 4.7 Hz, 1H), 3.49 (dt, *J* = 7.7, 5.5 Hz, 2H), 2.97 – 2.80 (m, 2H), 2.74 (s, 3H), 2.47 – 2.34 (m, *J* = 6.8 Hz, 1H), 1.10 (d, *J* = 6.6 Hz, 3H), 0.95 (d, *J* = 6.8 Hz, 3H). $^{13}$C NMR (101 MHz, MeOD) δ 174.4, 156.7, 152.9, 132.4, 123.2, 109.7, 107.4, 73.1, 64.4, 54.1, 36.4, 35.5, 28.3, 19.4, 18.6.

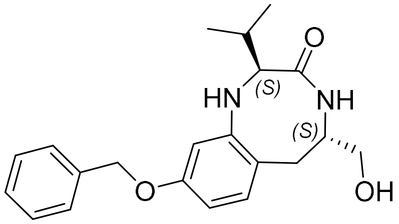

**Compound 12** was synthesized based on our previously described method[1] of phenol alkylation using benzyl bromide as the alkylating at a 10 mg scale and 98% yield. $^{1}$H NMR (400 MHz, MeOD) δ 7.46 – 7.26 (m, 5H), 6.98 (d, *J* = 8.4 Hz, 1H), 6.74 (d, *J* = 2.5 Hz, 1H), 6.59 (dt, *J* = 8.4, 2.2 Hz, 1H), 5.05 (d, *J* = 2.7 Hz, 2H), 4.26 (s, 1H), 3.61 (dd, *J* = 10.9, 4.7 Hz, 1H), 3.54 – 3.44 (m, 2H), 2.99 – 2.84 (m, 2H), 2.74 (d, *J* = 2.4 Hz, 3H), 2.47 – 2.34 (m, 1H), 1.09 (d, *J* = 6.6 Hz, 3H), 0.92 (dd, *J* = 6.8, 1.5 Hz, 3H). $^{13}$C NMR (101 MHz, MeOD) δ 174.3, 158.5, 152.8, 137.5, 132.1, 128.1, 127.4, 127.2, 124.7, 108.8, 107.4, 72.7, 69.6, 64.4, 54.0, 36.3, 35.3, 28.2, 19.3, 18.6.

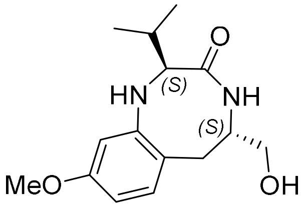

**Compound 13** was synthesized based on our previously described method[1] of phenol alkylation using MeI as the alkylating agent at a 10 mg scale and 61% yield. $^{1}H$ NMR (400 MHz, $CDCl_3$) δ 7.03 (d, *J* = 2.7 Hz, 1H), 6.96 (d, *J* = 8.3 Hz, 1H), 6.54 (d, *J* = 2.6 Hz, 1H), 6.44 (dd, *J* = 8.3, 2.6 Hz, 1H), 4.16 (s, 1H), 3.89 (d, *J* = 9.2 Hz, 1H), 3.79 (s, 3H), 3.68 (dd, *J* = 11.0, 3.8 Hz, 1H), 3.56 – 3.46 (m, 2H), 3.05 (dd, *J* = 16.8, 7.8 Hz, 1H), 2.78 (s, 3H), 2.76 – 2.70 (m, 1H), 2.43 (dp, *J* = 8.8, 6.5 Hz, 1H), 1.06 (d, *J* = 6.4 Hz, 3H), 0.88 (d, *J* = 6.7 Hz, 3H). $^{13}C$ NMR (101 MHz, $CDCl_3$) δ 174.3, 159.3, 152.6, 132.4, 123.3, 106.1, 105.8, 69.9, 65.8, 55.3, 54.6, 36.5, 35.1, 28.5, 20.5, 19.9.

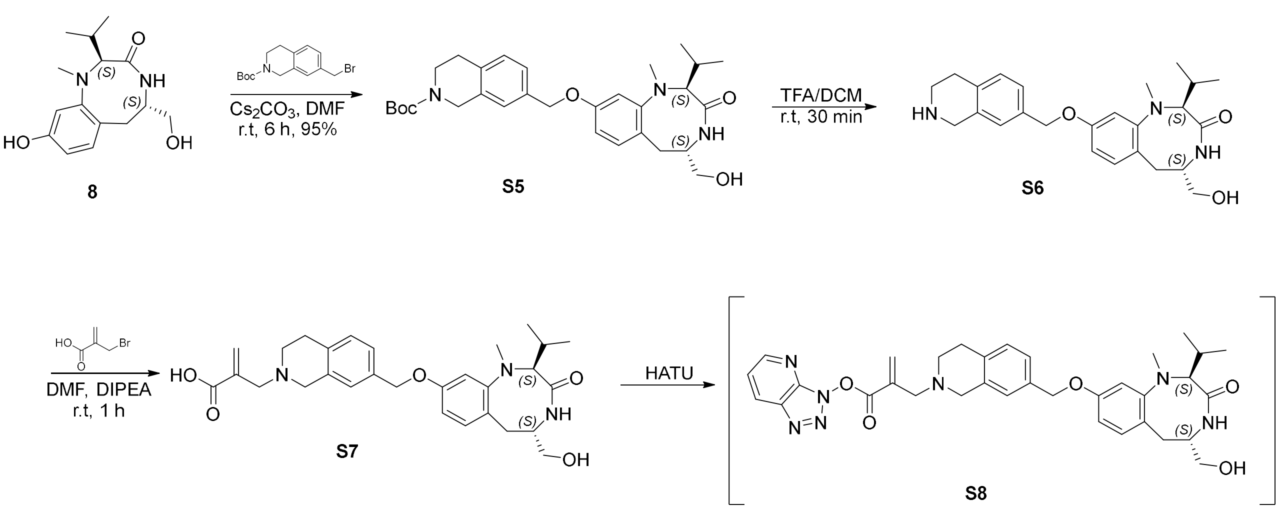

**Scheme S6**. Syntehtsis of benzolactam activated acid **S8**. This intermediate is used for the synthesis of the corresponding group transfer azido (**15**), cy5.5 (**16**), and JQ1 (**17**) probes.

**Activated benzolactam methylacrylic acid S8**: 2-((7-((((2S,5S)-5-(hydroxymethyl)-2-isopropyl-1-methyl-3-oxo-1,2,3,4,5,6-hexahydrobenzo[e][1,4]diazocin-9-yl)oxy)methyl)-3,4-dihydroisoquinolin-2(1H) yl)methyl)acrylic acid. The alkylated compound was accessed as previously described using tert-butyl 7-(bromomethyl)-3,4-dihydroisoquinoline-2(1H)-carboxylate as the alkylating agent at 111 mg scale and 95% yield. The product was monitored by LCMS and used for the next step without further characterization. To access the desired analogs a desired amount of N-Boc compound (depending on the scale) was deprotected in TFA/DCM (30 %) at r.t. After the reaction completion (30 min) the volatiles are removed under reduced pressure. The resulting compound is then dissolved in DMF and DIPEA (5 equiv.). To this solution 2-(bromomethyl)acrylic acid (1 equiv.) is added and the resulting solution is stirred at r.t for 1 h. Product formation is monitored by LCMS, and upon completion, HATU is added. The resulting activated acid is used for the next steps without further analysis.

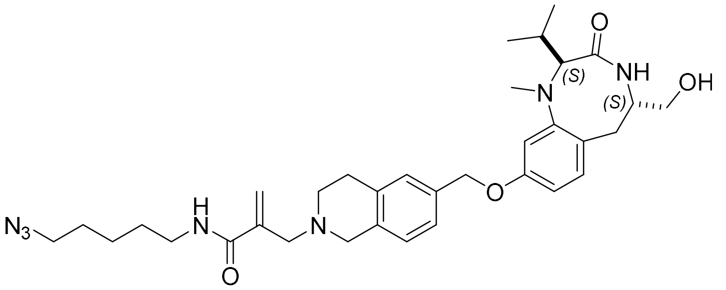

**Compound 15**: To a stirred solution of activated methacrylic acid **S8** (synthesized as shown in scheme S6 at a 0.05 mmol scale) 5-azidopentan-1-amine (2 equiv.) and DIPEA (2 equiv.) were added. The resulting mixture is stirred at r.t and product formation is monitored via LCMS. Upon reaction completion the crude mixture is purified on reverse phase HPLC using $H_2O$/MeCN plus 0.1% formic acid as mobile phase and gradient of 10-100% MeCN, providing pure azido-compound (**15**) at 63% overall yield. $^1$H NMR (400 MHz, MeOD) δ 7.26 – 7.19 (m, 2H), 7.09 (dd, *J* = 8.2, 5.1 Hz, 1H), 6.98 (dd, *J* = 8.4, 3.0 Hz, 1H), 6.74 (t, *J* = 2.9 Hz, 1H), 6.57 (ddd, *J* = 8.1, 4.9, 2.5 Hz, 1H), 6.16 (d, *J* = 1.4 Hz, 1H), 5.64 (d, *J* = 1.5 Hz, 1H), 5.00 (d, *J* = 7.4 Hz, 2H), 4.25 (d, *J* = 15.4 Hz, 1H), 3.73 (s, 2H), 3.61 (ddd, *J* = 11.0, 4.7, 2.5 Hz, 1H), 3.55 – 3.44 (m, 4H), 3.31 – 3.22 (m, 2H), 2.94 (ddq, *J* = 26.3, 11.8, 6.2 Hz, 8H), 2.75 (d, *J* = 5.1 Hz, 3H), 2.40 (dtd, *J* = 13.8, 8.3, 5.3 Hz, 1H), 1.54 – 1.34 (m, 4H), 1.30 – 1.18 (m, 2H), 1.09 (dd, *J* = 6.6, 3.2 Hz, 3H), 0.92 (dd, *J* = 6.7, 4.3 Hz, 3H). $^{13}$C NMR (101 MHz, MeOD) δ 174.2, 167.9, 158.5, 152.8, 138.1, 136.0, 133.6, 132.9, 132.1, 127.7, 126.4, 125.2, 124.7, 108.6, 107.5, 72.6, 69.4, 64.4, 60.0, 54.5, 54.0, 50.8, 49.7, 38.5, 36.3, 35.3, 28.4, 28.3, 28.2, 28.1, 23.9, 19.4, 18.7.

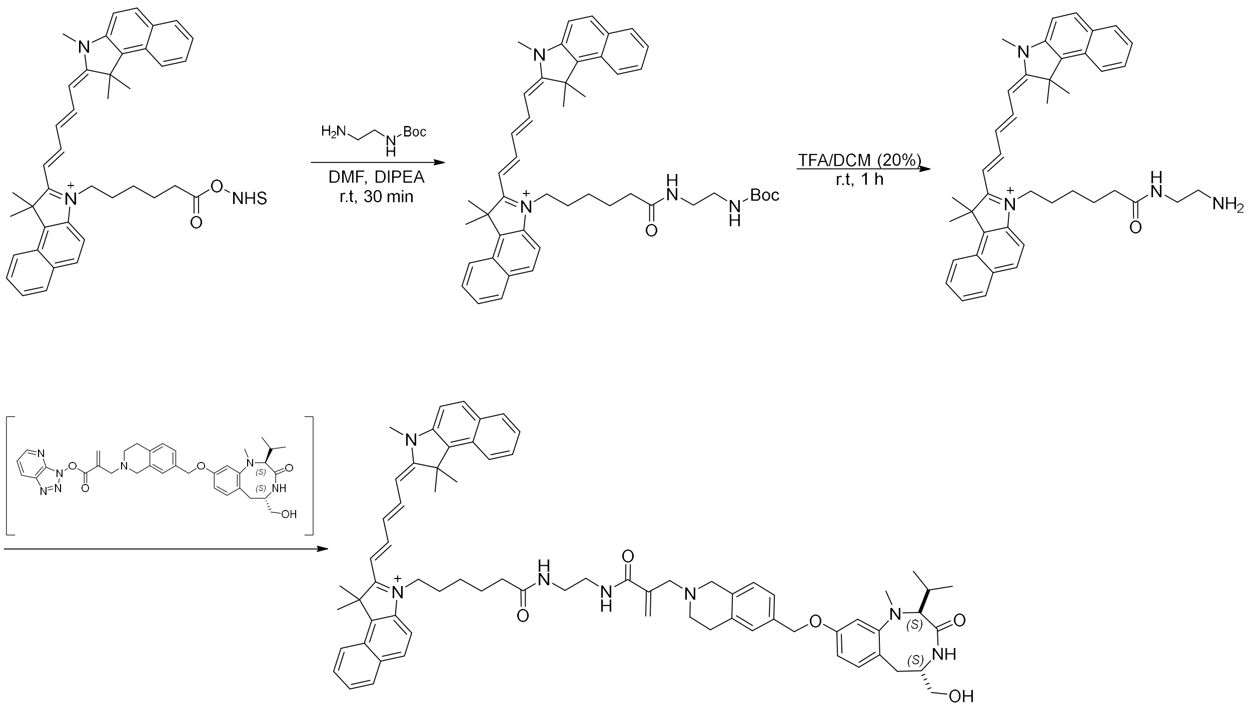

**Scheme S7**. Synthetic route for compound **16**.

**Compound 16**: To a stirred solution of Cy5.5-NHS ester (25 mg, 0.04 mmol) in anhydrous DMF, tert-butyl (2-aminoethyl)carbamate (2 equiv.) and DIPEA (2 equiv.) were added. The resulting mixture was stirred at r.t. for 30 min, and product formation was monitored via LCMS. Upon completion, the product was purified on reverse-phase HPLC and used for the next step without further characterization. The resulting Cy5.5-Nboc-Amine is dissoved in TFA/DCM (20%) and stirred at r.t for 1 h. Upon complete deprotection the volatiles are removed under reduced pressure. The Cy5.5 is then transferred into a reaction via containing 1 equiv. of benzolactam-activated acid **S8** (synthesized as shown in scheme 6.). The resulting mixture is then stirred at r.t for 1 h after which it was purified on reverse-phase HPLC using H2O/MeCN (+0.1% FA) as the mobile phase resulting 21mg of pure product (47% overall yield) . The product formation and isolation was monitored by HPLC. $^{1}$H NMR (400 MHz, MeOD) δ 8.47 (s, 1H), 8.39 – 8.27 (m, 2H), 8.23 (dd, *J* = 8.6, 3.7 Hz, 2H), 8.00 (q, *J* = 8.3 Hz, 4H), 7.69 – 7.44 (m, 5H), 7.19 – 7.09 (m, 2H), 7.03 (d, *J* = 7.8 Hz, 1H), 6.96 – 6.86 (m, 1H), 6.72 – 6.61 (m, 2H), 6.55 – 6.46 (m, 1H), 6.32 (dd, *J* = 17.2, 12.7 Hz, 2H), 6.11 (d, *J* = 1.6 Hz, 1H), 5.63 – 5.58 (m, 1H), 4.88 (s, 2H), 4.18 (q, *J* = 9.0 Hz, 2H), 3.70 (d, *J* = 17.7 Hz, 5H), 3.58 (dt, *J* = 10.9, 5.4 Hz, 1H), 3.52 – 3.44 (m, 4H), 3.33 (dq, *J* = 3.8, 1.9 Hz, 4H), 3.29 – 3.19 (m, 2H), 2.93 – 2.83 (m, 5H), 2.79 (q, *J* = 5.4 Hz, 2H), 2.67 (d, *J* = 3.8 Hz, 4H), 2.41 – 2.28 (m, 1H), 1.99 (d, *J* = 5.0 Hz, 12H), 1.88 – 1.78 (m, 2H), 1.59 (p, *J* = 7.1 Hz, 2H), 1.47 – 1.35 (m, 1H), 1.05 (dd, *J* = 10.7, 6.7 Hz, 3H), 0.88 (dd, *J* = 15.9, 7.6 Hz, 3H). $^{13}$C NMR (101 MHz, MeOD) δ 175.2, 174.7, 174.3, 174.1, 168.4, 167.4, 158.5, 152.9, 152.8, 140.2, 139.6, 138.6, 135.7, 133.8, 133.6, 133.5, 133.2, 132.1, 132.1, 132.0, 130.4, 130.3, 129.8, 128.1, 128.0, 127.6, 127.4,

127.4, 126.4, 125.2, 125.0, 124.7, 124.6, 122.0, 110.7, 110.6, 108.6, 107.4, 102.8, 102.5, 72.7, 69.4, 64.4, 59.9, 54.8, 53.9, 51.0, 49.6, 43.5, 39.0, 38.6, 38.5, 36.3, 35.4, 35.1, 30.5, 28.4, 28.2, 27.1, 26.3, 26.1, 25.9, 25.0, 19.4, 18.6.

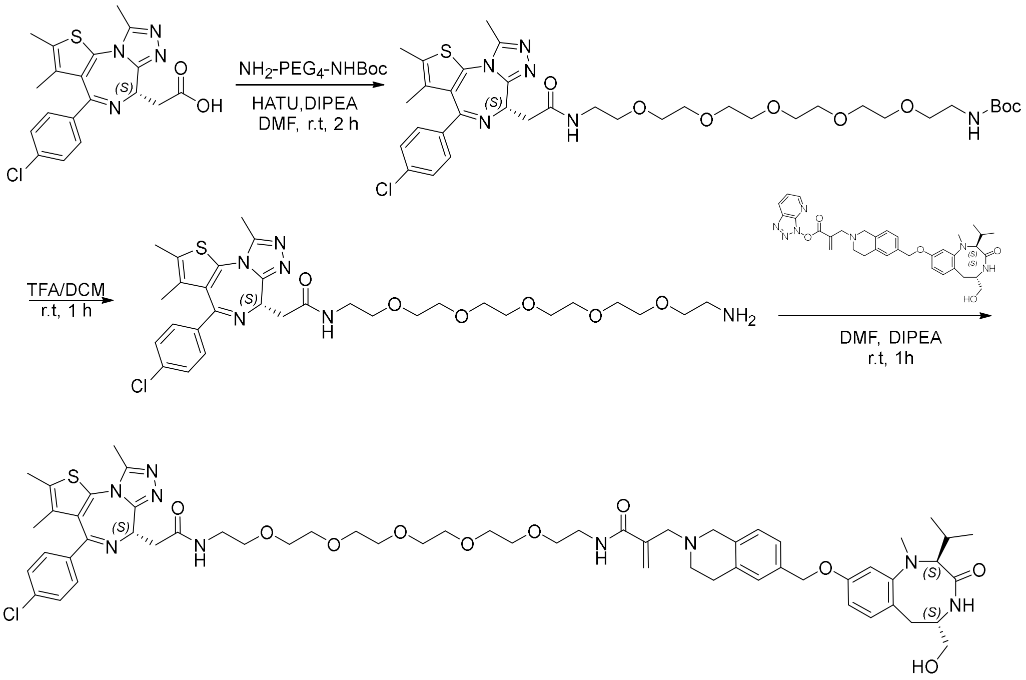

**Scheme S8**. Synthetic route for compound **17**.

**Compound 17**: To a stirred solution of JQ1-acid (40 mg, 0.1 mmol) in DMF (0.4 mL), HATU (1.5 equiv.), DIPEA (2 equiv.) and NH2-PEG4-NBoc (1.1 equiv.) were added. The resulting mixture is then stirred at r.t for 1 h . Upon completion, the reaction is then diluted with EtOAc and Brine. The organic layer is then collected, dried over anhydrous $Na_2SO_4$, filtered, and concentrated under reduced pressure. The resulting mixture is dissolved in TFA/DCM (50%), and upon completion, all volatiles are removed under reduced pressure. The resulting crude mixture is then purified on reverse-phase HPLC using $H_2O$/MeCN (+0.1% FA) as the mobile phase and used for the next step without further characterization. To access the final product, the resulting JQ1-PEG$_4$-NH$_2$ is transferred to a stirred solution of activated-benzolacatam acid **S8** (1 equiv.) and stirred at r.t for 2 h. After the completion of the reaction, the product is purified on reverse-phase HPLC using H2O/MeCN (+0.1% FA) as the mobile phase, which provides 26 mg of pure product (23% overall yield). $^1$H NMR (400 MHz, MeOD) δ 7.49 – 7.43 (m, 2H), 7.41 – 7.35 (m, 2H), 7.17 (d, *J* = 6.4 Hz, 2H), 7.03 (d, *J* = 8.2 Hz, 1H), 6.94 (d, *J* = 8.4 Hz, 1H), 6.70 (d, *J* = 2.5 Hz, 1H), 6.53 (dd, *J* = 8.4, 2.5 Hz, 1H), 6.14 (d, *J* = 1.7 Hz, 1H), 5.58 (d, *J* = 1.7 Hz, 1H), 4.96

(s, 2H), 4.62 (dt, *J* = 9.1, 4.8 Hz, 1H), 4.16 (s, 1H), 3.67 – 3.34 (m, 28H), 3.34 – 3.23 (m, 3H), 3.19 (dd, *J* = 6.1, 3.4 Hz, 2H), 2.88 (d, *J* = 5.9 Hz, 4H), 2.78 – 2.62 (m, 8H), 2.45 – 2.30 (m, 4H), 1.68 (d, *J* = 4.0 Hz, 3H), 1.05 (d, *J* = 6.5 Hz, 3H), 0.88 (d, *J* = 6.6 Hz, 3H).13C NMR (101 MHz, MeOD) δ 174.1, 171.4, 167.8, 164.6, 162.5, 158.5, 155.6, 152.8, 150.7, 138.3, 136.8, 136.5, 135.7, 134.0, 133.4, 132.2, 132.1, 131.8, 130.6, 130.6, 130.0, 128.4, 127.6, 126.4, 125.0, 124.8, 124.6, 108.5, 107.3, 72.4, 70.2, 70.2, 70.1, 70.1, 70.0, 69.9, 69.9, 69.4, 69.3, 69.1, 69.0, 64.5, 60.2, 54.8, 54.1, 53.8, 49.4, 39.2, 38.9, 37.6, 37.4, 36.4, 35.3, 28.6, 28.2, 19.4, 18.8, 13.2, 11.7, 10.3.

**1. 20. LCMS traces of key intermediates and final products.**

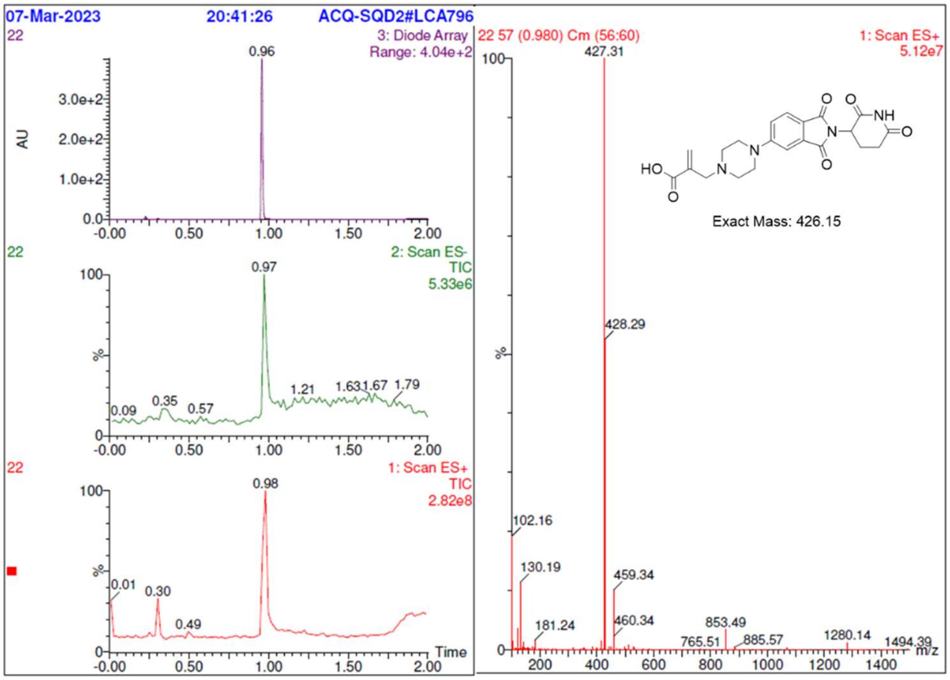

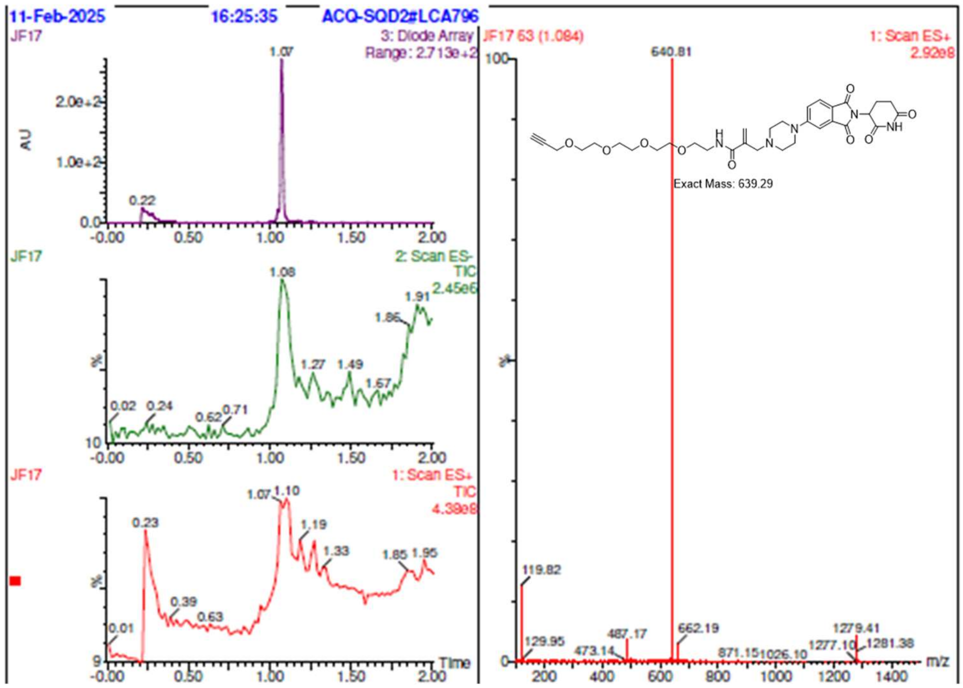
11-Feb-2025
16:25:35
ACQ-SQD2#LCA796
JF17
3: Diode Array
Range: 2.713e+2
AU
2: Scan ES-
TIC
2.45e6
1: Scan ES+
TIC
4.38e8
Time
JF17 63 (1.084)
1: Scan ES+
2.92e8
640.81
Exact Mass: 639.29
119.82
129.95
473.14
487.17
662.19
871.15
1026.10
1277.10
1279.41
1281.38
m/z

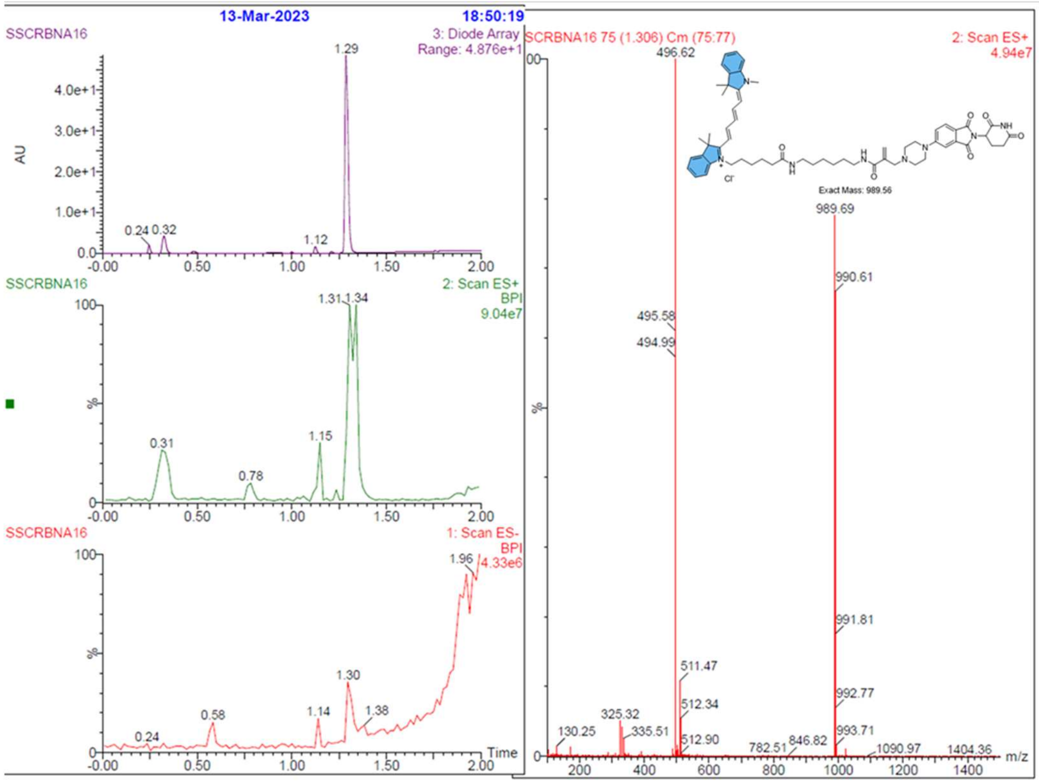
13-Mar-2023
18:50:19
SSCRBNA16
3: Diode Array
Range: 4.876e+1
AU
2: Scan ES+
BPI
9.04e7
1: Scan ES-
BPI
4.33e6
Time
SCRBNA16 75 (1.306) Cm (75:77)
2: Scan ES+
4.94e7
496.62
Exact Mass: 989.56
989.69
990.61
495.58
494.99
991.81
511.47
512.34
512.90
992.77
993.71
130.25
325.32
335.51
782.51
846.82
1090.97
1404.36
m/z

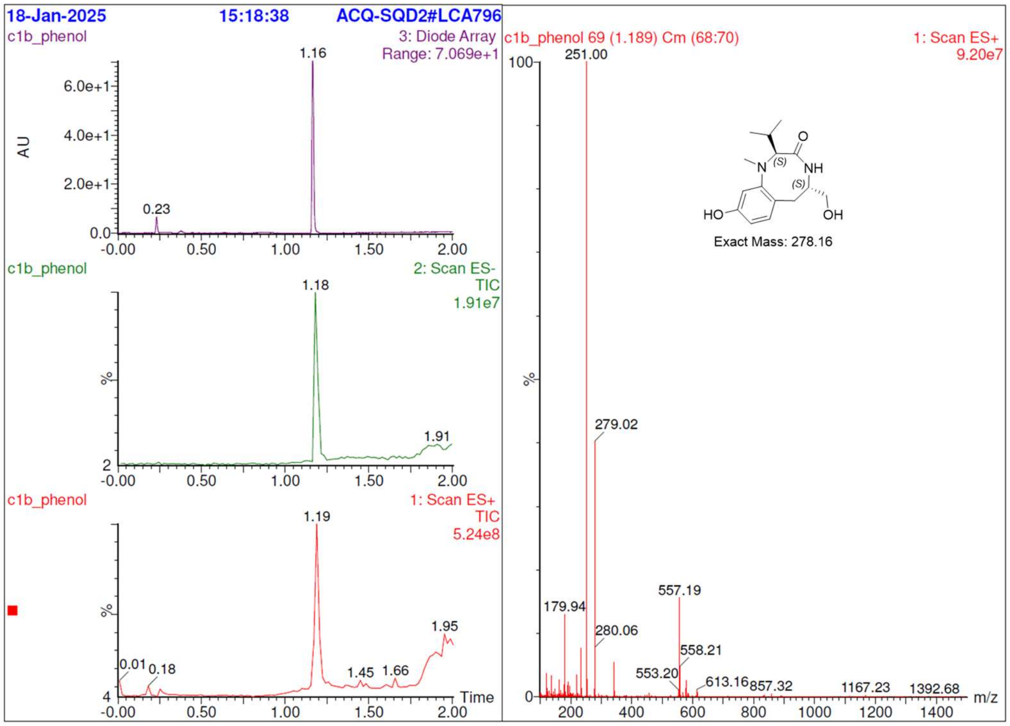
18-Jan-2025
15:18:38
ACQ-SQD2#LCA796
c1b_phenol
3: Diode Array
Range: 7.069e+1
AU
c1b_phenol
2: Scan ES-
TIC
1.91e7
c1b_phenol
1: Scan ES+
TIC
5.24e8
Time
c1b_phenol 69 (1.189) Cm (68:70)
1: Scan ES+
9.20e7
251.00
279.02
557.19
179.94
280.06
558.21
553.20
613.16
857.32
1167.23
1392.68
m/z
HO
N
(S)
O
NH
(S)
OH
Exact Mass: 278.16

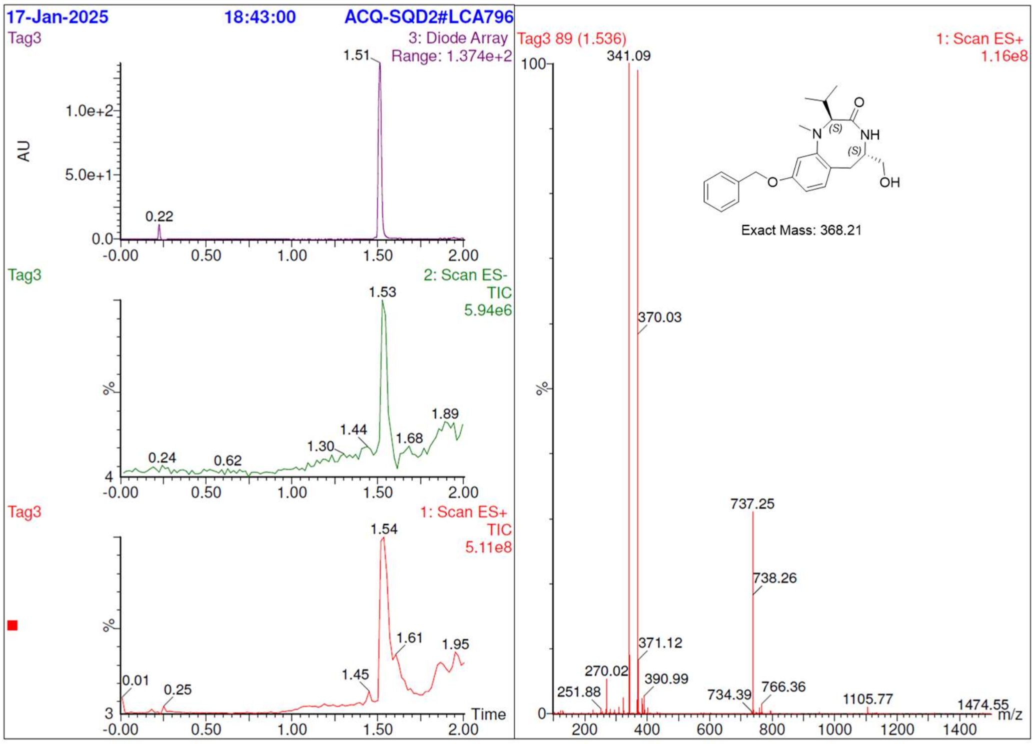
17-Jan-2025
18:43:00
ACQ-SQD2#LCA796
Tag3
3: Diode Array
Range: 1.374e+2
AU
Tag3
2: Scan ES-
TIC
5.94e6
Tag3
1: Scan ES+
TIC
5.11e8
Time
Tag3 89 (1.536)
1: Scan ES+
1.16e8
341.09
370.03
737.25
738.26
371.12
270.02
390.99
251.88
734.39
766.36
1105.77
1474.55
m/z
N
(S)
O
NH
(S)
O
OH
Exact Mass: 368.21

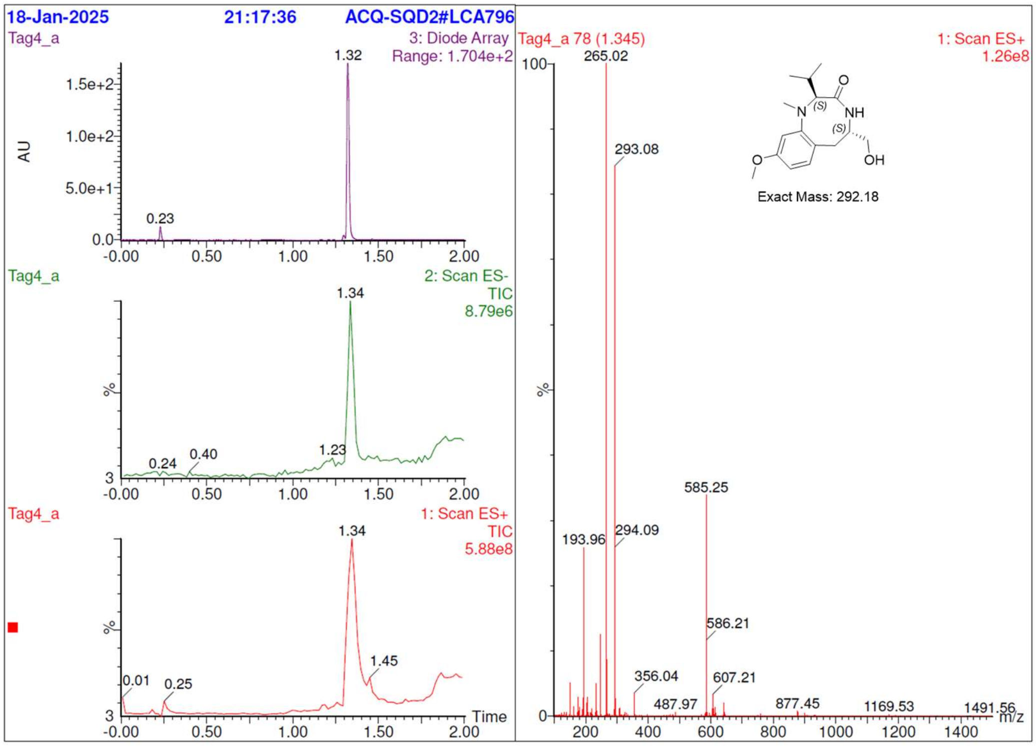
18-Jan-2025
21:17:36
ACQ-SQD2#LCA796
Tag4_a
3: Diode Array
Range: 1.704e+2
AU
1.32
0.23
Tag4_a
2: Scan ES-
TIC
8.79e6
1.34
0.24
0.40
1.23
Tag4_a
1: Scan ES+
TIC
5.88e8
1.34
1.45
0.01
0.25
Time
Tag4_a 78 (1.345)
1: Scan ES+
1.26e8
265.02
293.08
585.25
193.96
294.09
586.21
356.04
607.21
487.97
877.45
1169.53
1491.56
m/z
Exact Mass: 292.18

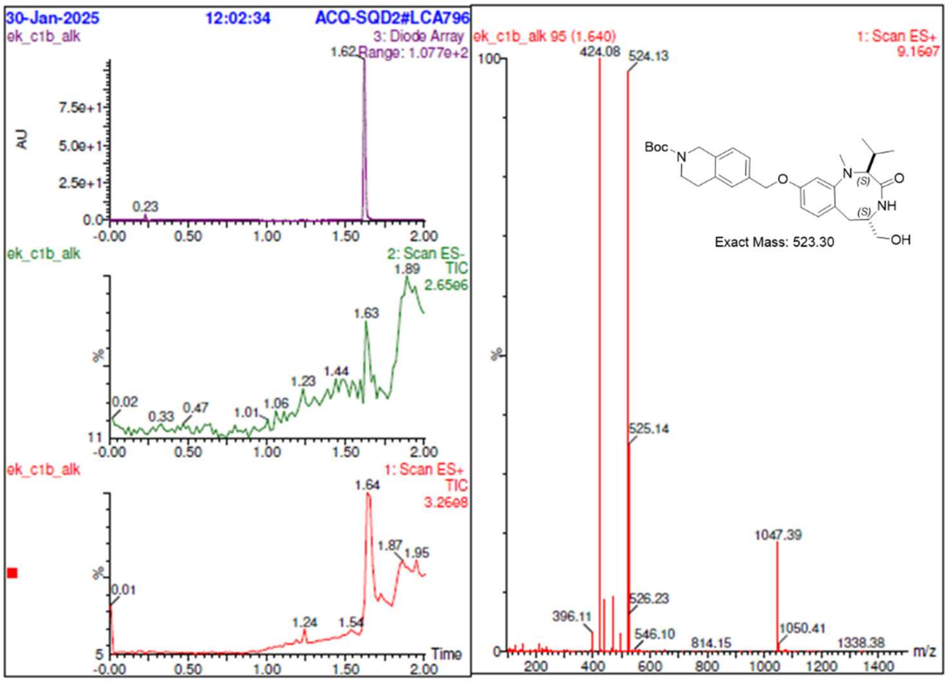
30-Jan-2025
12:02:34
ACQ-SQD2#LCA796
ek_c1b_alk
3: Diode Array
Range: 1.077e+2
AU
1.62
0.23
ek_c1b_alk
2: Scan ES-
TIC
2.65e6
1.89
1.63
1.44
1.23
0.02
0.33
0.47
1.01
1.06
ek_c1b_alk
1: Scan ES+
TIC
3.26e8
1.64
1.87
1.95
0.01
1.24
1.54
Time
ek_c1b_alk 95 (1.640)
1: Scan ES+
9.16e7
424.08
524.13
525.14
1047.39
396.11
526.23
546.10
814.15
1050.41
1338.38
m/z
Boc
Exact Mass: 523.30

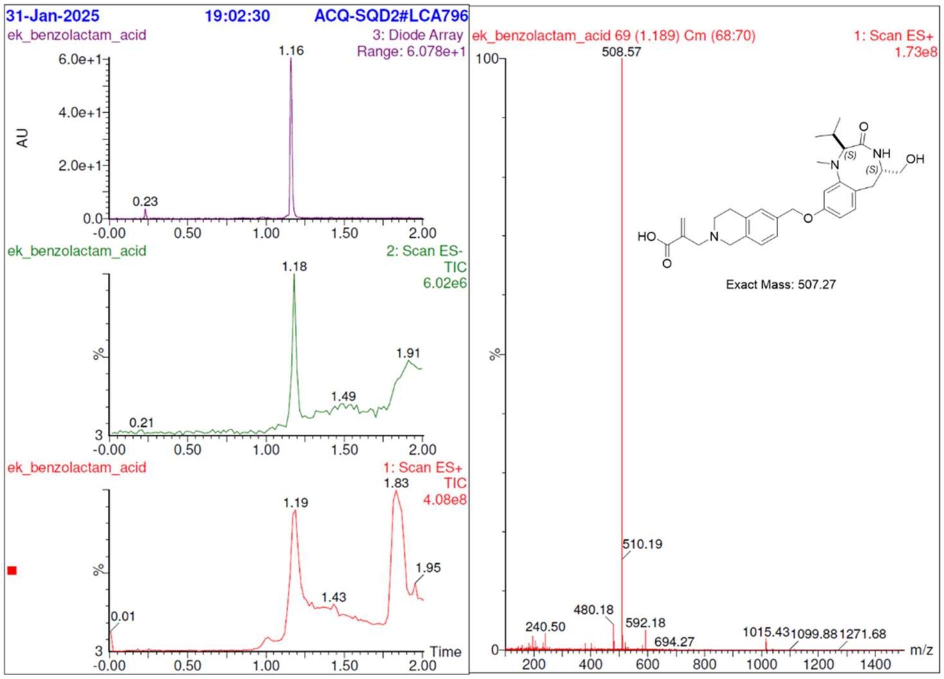
31-Jan-2025
19:02:30
ACQ-SQD2#LCA796
ek_benzolactam_acid
3: Diode Array
Range: 6.078e+1
2: Scan ES-
TIC
6.02e6
1: Scan ES+
TIC
4.08e8
ek_benzolactam_acid 69 (1.189) Cm (68:70)
1: Scan ES+
1.73e8
508.57
510.19
480.18
240.50
592.18
694.27
1015.43
1099.88
1271.68
Exact Mass: 507.27
AU
%
Time
m/z

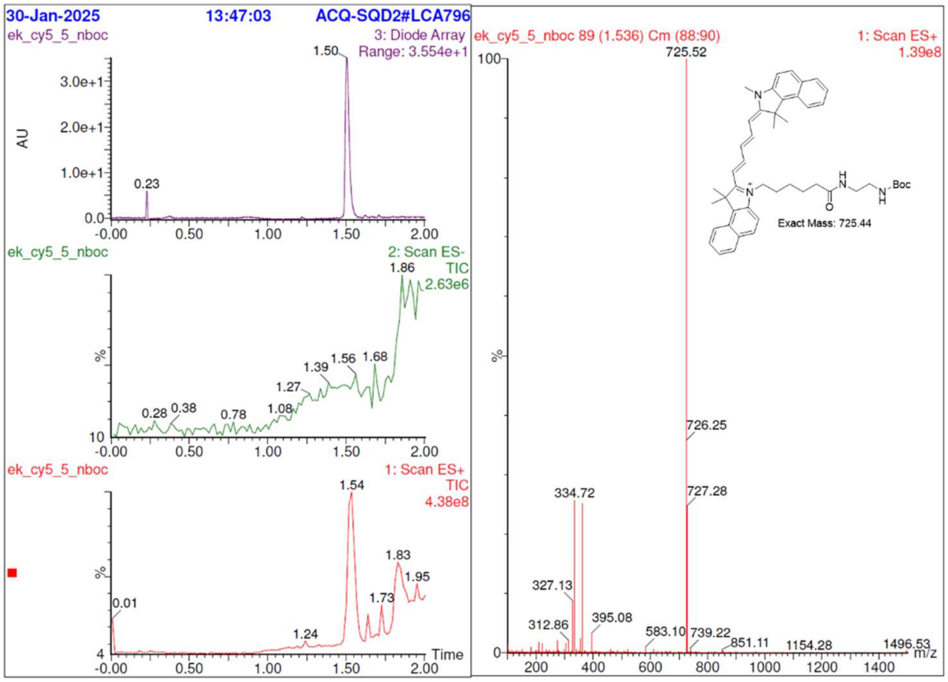
30-Jan-2025
13:47:03
ACQ-SQD2#LCA796
ek_cy5_5_nboc
3: Diode Array
Range: 3.554e+1
2: Scan ES-
TIC
2.63e6
1: Scan ES+
TIC
4.38e8
ek_cy5_5_nboc 89 (1.536) Cm (88:90)
1: Scan ES+
1.39e8
725.52
726.25
727.28
334.72
327.13
312.86
395.08
583.10
739.22
851.11
1154.28
1496.53
Exact Mass: 725.44
AU
%
Time
m/z

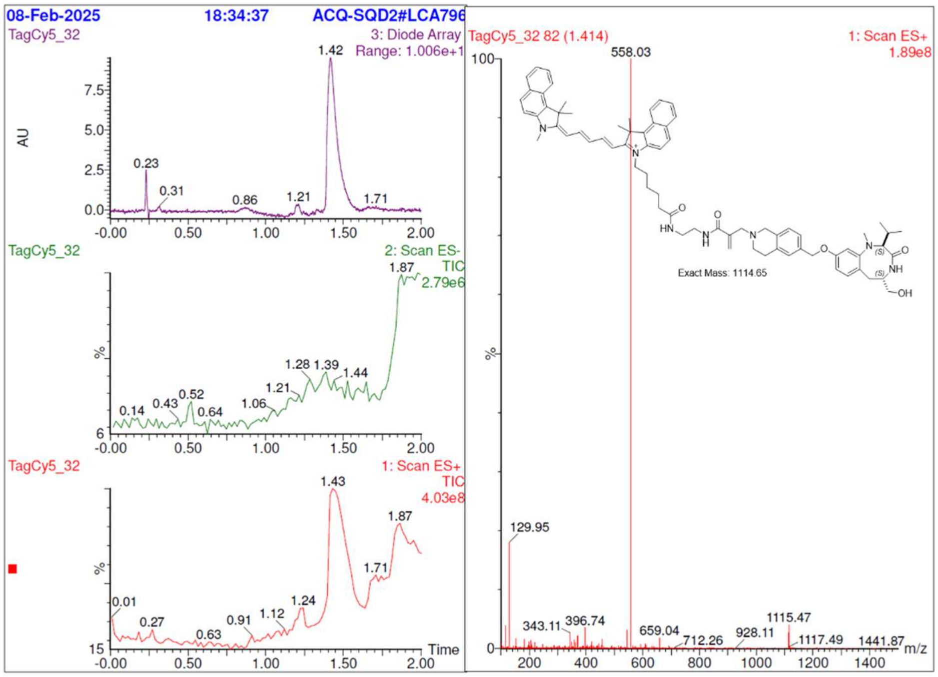

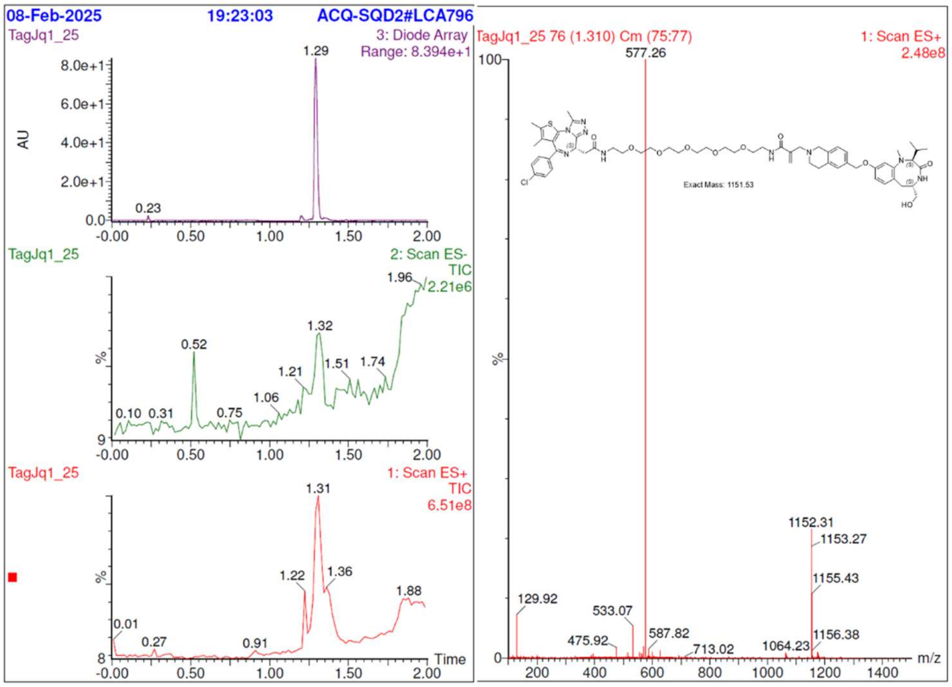

### 1. 21. HRMS spectra of final products.

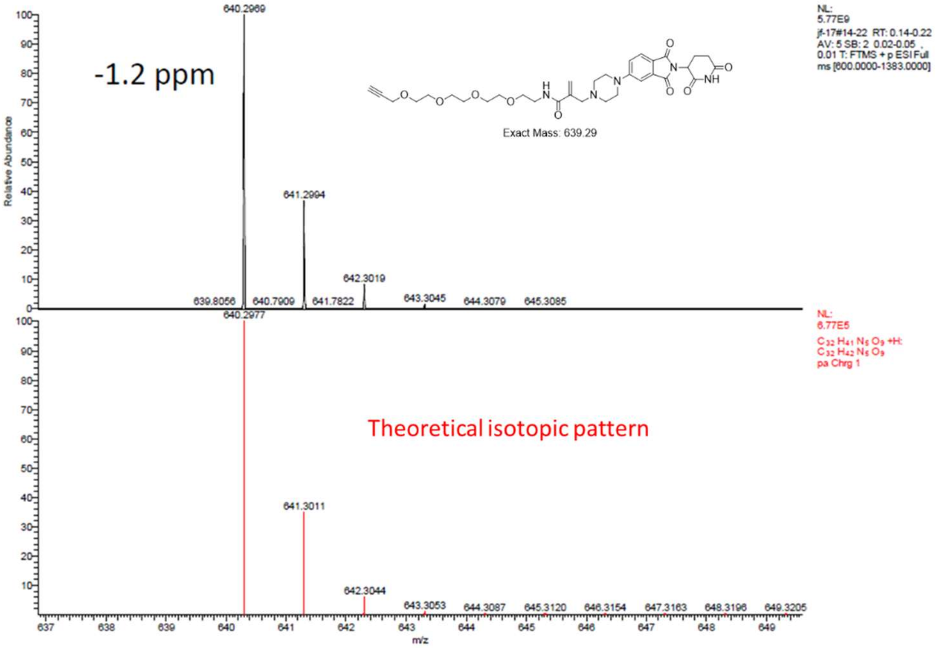
-1.2 ppm
640.2969
641.2994
642.3019
639.8056
640.7909
641.7822
643.3045
644.3079
645.3085
Relative Abundance
Exact Mass: 639.29
NL:
5.77E9
jf-17#14-22 RT: 0.14-0.22
AV: 5 SB: 2 0.02-0.05 ,
0.01 T: FTMS + p ESI Full
ms [600.0000-1383.0000]
NL:
6.77E5
pa Chrg 1
Theoretical isotopic pattern
640.2977
641.3011
642.3044
643.3053
644.3087
645.3120
646.3154
647.3163
648.3196
649.3205
m/z

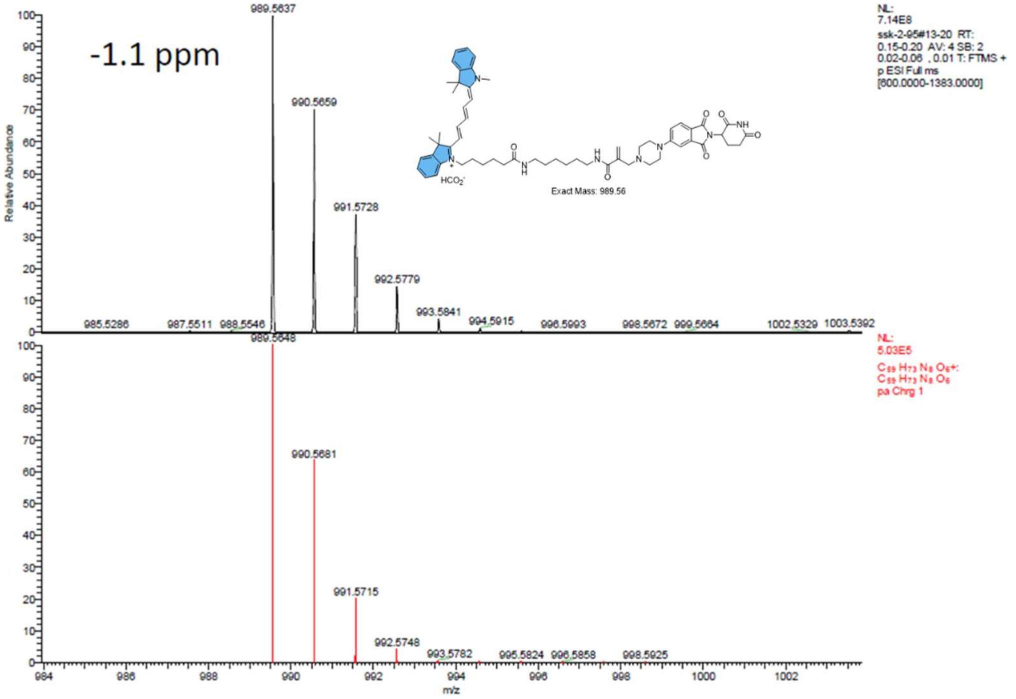
-1.1 ppm
989.5637
990.5659
991.5728
992.5779
993.5841
994.5915
996.5993
998.5672
999.5664
1002.5329
1003.5392
985.5286
987.5511
988.5546
Relative Abundance
Exact Mass: 989.56
NL:
7.14E8
ssk-2-95#13-20 RT:
0.15-0.20 AV: 4 SB: 2
0.02-0.06 , 0.01 T: FTMS +
p ESI Full ms
[600.0000-1383.0000]
NL:
5.03E5
pa Chrg 1
989.5648
990.5681
991.5715
992.5748
993.5782
995.5824
996.5858
998.5925
m/z

### 1. 22. NMR

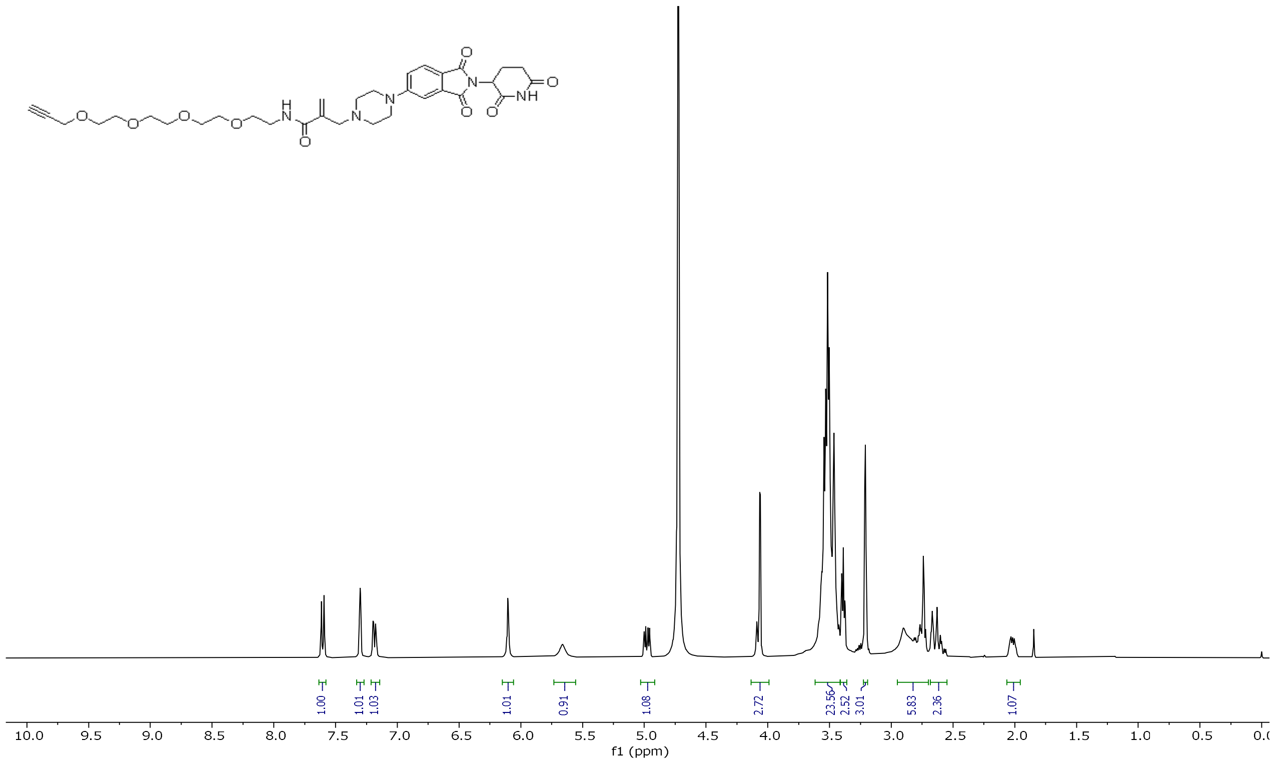
O
N
NH
O
O
N
N
H
N
O
O
O
O
O
1.00
1.01
1.03
1.01
0.91
1.08
2.72
23.56
2.52
3.01
5.83
2.36
1.07
10.0 9.5 9.0 8.5 8.0 7.5 7.0 6.5 6.0 5.5 5.0 4.5 4.0 3.5 3.0 2.5 2.0 1.5 1.0 0.5 0.0
f1 (ppm)

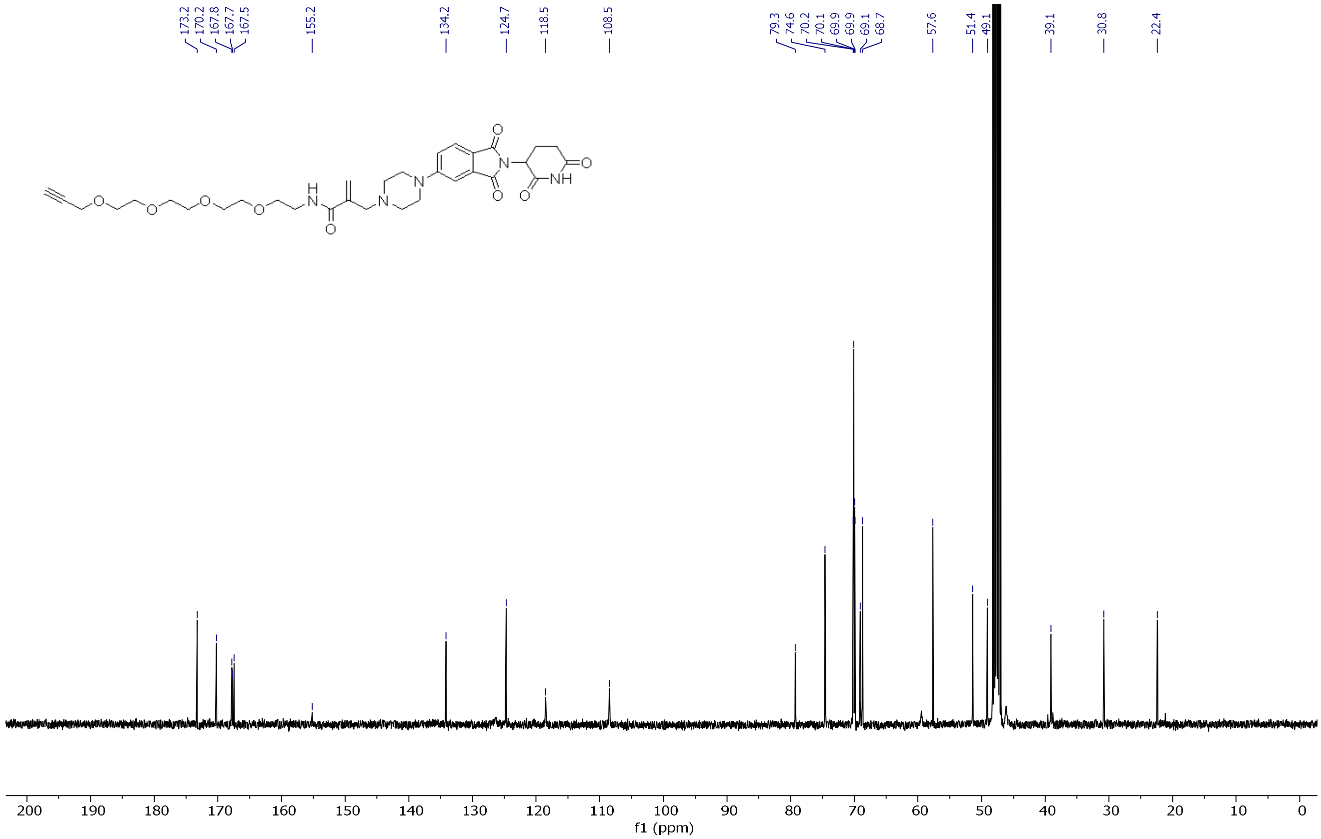
173.2
170.2
167.8
167.7
167.5
155.2
134.2
124.7
118.5
108.5
79.3
74.6
70.2
70.1
69.9
69.9
69.1
68.7
57.6
51.4
49.1
39.1
30.8
22.4
O
N
NH
O
O
N
N
H
N
O
O
O
O
O
200 190 180 170 160 150 140 130 120 110 100 90 80 70 60 50 40 30 20 10 0
f1 (ppm)

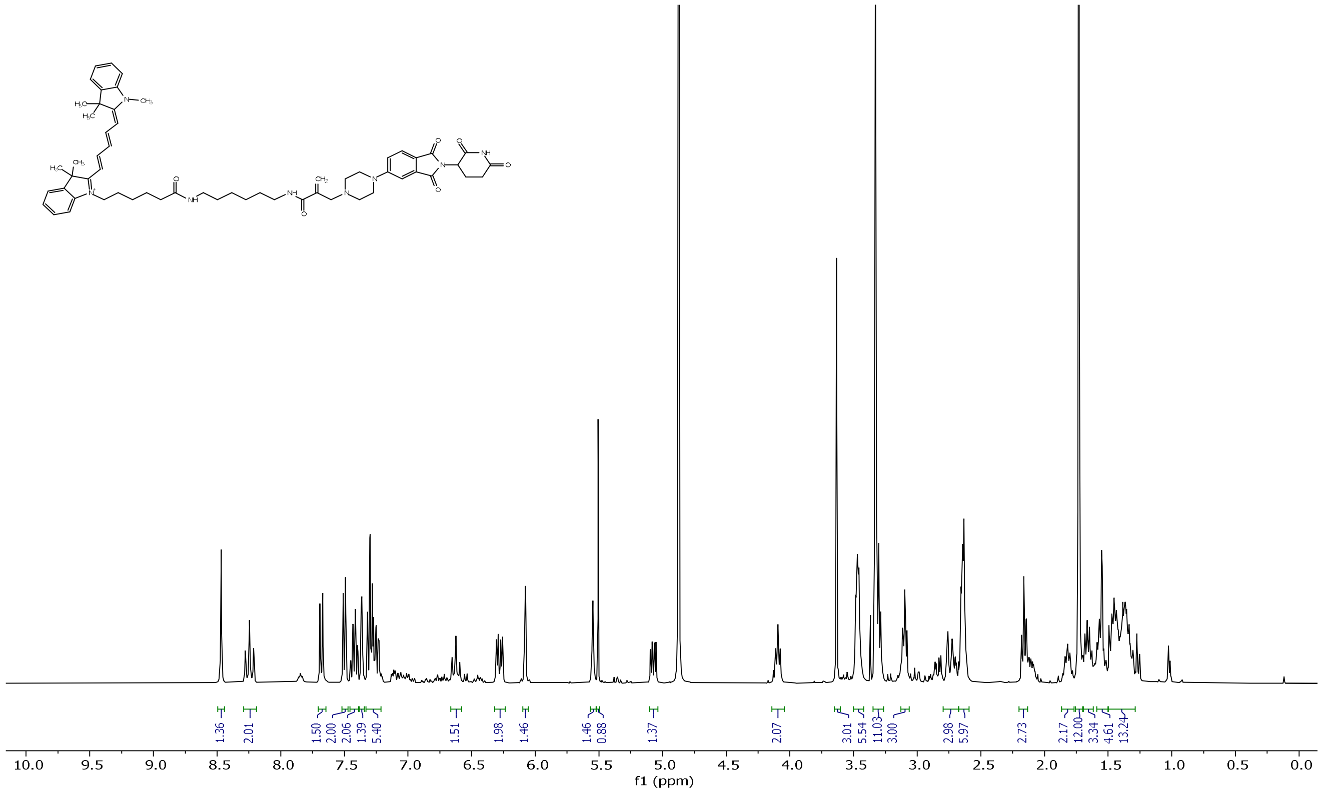
1.36
2.01
1.50
2.00
2.06
1.39
5.40
1.51
1.98
1.46
1.46
0.88
1.37
2.07
3.01
5.54
11.03
3.00
2.98
5.97
2.73
2.17
12.00
3.34
4.61
13.24
10.0 9.5 9.0 8.5 8.0 7.5 7.0 6.5 6.0 5.5 5.0 4.5 4.0 3.5 3.0 2.5 2.0 1.5 1.0 0.5 0.0
f1 (ppm)

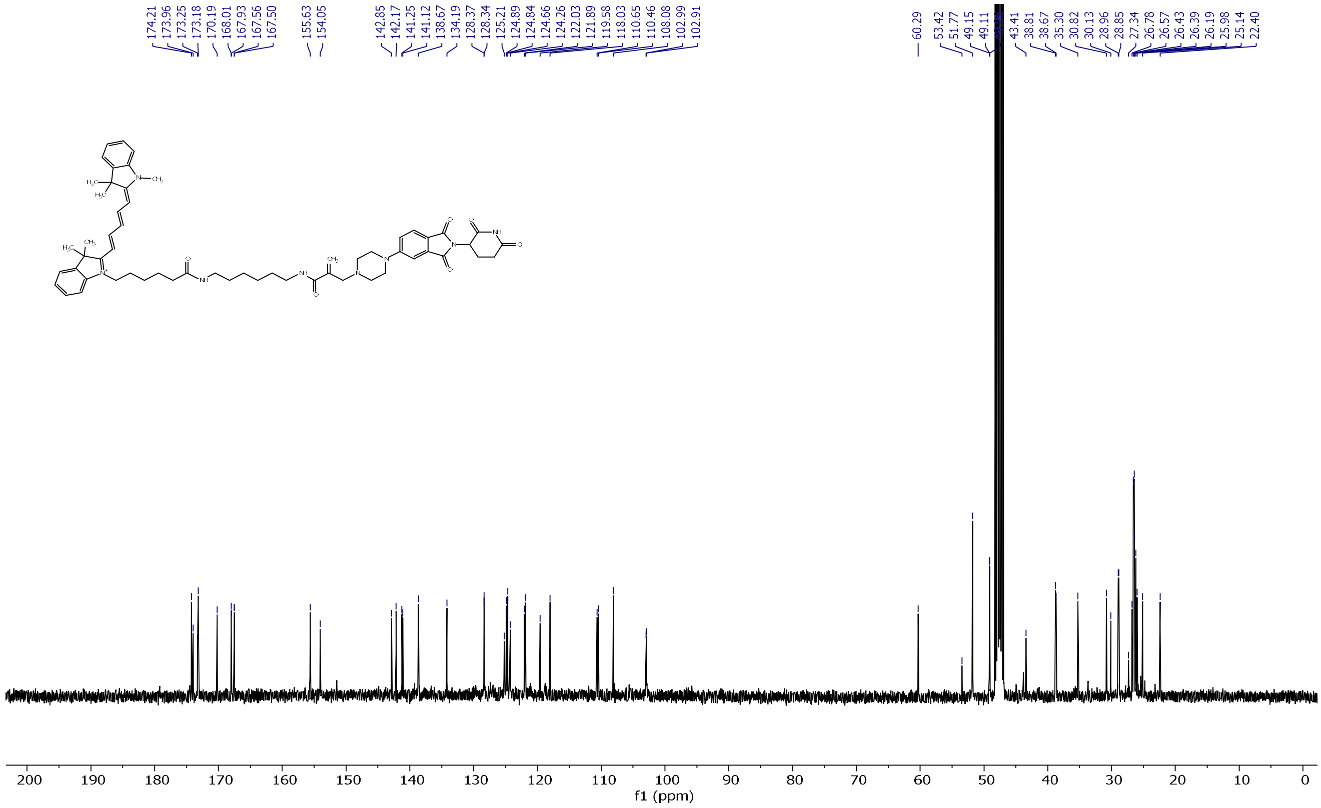
174.21
173.96
173.25
173.18
170.19
168.01
167.93
167.56
167.50
155.63
154.05
142.85
142.17
141.25
141.12
138.67
134.19
128.37
128.34
125.21
124.89
124.84
124.66
124.26
122.03
121.89
119.58
118.03
110.65
110.46
108.08
102.99
102.91
60.29
53.42
51.77
49.15
49.11
[illegible]
43.41
38.81
38.67
35.30
30.82
30.13
28.96
28.85
27.34
26.78
26.57
26.43
26.39
26.19
25.98
25.14
22.40
200 190 180 170 160 150 140 130 120 110 100 90 80 70 60 50 40 30 20 10 0
f1 (ppm)

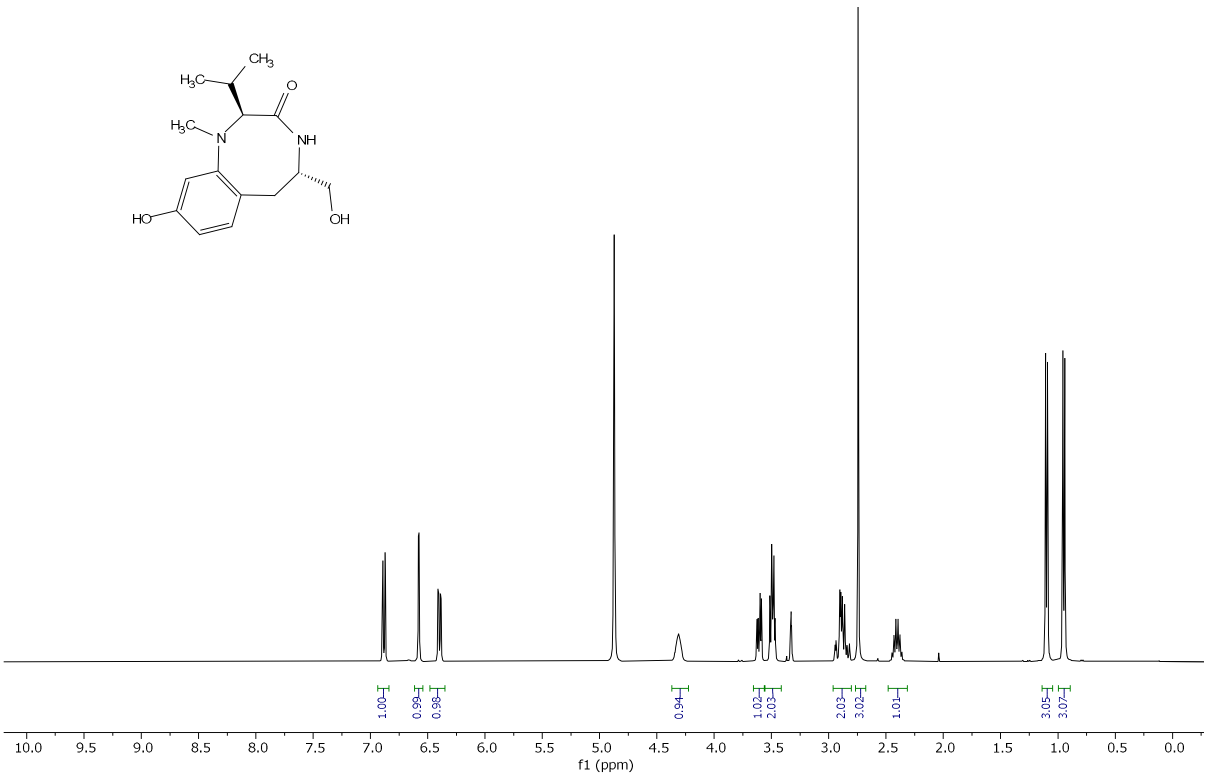

CH3
H3C
O
H3C
N
NH
HO
OH
1.00
0.99
0.98
0.94
1.02
2.03
2.03
3.02
1.01
3.05
3.07
10.0 9.5 9.0 8.5 8.0 7.5 7.0 6.5 6.0 5.5 5.0 4.5 4.0 3.5 3.0 2.5 2.0 1.5 1.0 0.5 0.0
f1 (ppm)

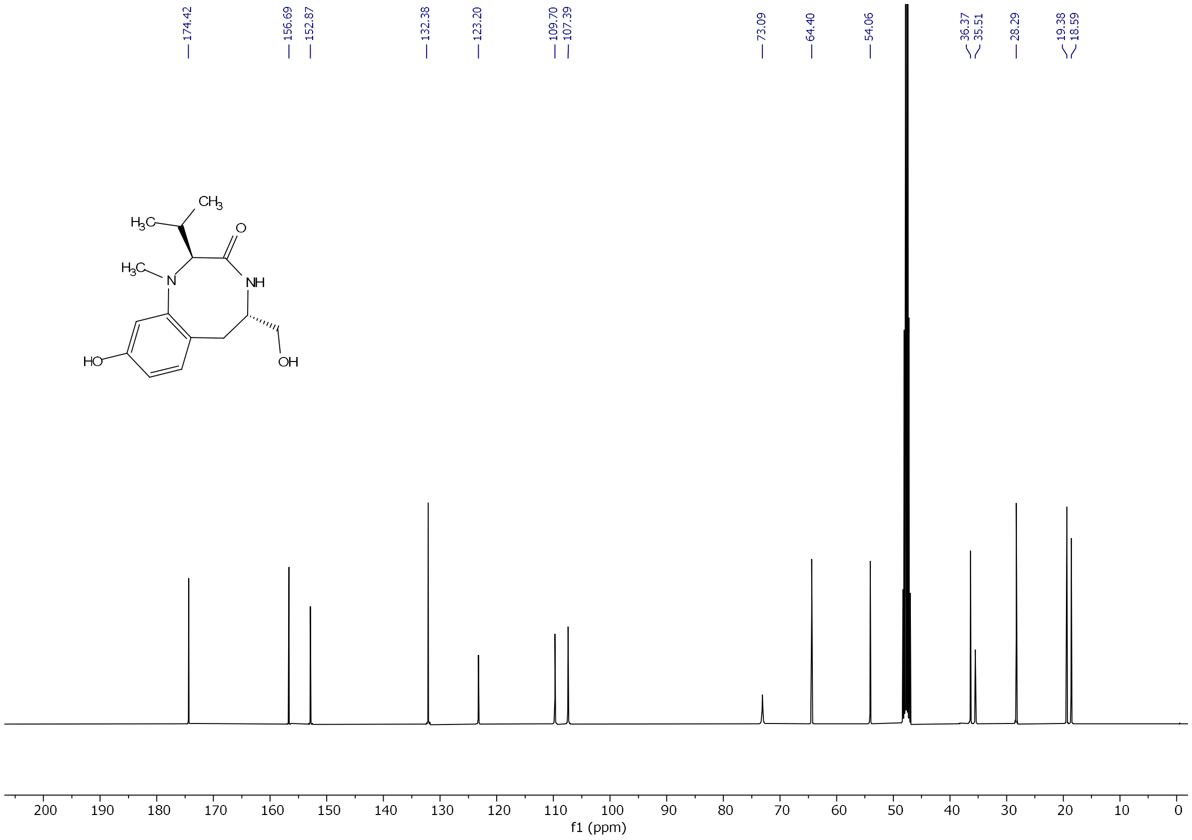

174.42
156.69
152.87
132.38
123.20
109.70
107.39
73.09
64.40
54.06
36.37
35.51
28.29
19.38
18.59
CH3
H3C
O
H3C
N
NH
HO
OH
200 190 180 170 160 150 140 130 120 110 100 90 80 70 60 50 40 30 20 10 0
f1 (ppm)

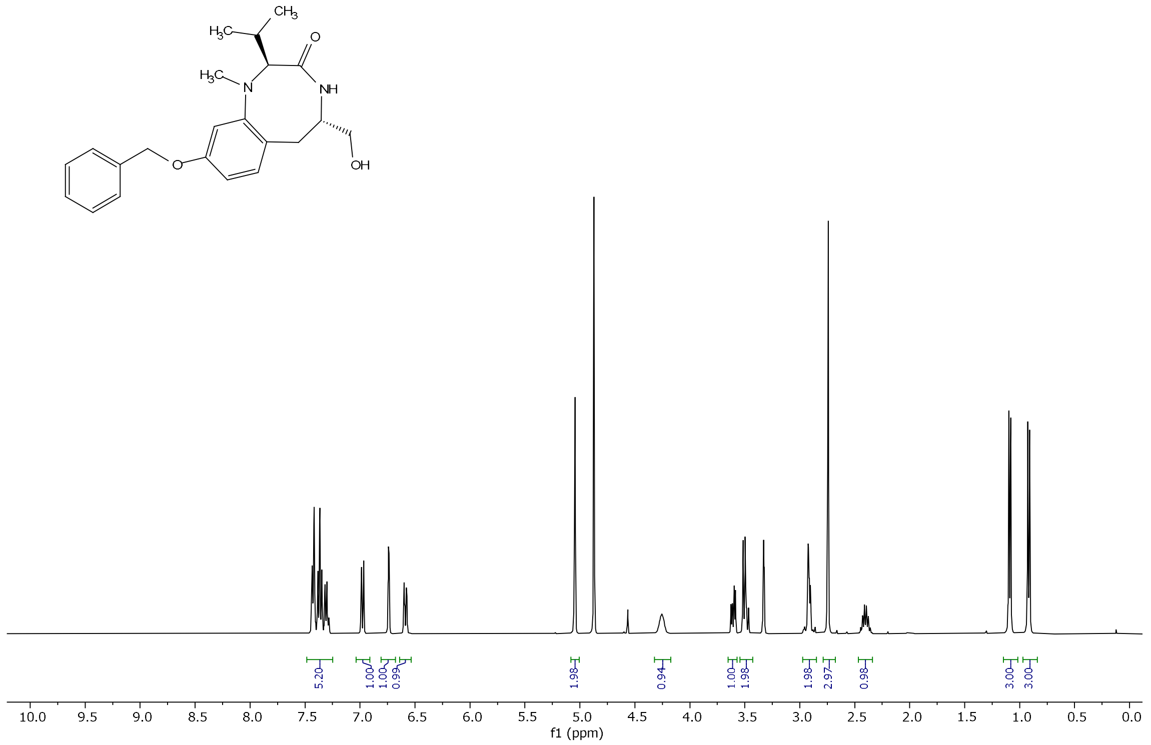

H3C
CH3
O
H3C
N
NH
O
OH
5.20
1.00
1.00
0.99
1.98
0.94
1.00
1.98
1.98
2.97
0.98
3.00
3.00
10.0
9.5
9.0
8.5
8.0
7.5
7.0
6.5
6.0
5.5
5.0
4.5
4.0
3.5
3.0
2.5
2.0
1.5
1.0
0.5
0.0
f1 (ppm)

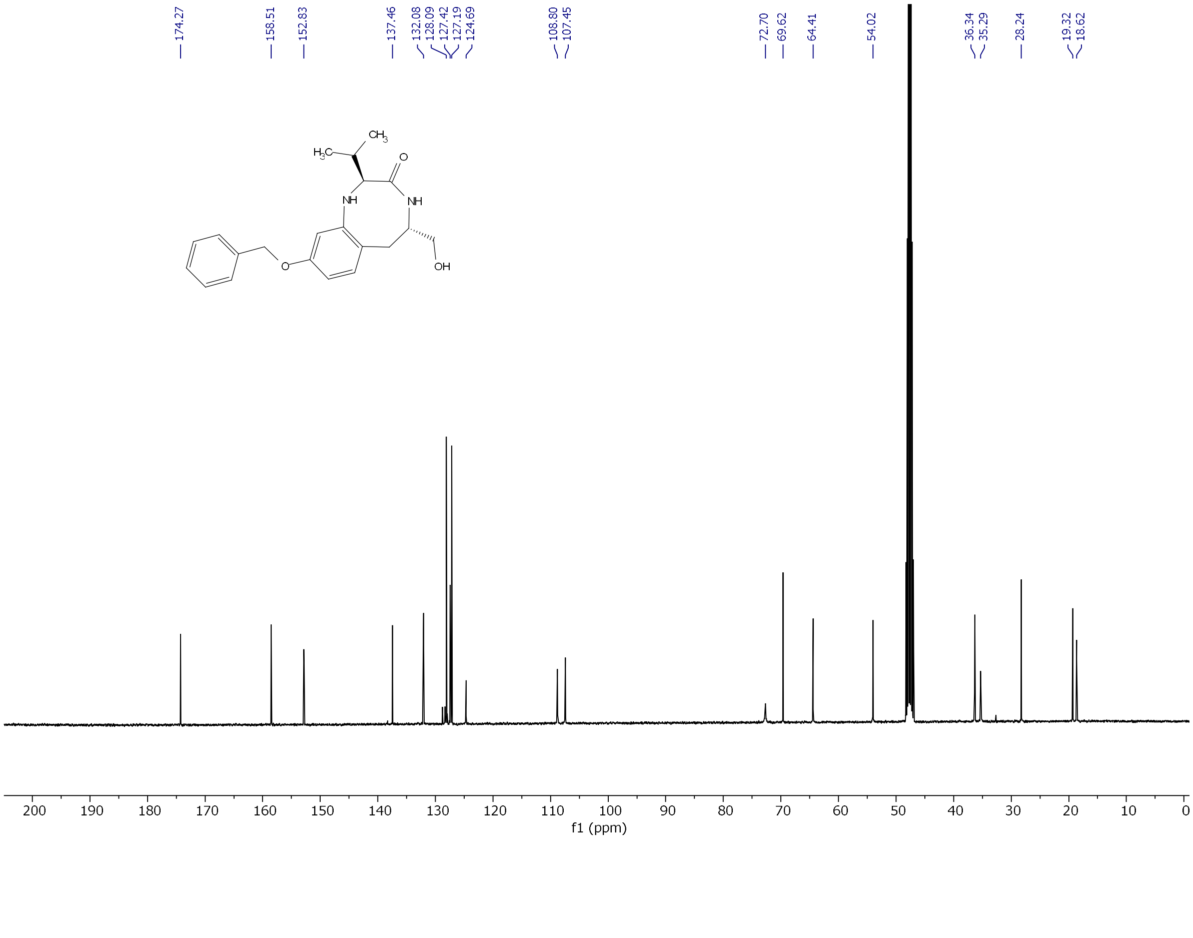

174.27
158.51
152.83
137.46
132.08
128.09
127.42
127.19
124.69
108.80
107.45
72.70
69.62
64.41
54.02
36.34
35.29
28.24
19.32
18.62
H3C
CH3
O
NH
NH
O
OH
200
190
180
170
160
150
140
130
120
110
100
90
80
70
60
50
40
30
20
10
0
f1 (ppm)

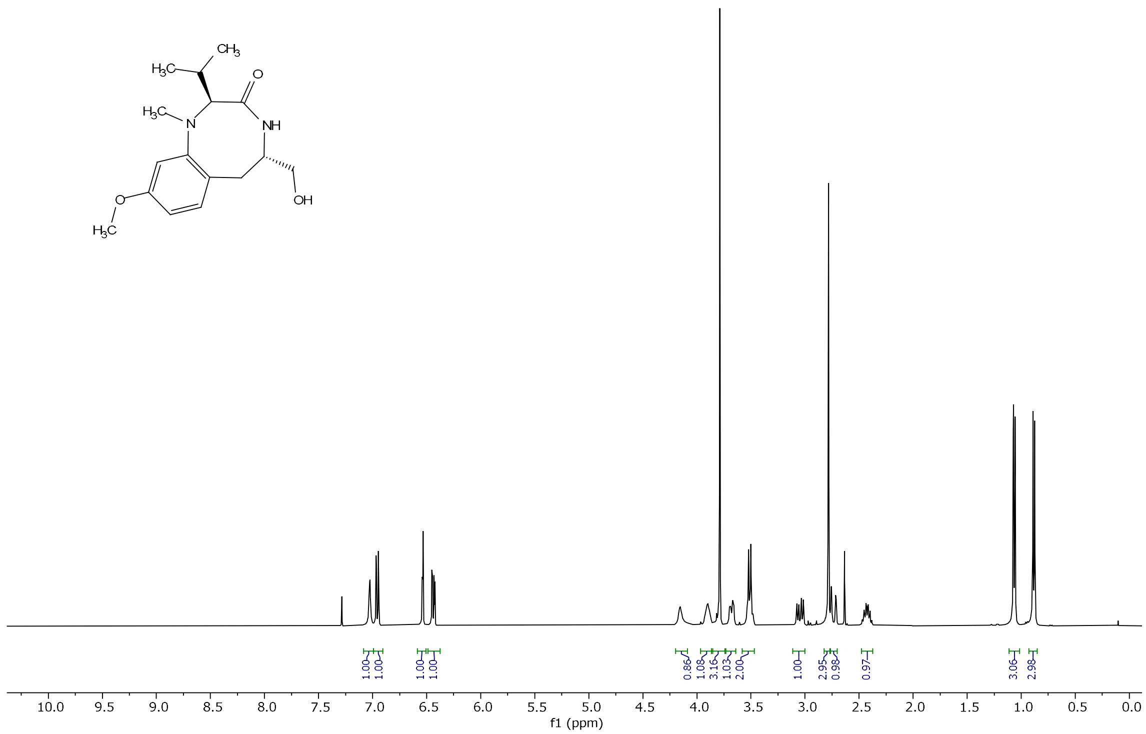
CH3
H3C
O
H3C
N
NH
O
OH
H3C
1.00
1.00
1.00
1.00
0.86
1.08
3.16
1.03
2.00
1.00
2.95
0.98
0.97
3.06
2.98
10.0 9.5 9.0 8.5 8.0 7.5 7.0 6.5 6.0 5.5 5.0 4.5 4.0 3.5 3.0 2.5 2.0 1.5 1.0 0.5 0.0
f1 (ppm)

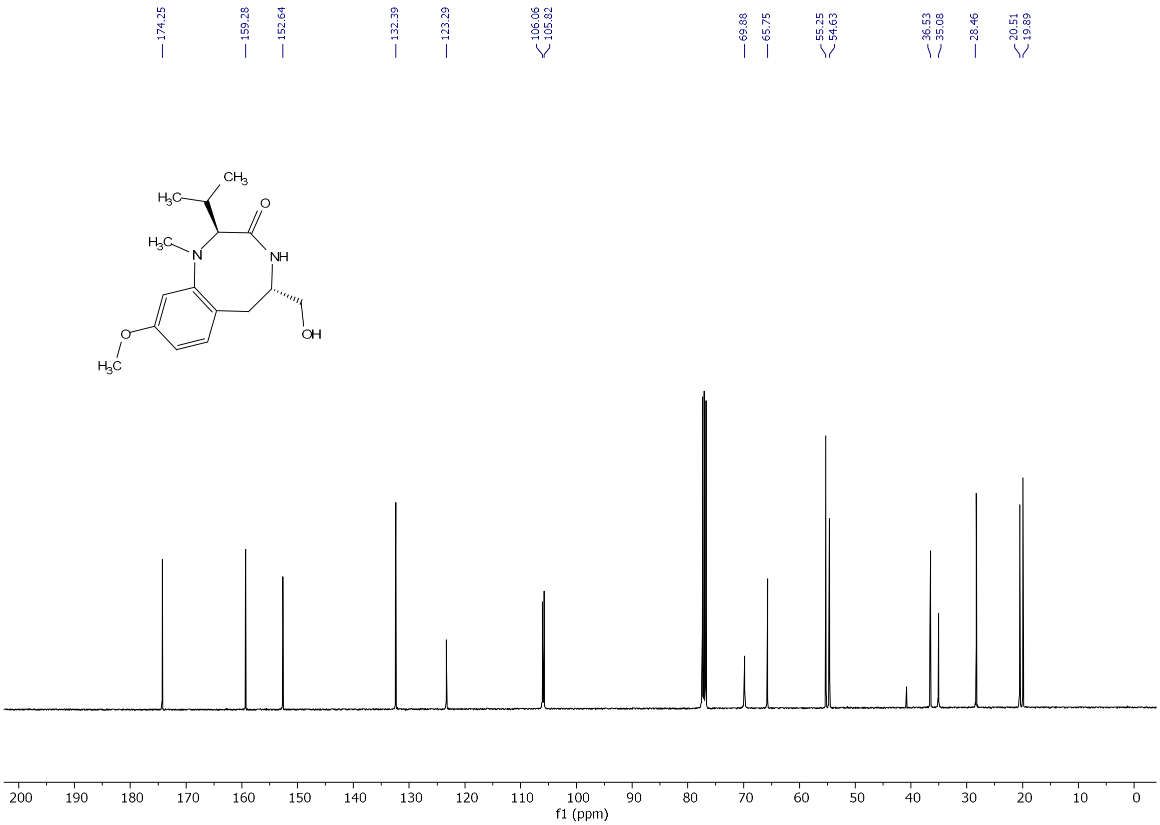
174.25
159.28
152.64
132.39
123.29
106.06
105.82
69.88
65.75
55.25
54.63
36.53
35.08
28.46
20.51
19.89
CH3
H3C
O
H3C
N
NH
O
OH
H3C
200 190 180 170 160 150 140 130 120 110 100 90 80 70 60 50 40 30 20 10 0
f1 (ppm)

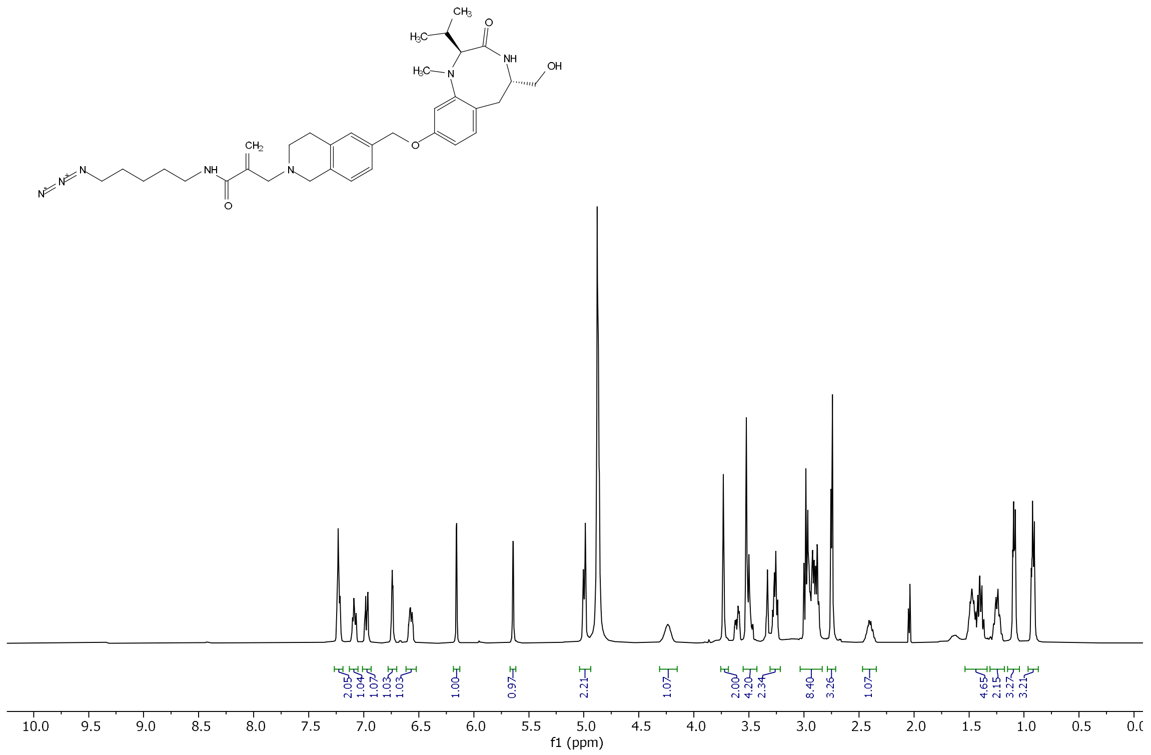
2.05
1.04
1.07
1.03
1.03
1.00
0.97
2.21
1.07
2.00
4.20
2.34
8.40
3.26
1.07
4.65
2.15
3.27
3.21
f1 (ppm)

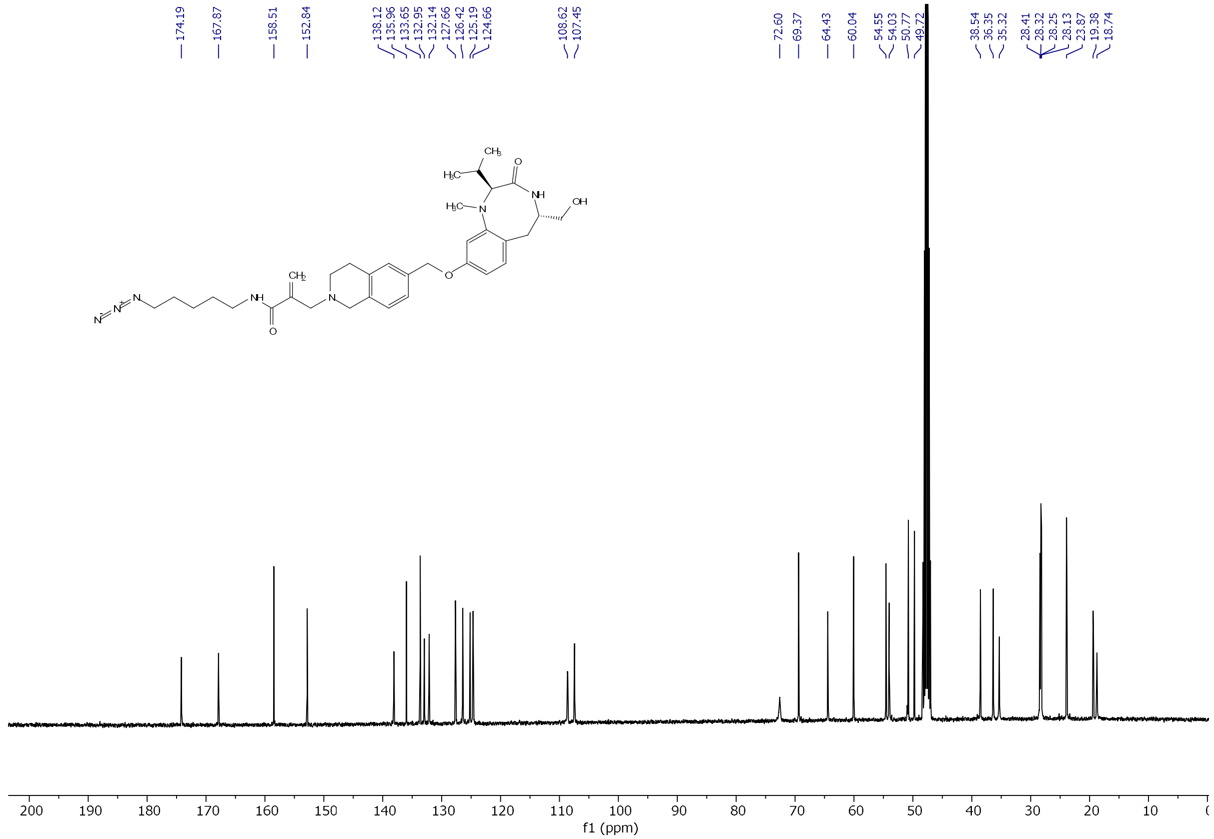
174.19
167.87
158.51
152.84
138.12
135.96
133.65
132.95
132.14
127.66
126.42
125.19
124.66
108.62
107.45
72.60
69.37
64.43
60.04
54.55
54.03
50.77
49.72
38.54
36.35
35.32
28.41
28.32
28.25
28.13
23.87
19.38
18.74
f1 (ppm)

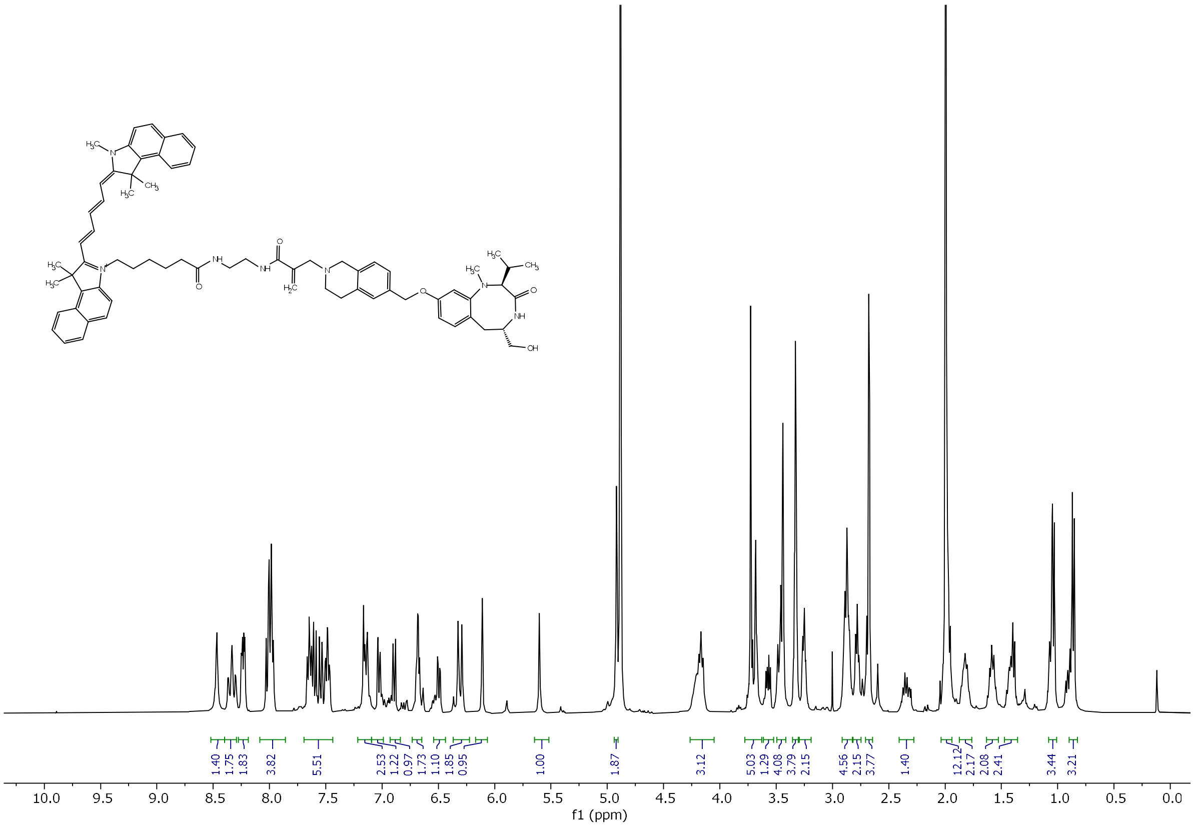
1.40
1.75
1.83
3.82
5.51
2.53
1.22
0.97
1.73
1.10
1.85
0.95
1.00
1.87
3.12
5.03
1.29
4.08
3.79
2.15
4.56
2.15
3.77
1.40
12.12
2.17
2.08
2.41
3.44
3.21
f1 (ppm)

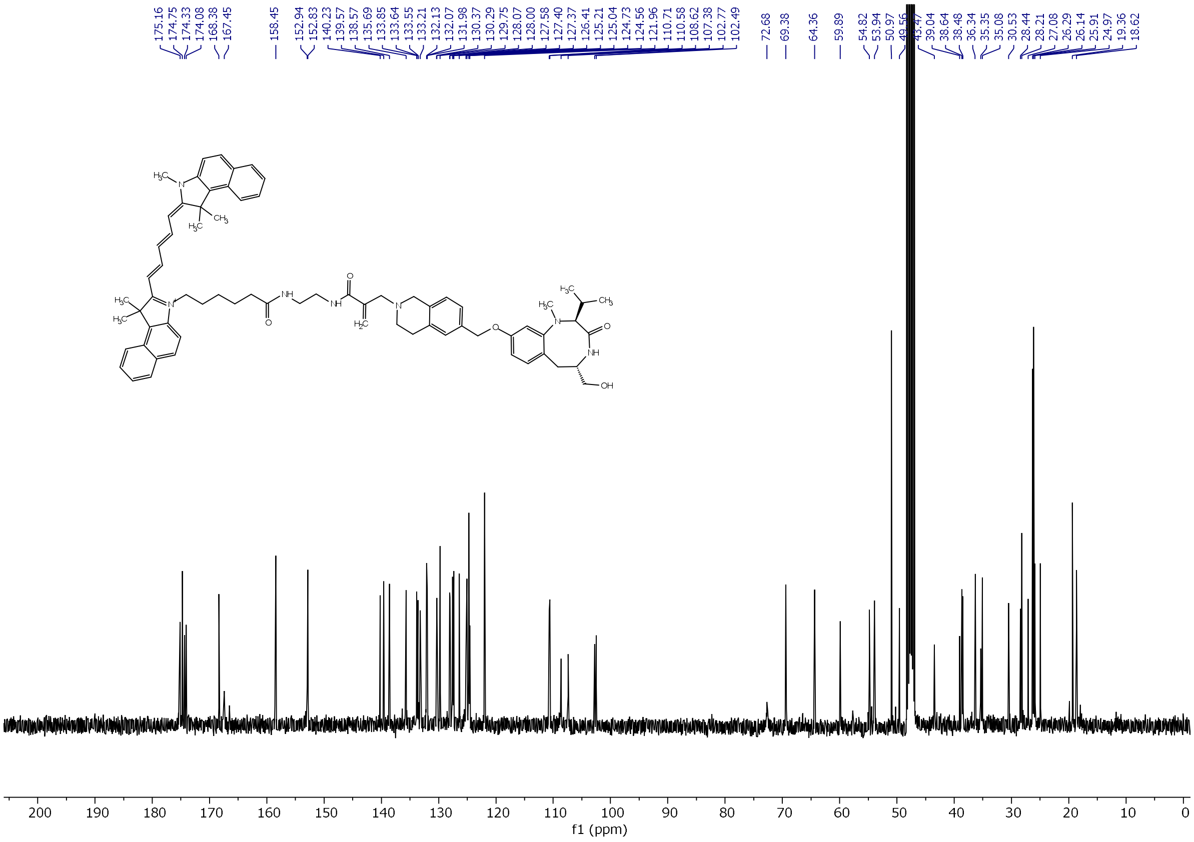
175.16
174.75
174.33
174.08
168.38
167.45
158.45
152.94
152.83
140.23
139.57
138.57
135.69
133.85
133.64
133.55
133.21
132.13
132.07
131.98
130.37
130.29
129.75
128.07
128.00
127.58
127.40
127.37
126.41
125.21
125.04
124.73
124.56
121.96
110.71
110.58
108.62
107.38
102.77
102.49
72.68
69.38
64.36
59.89
54.82
53.94
50.97
49.56
39.04
38.64
38.48
36.34
35.35
35.08
30.53
28.44
28.21
27.08
26.29
26.14
25.91
24.97
19.36
18.62
f1 (ppm)

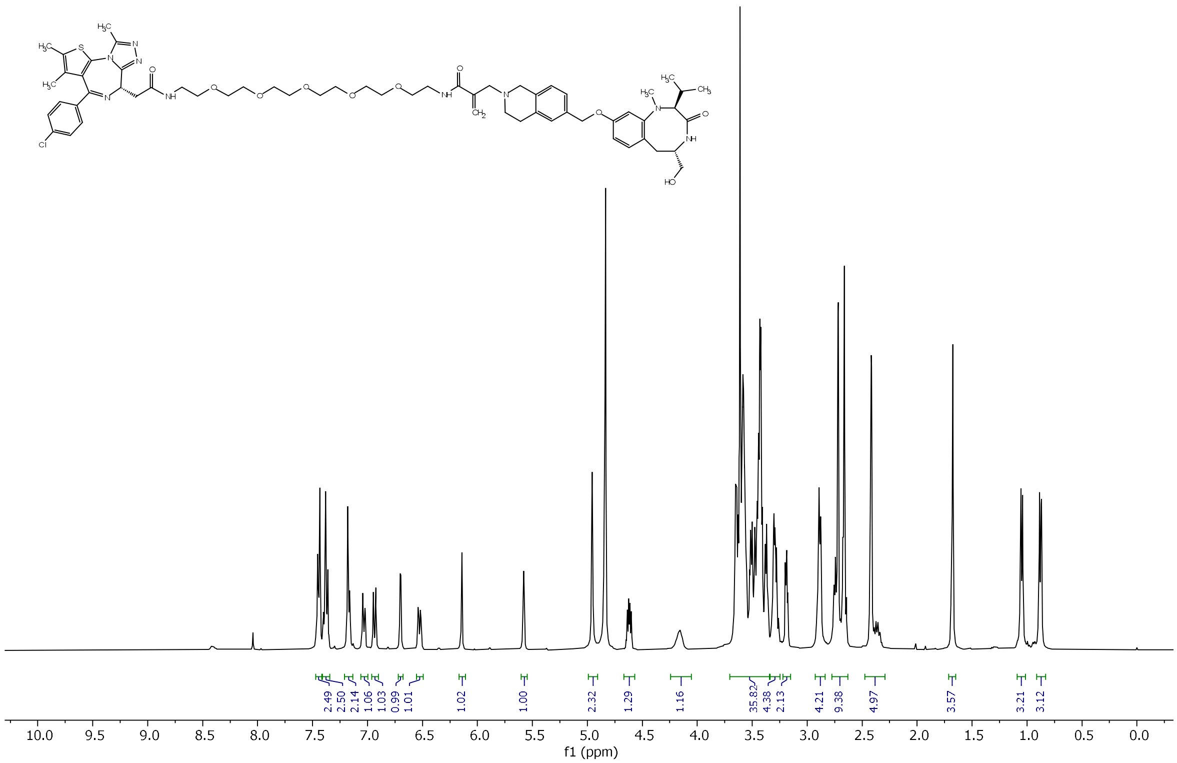

2.49
2.50
2.14
1.06
1.03
0.99
1.01
1.02
1.00
2.32
1.29
1.16
35.82
4.38
2.13
4.21
9.38
4.97
3.57
3.21
3.12
10.0 9.5 9.0 8.5 8.0 7.5 7.0 6.5 6.0 5.5 5.0 4.5 4.0 3.5 3.0 2.5 2.0 1.5 1.0 0.5 0.0
f1 (ppm)

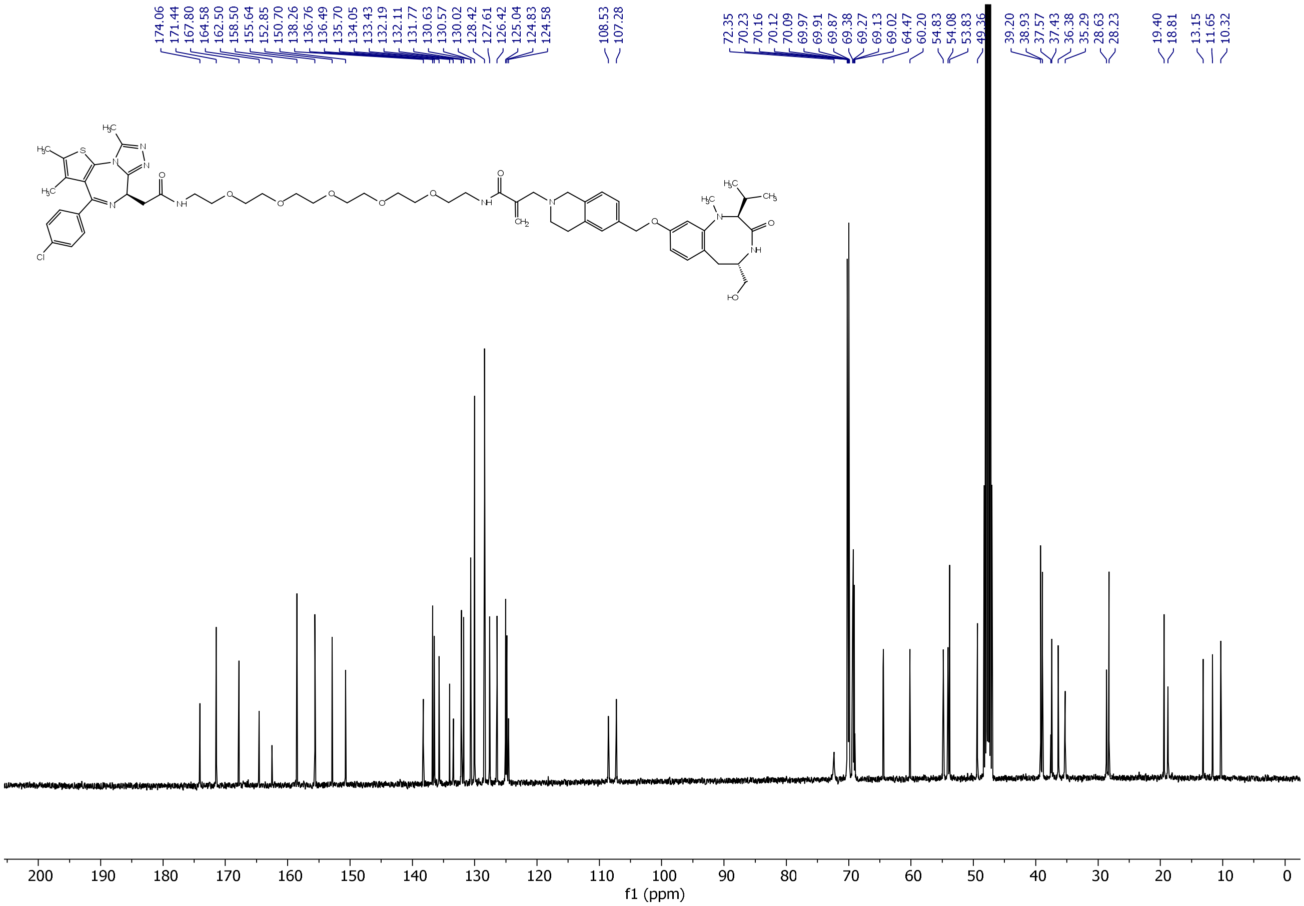

174.06
171.44
167.80
164.58
162.50
158.50
155.64
152.85
150.70
138.26
136.76
136.49
135.70
134.05
133.43
132.19
132.11
131.77
130.63
130.57
130.02
128.42
127.61
126.42
125.04
124.83
124.58
108.53
107.28
72.35
70.23
70.16
70.12
70.09
69.97
69.91
69.87
69.38
69.27
69.13
69.02
64.47
60.20
54.83
54.08
53.83
49.36
39.20
38.93
37.57
37.43
36.38
35.29
28.63
28.23
19.40
18.81
13.15
11.65
10.32
200 190 180 170 160 150 140 130 120 110 100 90 80 70 60 50 40 30 20 10 0
f1 (ppm)

1. Shoba, V.M. et al. Synthetic Reprogramming of Kinases Expands Cellular Activities of Proteins. *Angew Chem Int Ed Engl* **61**, e202202770 (2022).
